# Modeling Aerobic-Anaerobic Microbial Electrolysis Cells for Multi-step Methane Conversion

**DOI:** 10.64898/2026.11.18.767373

**Authors:** Spencer Gales, Ethan Howley, Reyes Sierra-Alvarez, Joel Cuello, Thomas K. Wood, Ingmar H. Riedel-Kruse

## Abstract

The bioconversion of methane by anaerobic methanotrophs has a higher carbon and energy efficiency compared to aerobic processes, suggesting significant potential for producing value-added chemicals and reducing greenhouse gas emissions. However, their practical use still faces many challenges regarding their culturing, genetic manipulation, and integration into multi-species consortia. Here we mathematically model a microbial electrolysis cell (MEC) to study the division of labor inside an anode-attached vertically stratified biofilm, where genetically modified *Methanosarcina acetivorans* and *Escherichia coli* perform the anaerobic oxidation of methane (AOM) to acetate followed by the aerobic conversion to 2,3-butanediol, respectively. We find that such a design is theoretically feasible, and we identify minimum biological improvements and operational conditions so that the aerobic layer effectively acts as oxygen-consuming living membrane protecting the anaerobic layer. Considering current biotechnological parameters points to a significant yet feasible need for further microbial engineering to increase the AOM rate. A comparison to a more traditional two-stage reactor at equivalent biomass reveals advantages and drawbacks of both designs, e.g., tradeoffs between peak product yield, reactor volume, and operational complexity. These results inform AOM driven microbial upgrading via aerobic-anaerobic stratified biofilms as well as value-added production via cooperative consortia in general.

## 1 Introduction

Methane as the main component in natural gas is an important energy source for modern industry and households, but also a major greenhouse gas [25]. Capturing methane released at localized sources, for conversion into a value-added product presents an opportunity to make industrial use of an abundant resource while minimizing the sites’ carbon footprints [25, 66]. A prime candidate for localized methane emissions is at wastewater treatment (WWT) plants where methane emissions consistently occur during multiple steps in the treatment process [3, 12]. Current technologies utilized to perform chemical conversion of methane related gases incur high capital and energy costs [25] rendering them unsuitable for localized source sites. Reactors utilizing biological catalysts, on the other hand, can provide lower cost and less energy intensive systems to achieve such tasks [15, 54]. However, biological oxidation of methane in industrial settings typically suffers from carbon inefficiency due to metabolic pathways resulting in carbon dioxide, slow growth times of bacterial methanotrophs, unculturable anaerobic methanotrophic archaea (ANME), low solubility of methane and challenges in bioreactor design and operation [54, 58]. Carbon and energy inefficiencies are more pronounced in aerobic bacterial methanotrophs compared to their archaeal counterparts, motivating efforts to harness archaeal hosts [25, 70]. Alternatively to native metabolic pathways in wild type cells, synthetic biology utilizes engineered metabolic pathways in culturable hosts for methane conversion to overcome highlighted limitations [6, 56, 70].

Leveraging synthetic biology combined with natural biofilm stratification, this work introduces a bioreactor design to expand the scope of methane oxidizing microbial consortia. In reactors, living aerobic biofilms can act as ‘shields’, using stratification to protect anaerobic biofilms [59, 69]. In WWT, while anodic biofilms often stratify naturally to shield the electroactive bacteria, cathodic stratification is typically an engineered design feature [59, 69]. For instance, in single chamber microbial fuel cells (SC-MFCs) stratified biofilms on an air cathode can be deliberately utilized for simultaneous nitrification and denitrification (SND) protecting the oxygen sensitive denitrifying bacteria and anode respiring bacteria. [59, 61–63]. While these setups focus on optimizing removal efficiency and energy recovery as their primary goal, there is a significant opportunity to actively leverage such stratification on a (porous) anode for targeted value-added product synthesis. To this end, we propose a microbial electrochemical cell (MxC) to actively exploit stratified microbial consortia via engineered metabolic pathways for methane oxidation and conversion (Fig. 1). MxCs include both exergonic MFCs and endergonic microbial electrolysis cells (MECs), the latter using an external voltage to drive hydrogen production. We adopt an MEC configuration in this study, as the applied voltage introduces an additional parameter for modulating the reaction rate [18, 24, 48, 49].We model a pseudo three chamber double-layer MEC, denoted DL-MEC, to turn methane to acetate to 2,3-butanediol (2,3-BDO). This proposed process uses engineered reverse methanogenesis in *Methanosarcina acetivorans* [29, 56] and aerobic product synthesis in the modified bacterium *Escherichia coli* [44, 51] (Fig. 1A).

**Figure 1.**
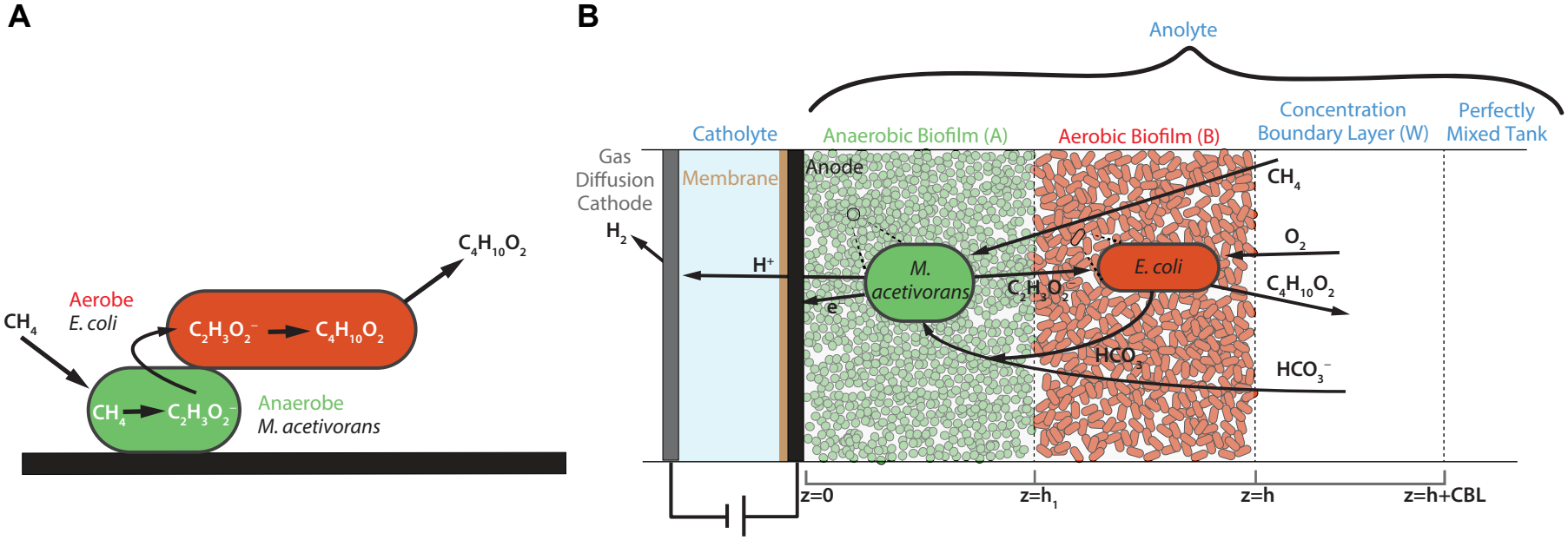
The pseudo three chamber double-layer microbial electrolysis cell (DL-MEC), enables the multi-step synthesis of value-added products coupled to anaerobic oxidation of methane (AOM). A.The motivating example of conversion from methane to 2,3-butanediol via intermediate acetate. The two sub-reactions take place in anaerobic and aerobic biofilm layers respectively. **B**. A detailed description of bioelectrochemical reactions. From the left to right: gas diffusion cathode where the electrons (e^−^) and protons (H^+^) come together to produce hydrogen, catholyte, proton exchange membrane (PEM) and bioanode, anaerobic biofilm (A), aerobic biofilm (B), concentration boundary layer (W) and, the continuously stirred tank. At = 0 a porous conductive surface is assumed, at *z* = *h*_1_, the biofilm-biofilm interface, at *z* = *h* = *h*_1_ + *h*_2_ is the biofilm-liquid interface, at *z* = *h* + *L*_*L*_ a well-mixed large liquid volume which serves as a reservoir for reactants and products excluding hydrogen.

Given the complexity of engineering such a system, multiscale modeling is a cost effective first approach to identifying potential opportunities and challenges for reactor design, and for providing engineers and microbiologists with microbial engineering targets. This paper performs the corresponding evaluation by using a targeted global sensitivity analysis to determine the minimum biological values required to sustain a physically feasible biofilm thickness with distinctly layered environments (Fig. 1**B**). Specifically sensitivity analysis is performed over a developed 1D stationary biofilm model embedded in a perfectly mixed continuously stirred anolyte in the electric potential non-limiting regime. This approach simultaneously estimates the maximum theoretical product yield at these values. While the DL-MEC is specific to 2,3-BDO production, it serves as a template for aerobic-anaerobic biofilm conversion to high value products in general, as acetate is a common precursor for other products [22, 32].

## 2 Background

The proposed DL-MEC relies on two synthetically engineered microbes to achieve multi-step methane to product conversion. We consider engineered archaeal methanogen Methanosarcina acetivorans, for AOM via reverse methanogenesis. Enzyme methyl-coenzyme M reductase (MCR) was cloned from an unculturable ANME taken from a Black Sea mat into the archaeal methanogen *M. acetivorans*. As a result, synthesis of acetate from methane, with ferric iron particulates serving as the terminal electron acceptor (TEA) was achieved with a concentration of 10.3 mM over five days. Then, acetate production was increased to 5.4 *µ*mol acetate per mg protein over six days [29]. Other candidate ANME have been considered for anaerobic conversion of methane to acetate, including a microbial culture dominated by *Methanoperedens nitroreducens* [6]. Although progress has been made [40], *M. acetivorans* cannot produce a wide array of complex products, therefore a second species of engineered microbes is necessary.

The facultative anaerobe engineered bacteria *E. coli* was shown to aerobically convert acetate to 2,3-BDO [44, 51]. *E. coli* has been further engineered to increase the 2,3-BDO yield and uptake consumption rate of acetate, the sole carbon source, to 0.78 g acetate (g cell dry weight (CDW))^−1^ day ^−1^ [51]. While product inhibition and backwards conversion of 2,3-BDO has been observed, this occurs at concentrations that exceed this papers considered values. Hence, complete conversion from the precursor acetoin to 2,3-BDO is assumed [36]. The 2,3-BDO production from acetate was shown to be positively associated with oxygen concentration and biomass growth *E. coli* [8]. These engineered *E. coli* were studied while microbes were in planktonic form in a fully aerobic environment. However, *M. acetivorans* requires a strictly anaerobic environment, necessitating aerobic *E. coli* to sufficiently reduce oxygen before it reaches the anaerobic biofilm.

In order to harness microbial consortia consisting of aerobes and anaerobes, a physical or biological mechanism is required to enforce oxidative conditions in both bioreactors and MxCs. Two standard setups are typically employed: single chamber MxCs utilizing biological shielding/stratification usually occurring on an (air) cathode [11, 19, 23, 33, 35, 59, 69] and dual chamber MxCs using a physical mechanism such as a membrane [2, 4]. SC-MxCs tend to be more cost efficient relying on biological shielding and air cathode aeration rather than membranes and active aeration control. Despite these advantages, SC-MxCs are limited by oxygen inhibition/toxicity [59]. Excessive oxygen interferes with several microbial and electrochemical processes: (i) inhibiting the metabolism of anaerobes, (ii) diverting electrons to oxygen as the terminal electron acceptor (TEA), (iii) stealing electrons directly from the anode. These can cause dire effects decreasing the Coulombic efficiency and effectively short circuiting the ‘circuit’ [1, 13, 69].

In the DL-MEC, similar to stratified SC-MFCs, we use a ‘pseudo/living membrane’ to reduce oxygen and limit diffusion resulting in an aerobic anolyte and anaerobic anodic chamber. Multi-species biofilms naturally stratify in response to oxygen concentrations, with outer aerobic regions aiming to reduce oxygen that would damage the inner regions anaerobic microbes [69]. To date, only solid phase electron acceptors have been shown to work with reverse methanogenesis in *M. acetivorans* [34]. Consequently, the aerobic biofilm limits insoluble particulate diffusion to the anaerobes. Instead, a nondepletable conductive anode is used as the TEA. Between the porous bioanode and cathode a proton exchange membrane (PEM) only allows protons into the catholyte towards a gas diffusion cathode (GDC) where hydrogen is recovered (Fig. 1) [57]. By substituting a particulate TEA with a porous anode to produce hydrogen gas, the consideration of microbial electrokinetic mechanisms for electron transfer becomes salient [5]. In a synthetic microbial consortium consisting of *M. acetivorans, Geobacter sulfurreducens* and microbes from methane-acclimated sludge, electricity was generated via AOM to acetate with humic acids as the TEA [39]. Diffusible redox mediators also support long distance extracellular electron transfer (EET) without a co-culture, for example, methylene blue facilitated EET with engineered *E. coli* for acetoin production [16]. By the assumption of non-limiting electron transfer to the anode, the electron transfer mechanism is discussed but not incorporated into the present model and is deferred to further work.

The 1D DL-MEC biofilm, assumed in stationary phase, is operated in continuous cultivation, though batch dynamics motivate reactor design choices. We assume no direct modeling of growth, planktonic attachment/detachment or pH. Global sensitivity analysis over reaction rates and biofilm densities identifies feasible layer thicknesses for product synthesis, providing engineers and biologists guidance on preferential operating conditions. The findings of this study must be contextualized by the model’s inherent limitations (i) While growth is not modeled, the stratified biofilm represents a state with ongoing growth. The *M. acetivorans* biofilm is anchored against the fixed, nondepletable porous anode, thus any growth is expected to displace the anaerobic-aerobic interface upward, while at the liquid-biofilm boundary excess biomass of *E. coli*, being fast growing with growth associated production, would accumulate preferentially at the shear-exposed surface constrained to 100–150 *µ*m [42]. A steady stratification between the layers is further supported by the low growth yield of methanogens forward methanogenic pathway [52] as they are not expected to competitively displace *E. coli* biofilm. However, whether such a steady state interface exists or *E. coli* grows downwards displacing the methanogens, is assumed in this paper, and is left to future work. (ii) A non-limiting electric potential regime is assumed. While electrochemical processes, such as electron transfer mechanisms and electric potential, influence effective reaction rates, they cannot exceed the cell’s metabolic maximum reaction rates. The present results, therefore, provide lower bounds on the required reaction rates. (iii) The analysis focuses on catabolic reactions favorable for product synthesis, overestimating available resources. In practice, endogenous anabolic metabolism reduces the energy and nutrients available for product synthesis and inert biomass reduces the total active biomass reflecting a snapshot of the biofilm rather than a time dependent trajectory. Reaction rates are centered on acetate production/consumption, with 2,3-BDO synthesis as a production upper bound defined by the theoretical yield enabling a direct assessment of DL-MEC feasibility. Significant computational bioelectrochemical work has already been performed mainly with regards to a bioanode and to a lesser extent for a biocathode [17, 18, 24, 30, 46–48]. These works represent modeling of important electrochemical features, however, such electrochemical phenomena have not yet been incorporated into the DL-MEC model and constitutes a critical direction for future development.

## 3 Multiscale Model

We introduce the multiscale reactor, denoted DL-MEC, addressing the biochemical, mathematical and electrochemical components central to designing such a system. The DL-MEC is a continuously stirred pseudo three chamber cell where the anolyte compartment is on the order of cm to meters while the thickness of the biofilms is on the order of *µ*m.

### 3.1 Reactor and Biochemical Model: Anaerobic Oxidation of Methane and 2,3 -Butanediol Synthesis

We introduce the DL-MEC layout and the chemical equations that govern the redox reactions of particular interest in each layer (Fig. 1). From left to right, the hydrogen gas collection chamber, gas diffusion cathode (GDC), catholyte and porous bioanode that govern hydrogen electrolysis setup of the DL-MEC. Attached to the porous bioanode sit the stratified biofilms. In the biofilm the net reactions happen in sequence, first AOM occurs in the anaerobic *M. acetivorans* biofilm, denoted A, then 2,3-BDO production occurs in the aerobic *E. coli* biofilm, denoted B. On the other side of the biofilm sits the concentration boundary layer (CBL) and perfectly mixed tank which serves as a reservoir for the initial reactants and final product aside from hydrogen gas. The detailed governing chemical equations are described in the remainder of the section.

#### 3.1.1 Anaerobic Reaction in the *M. acetivorans* Biofilm

Engineered *M. acetivorans* was established to convert CH_4_ to C_2_ H_3_ O_2_ ^−^ via reduction of iron Fe^3+^ [56]. We emphasize that this reaction is energetically favorable with Gibbs free energy 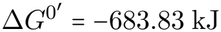. For our proposed design, we cannot efficiently feed Fe^3+^ to the anaerobic biofilm because it is fixed between the aerobic biofilm and the substratum, instead we utilize a porous bioanode as a nondepletable TEA (Fig. 1**B**) via an electrolysis cell leading to the production of hydrogen gas. The reactions inside the anaerobic biofilm at the anode:

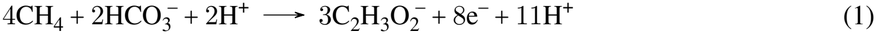

With 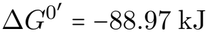. Reaction at the GDC:

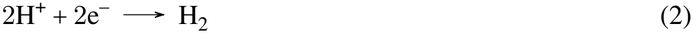

with 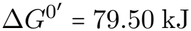 Leading to an overall reaction:

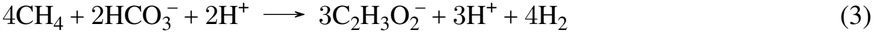

with a resultant 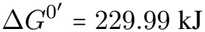 or 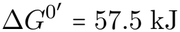 (mol of methane) ^−1^. Using 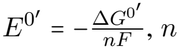 is the number of electrons transferred and *F* is Faraday’s constant, the standard reduction/cathode potential of (2) is 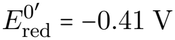 and the standard oxidation potential (1) is 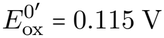. Resulting in the open circuit cell potential or counter electromotive force, 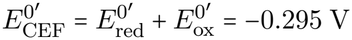. This reaction is not energetically favorable exemplifying the need for an energy input of at least 0.295 V with the assumption of equimolar concentrations and not accounting for overpotential energy requirements.

#### 3.1.2 Aerobic Reaction in the *E. coli* Biofilm

For production of 2,3-BDO occuring in Layer B, acetate is the sole carbon source and the electron donor

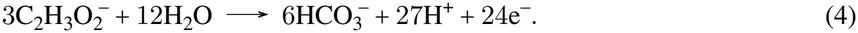

Oxygen is the electron acceptor for respiration

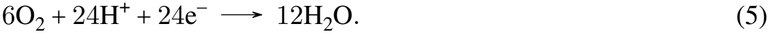

Acetate is the electron acceptor for conversion to 2,3-BDO [44]

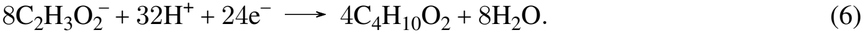

While the fraction of electron equivalents oxidized from acetate that reduce oxygen changes depending on several factors including the *E. coli* microbes oxygen availability and metabolic priorities [8] we assume the stoichiometry in following chemical reaction [8]

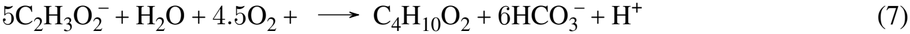

Resulting in 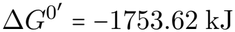 or 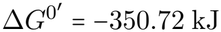 (mol of acetate) ^−1^.

### 3.2 Mathematical Model

We present the mathematical equations by compartment, starting with the material balance continuously stirred bulk anolyte then the reaction diffusion equations in the biofilm.

#### 3.2.1 Material Balance in the Bulk Anolyte

In the bulk anolyte we assume that no reactions take place. For the concentrations of chemical species, let *S*_1_ = [O_2_], *S*_2_ = [CH_4_], *S*_3_ = [C_2_ H_3_ O_2_ ^−^], *S*_4_ = [C_4_H_10_O_2_], *S*_5_ = [HCO_3_ ^−^], *S*_*H*2_ = [H_2_] where [S] is the concentration of chemical species S. Refer to Table 2 and 3 for all variable and parameter values. The first set of equations is for the continuously stirred tank reactor of water volume *V*_*W*_. Then for *i* ∈ {2, … , 5}

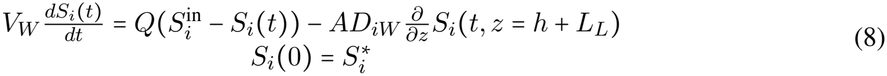

where volumetric flow rate *Q* is 0 for batch and positive for a continuous cultivation, *D*_*iW*_ is the diffusion coefficient of the ith species in water, *A* is the area of the biofilm and 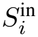 is the influent chemical species *S*_*i*_. At the onset of the cultivation there is no influent oxygen. After some time *τ* > 0, oxygen is delivered at delivery rate *η* and settles to a steady state where the oxygen bulk concentration is kept at a constant value, governed by the following delay onset differential equation

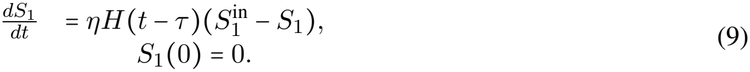

where *H* is the heaviside function. This delayed delivery allow acetate to accumulate in the aerobic biofilm before respiration can occur, diminishing the amount of oxygen entering the anaerobic biofilm as will be explored later.

**Table 1:**
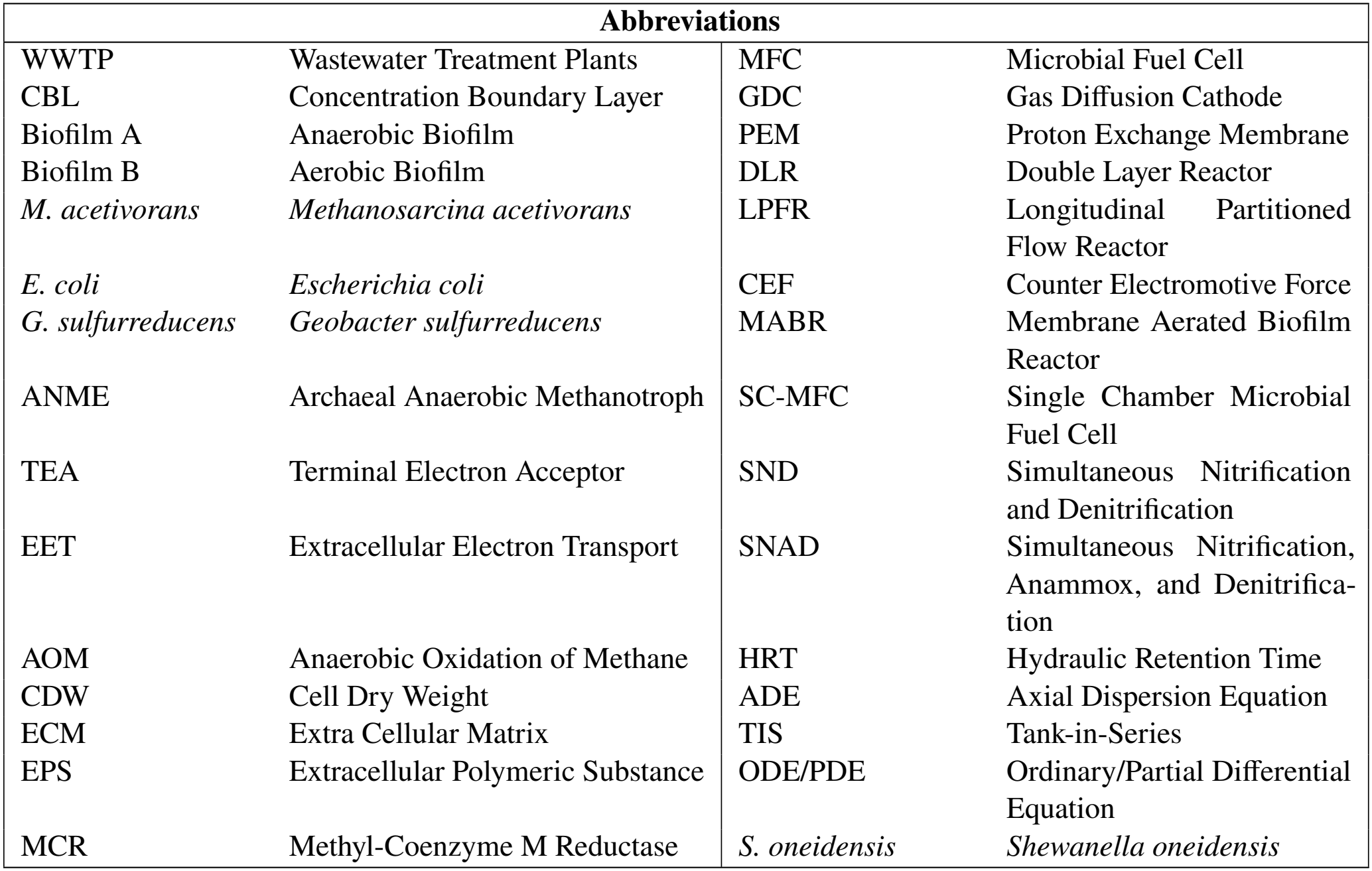
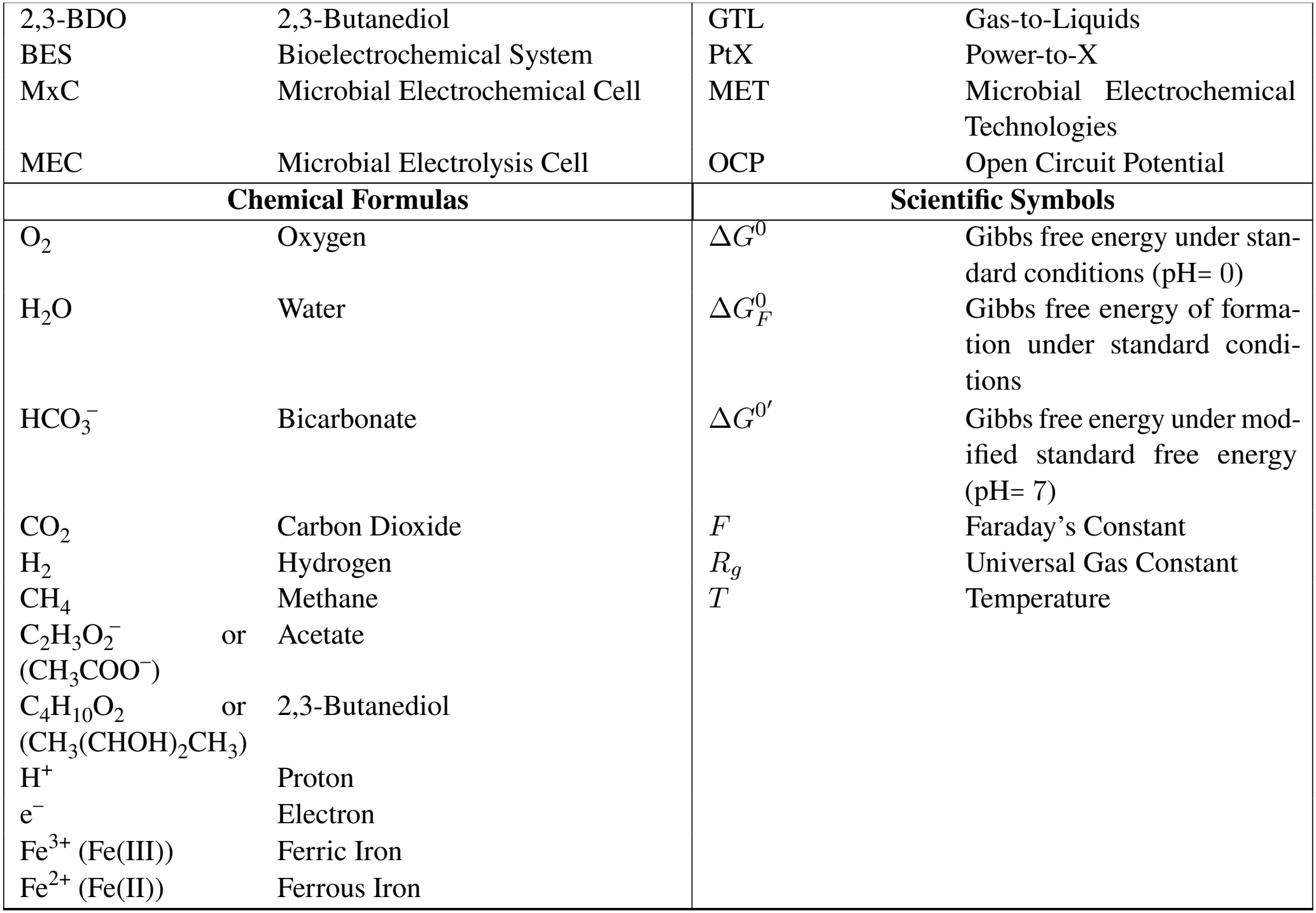
Nomenclature: Abbreviations, chemical formulas, and scientific symbols.

| **Abbreviations** |  |  |  |
|---|---|---|---|
| WWTP | Wastewater Treatment Plants | MFC | Microbial Fuel Cell |
| CBL | Concentration Boundary Layer | GDC | Gas Diffusion Cathode |
| Biofilm A | Anaerobic Biofilm | PEM | Proton Exchange Membrane |
| Biofilm B | Aerobic Biofilm | DLR | Double Layer Reactor |
| *M. acetivorans* | *Methanosarcina acetivorans* | LPFR | Longitudinal Partitioned Flow Reactor |
| *E. coli* | *Escherichia coli* | CEF | Counter Electromotive Force |
| *G. sulfurreducens* | *Geobacter sulfurreducens* | MABR | Membrane Aerated Biofilm Reactor |
| ANME | Archaeal Anaerobic Methanotroph | SC-MFC | Single Chamber Microbial Fuel Cell |
| TEA | Terminal Electron Acceptor | SND | Simultaneous Nitrification and Denitrification |
| EET | Extracellular Electron Transport | SNAD | Simultaneous Nitrification, Anammox, and Denitrifica-tion |
| AOM | Anaerobic Oxidation of Methane | HRT | Hydraulic Retention Time |
| CDW | Cell Dry Weight | ADE | Axial Dispersion Equation |
| ECM | Extra Cellular Matrix | TIS | Tank-in-Series |
| EPS | Extracellular Polymeric Substance | ODE/PDE | Ordinary/Partial Differential Equation |
| MCR | Methyl-Coenzyme M Reductase | *S. oneidensis* | *Shewanella oneidensis* |
| 2,3-BDO | 2,3-Butanediol | GTL | Gas-to-Liquids |
| BES | Bioelectrochemical System | PtX | Power-to-X |
| MxC | Microbial Electrochemical Cell | MET | Microbial Electrochemical Technologies |
| MEC | Microbial Electrolysis Cell | OCP | Open Circuit Potential |
| **Chemical Formulas** |  | **Scientific Symbols** |  |
| O<sub>2</sub> | Oxygen | Δ*G*<sup>0</sup> | Gibbs free energy under stan-dard conditions (pH= 0) |
| H<sub>2</sub>O | Water | Δ*G*<sup>0</sup><sub>*F*</sub> | Gibbs free energy of forma-tion under standard condi-tions |
| HCO<sub>3</sub><sup>−</sup> | Bicarbonate | Δ*G*<sup>0′</sup> | Gibbs free energy under mod-ified standard free energy (pH= 7) |
| CO<sub>2</sub> | Carbon Dioxide | *F* | Faraday's Constant |
| H<sub>2</sub> | Hydrogen | *R*<sub>*g*</sub> | Universal Gas Constant |
| CH<sub>4</sub> | Methane | *T* | Temperature |
| C<sub>2</sub>H<sub>3</sub>O<sub>2</sub><sup>−</sup> or (CH<sub>3</sub>COO<sup>−</sup>) | Acetate |  |  |
| C<sub>4</sub>H<sub>10</sub>O<sub>2</sub> or (CH<sub>3</sub>(CHOH)<sub>2</sub>CH<sub>3</sub>) | 2,3-Butanediol |  |  |
| H<sup>+</sup> | Proton |  |  |
| e<sup>−</sup> | Electron |  |  |
| Fe<sup>3+</sup> (Fe(III)) | Ferric Iron |  |  |
| Fe<sup>2+</sup> (Fe(II)) | Ferrous Iron |  |  |

**Table 2:** Stoichiometric Matrix (Peterson Matrix). We compute the reaction rate of component *S*_*k*_ as 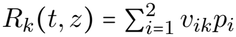, and the reaction rate of 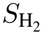 as 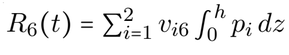.

| **Process** *i* | **Component** *k* |  |  |  |  |  | **Process Rate (***p*<sub>*i*</sub>**)** |
|---|---|---|---|---|---|---|---|
|  | *S*<sub>1</sub> | *S*<sub>2</sub> | *S*<sub>3</sub> | *S*<sub>4</sub> | *S*<sub>5</sub> | *S*<sub>H<sub>2</sub></sub> |  |
| Utilization & Production in Anaerobic Biofilm | 0 | −4 | 3 | 0 | −2 | 4 | $\mu_A \frac{K_{1A}}{K_{1A}+S_1} \cdot \frac{S_2}{K_{2A}+S_2} \cdot \frac{S_5}{K_{5A}+S_5} \cdot X_A$ |
| Utilization & Production in Aerobic Biofilm | −4.5 | 0 | −5 | 1 | 6 | 0 | $\mu_B \frac{S_3}{K_{3B}+S_3} \cdot \frac{S_1}{K_{1B}+S_1} \cdot X_B$ |

**Table 3:**
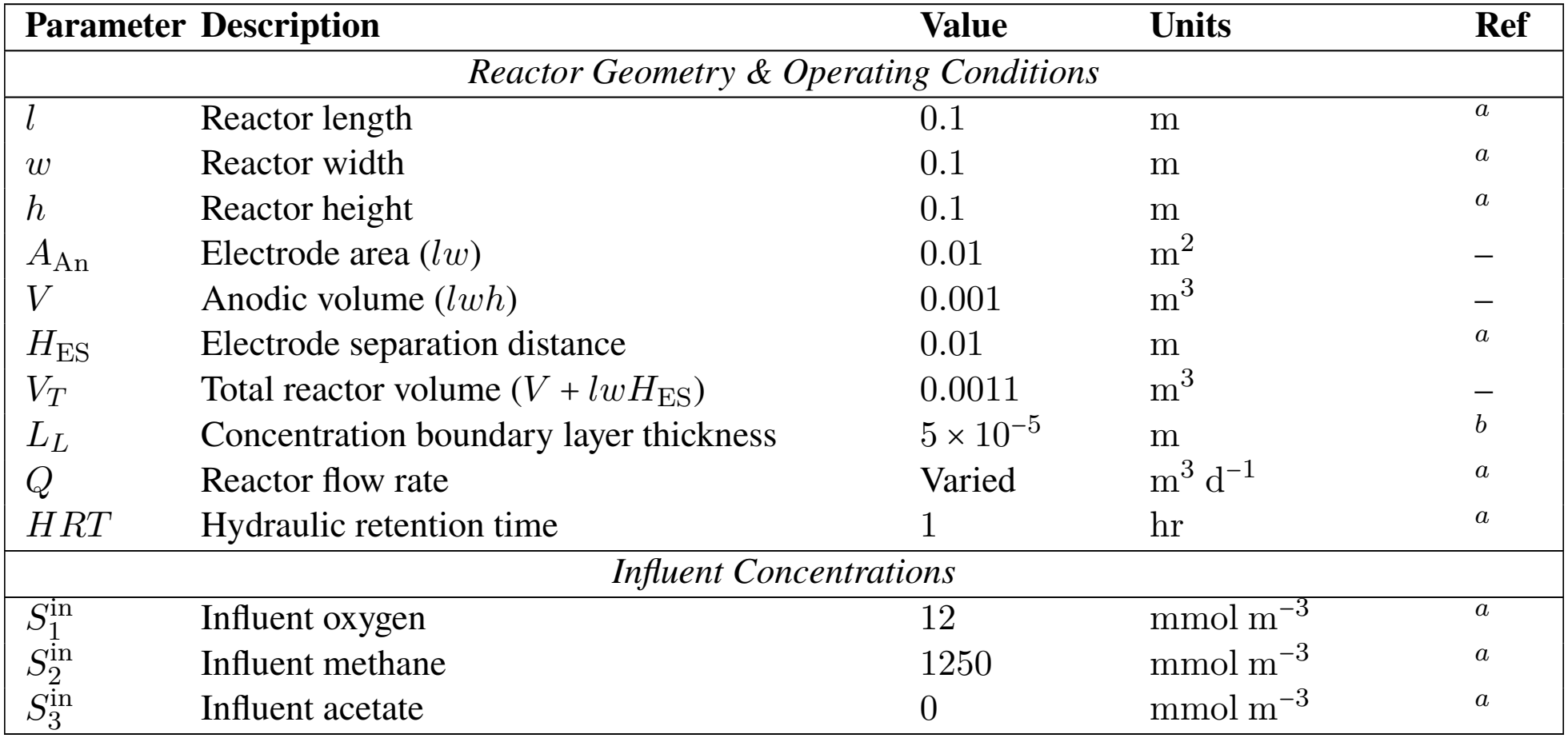
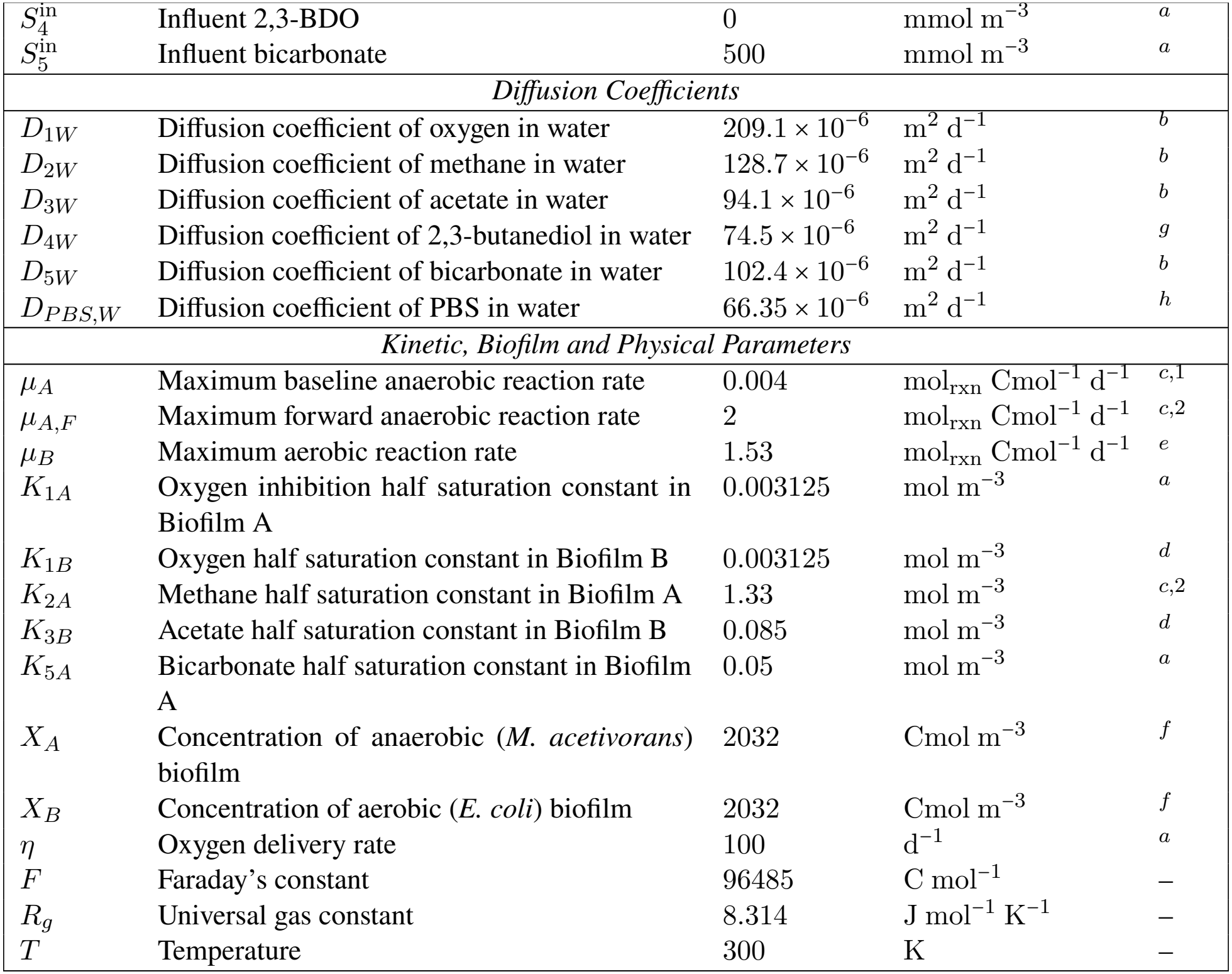
Model Parameters, Reactor Dimensions, and Operating Conditions. We assume *H*_ES_ includes anode thickness, membrane thickness, and cathode thickness. For baseline parameters inline with typical biofilm assumptions, *D*_*iA*_ = 0.8 *D*_*iW*_ and *D*_*iB*_ = 0.8 *D*_*iW*_ for all species *i*. ^*a*^ Assumed parameters; ^*b*^ [9]; ^*c*,1^ acetate production rate in reverse methanogenesis pathway from [29]; ^*c*,2^ forward methanogenic pathway from [48], used in reverse to establish an upper bound; ^*d*^ [64]; ^*e*^ acetate consumption rate from [51]; ^*f*^ Biomass concentration in units of Cmol m ^−3^ using the empirical formulation CH O N with a molecular weight of 24.6 g Cmol ^−1^ dry weight biomass, where 50 kg m ^−3^ is adopted from [30] as 5% assumed active biomass of the wet total biofilm density of 1000 kg m ^−3^, accounting for higher pore space for transport of redox mediators [52]; ^*g*^ computed using diffusion coefficient for 1,3-BDO at 298 K [60]; ^*h*^ For simplicity we take the average of 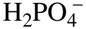 and 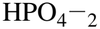 (occurs at pH = 7.2) to obtain the diffusion coefficient [45].

#### 3.2.2 Material Balance in the Biofilm

The chemical equations that govern the proposed biofilms found in Subsection 3.1 are restated here:

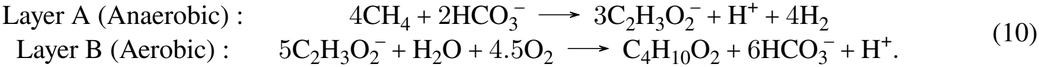

The total equation of the reactor after simplifying:

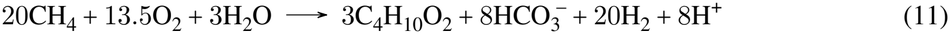

For simulations, the (10) equations ignore H_2_O and H^+^ (balanced in only C atoms!). The next set of equations represent chemical species reaction-diffusion in the biofilm, for all *i* ∈ {1, … , 5} are:

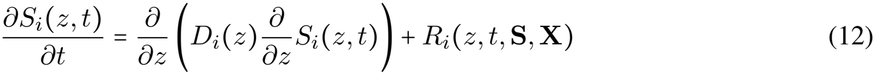

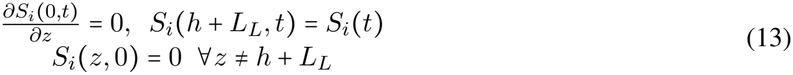

where is *D_i_*(*z*) the space dependent diffusion coefficient and *R_i_* is the reaction rate of substrate found in Table 2, **S** = (*S*_1_, … , *S*_5_) and **X** = (*X*_*A*_, *X*_*B*_) is the set of biomass concentrations in Layer A, Layer B respectively. At each interface we use the conditions (18). Each diffusion coefficient and maximum reaction rate has a jump discontinuity at interface boundaries. It is important to note the equation for 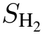 is an ODE governing the amount of moles created with initial condition 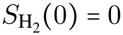. An example of this system in batch mode can be seen in Figure 2.

**Figure 2.**
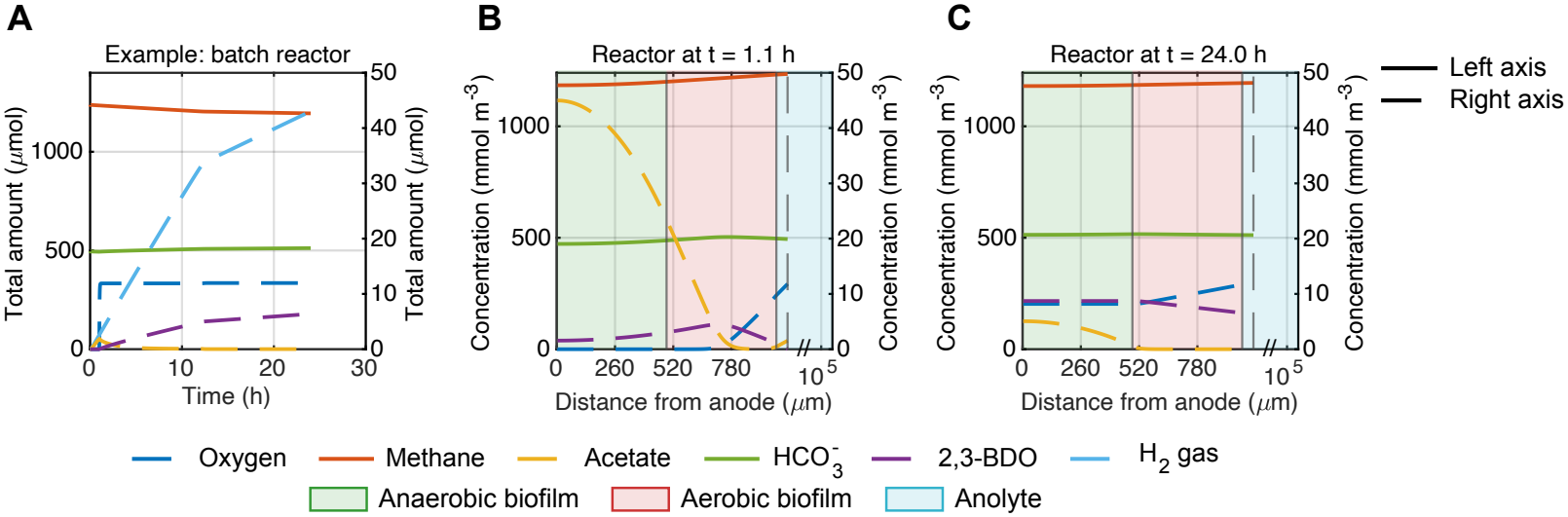
Oxygen inhibition in the anaerobic biofilm is triggered by the decreasing rate of AOM in a batch cultivation simulation of the DL-MEC. **A.**An example batch simulation with for all six substances in the DL-MEC *M*_*i*_ (1). **B**. The molar concentration of five species in the biofilm at *t* ≈ 1.1 hours where the aerobes shield the anaerobes from oxygen inhibition. **C**. The molar concentration of five species in the biofilm at the final time *T* = 24 hours =1 day after oxygen inundation. Biofilms of thickness *h*_1_ = *h*_2_ = 490 m up to *T* = 24 hours = 1 days.

**Figure 3.**
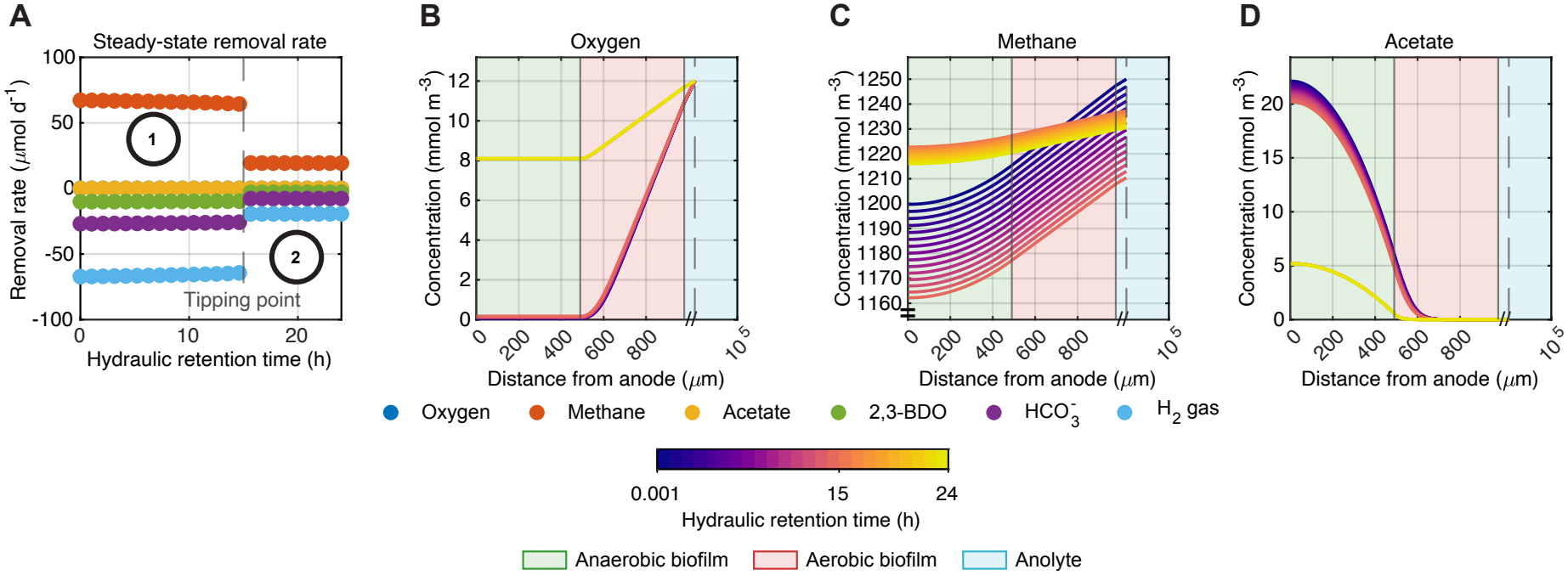
Regime change behaviour is influenced by steady state bulk methane to oxygen ratio. A.The tipping point (dashed line) delineates between two regimes defined by in the removal rate of all species in the tank, (1) oxygen minimized regime and (2) oxygen inhibited regime. **B**. The oxygen concentration profile in the biofilm at different HRTs. **C**. The methane concentration profile in the biofilm at different HRTs. **D**. The acetate concentration profile in the biofilm at different HRTs. On **B**., **C**., **D**. a red mark on the colormap marks the tipping point and the dashed line marking the transition from the CBL to the bulk anolyte. Biofilms of thickness *h*_1_ = *h*_2_ = 490 *µ*m for varied *T*_HRT_ ≈ 0 hours to 24 hours with *L*_*L*_ = 50 *µ*m.

**Figure 4.**
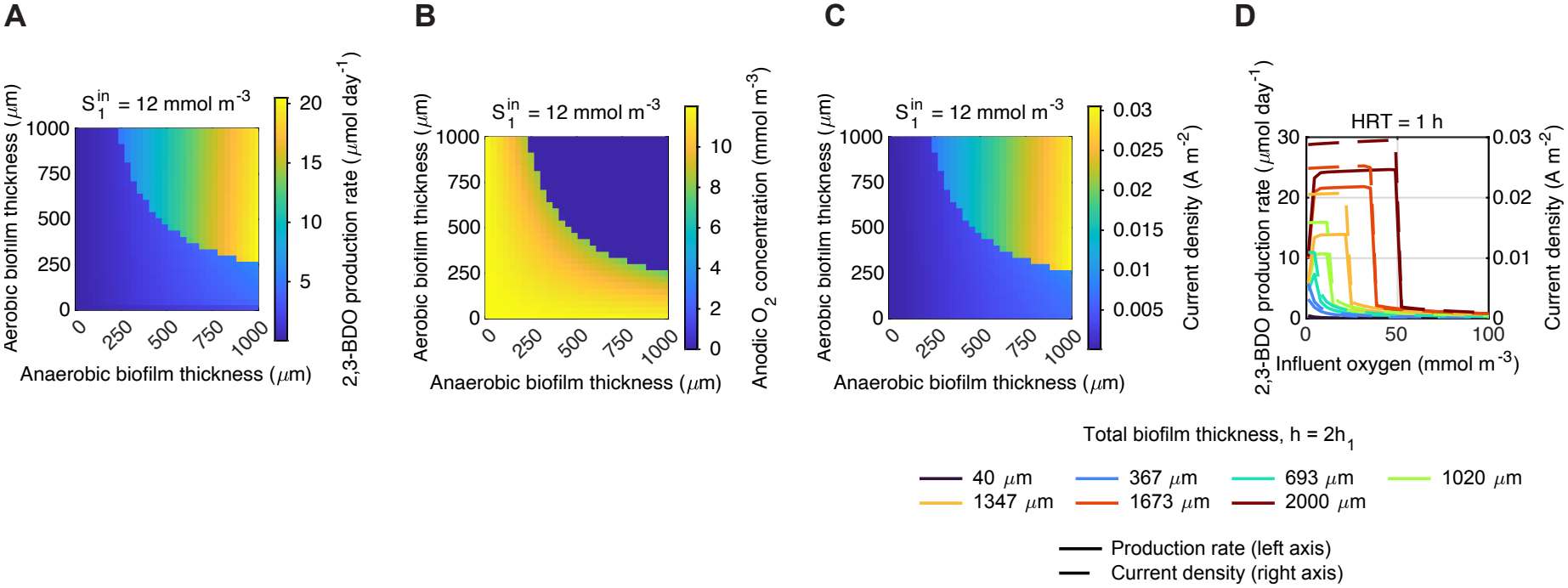
The production rate of 2,3-BDO is maximized where the highest acetate supply meets fixed oxygen demand. A.Production rate of 2,3-BDO. **B**. Oxygen concentration at the anode, **C**. Non potential limiting current density. **D**. Production rate of 2,3-BDO (left axis) and current density (right axis) for varied oxygen influent concentrations 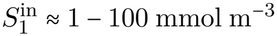 and varied biofilm thicknesses where *h*_1_ = *h*_2_ and *h* ≈ 20 − 1000 *µ*m. For **A**-**C** *T*_HRT_ = 1 hour for *S*_1_ = 12 mmol d^−1^. Legend is for **D**.

#### 3.2.3 Reaction Rates

Incorporating experimentally reported reaction rates into the model requires careful consideration of their units. The volumetric production or consumption rate of species *i, R*_*i*_ (*z, t*, **S, X**) has units mol species m^− 3^ day ^−1^, achieved by

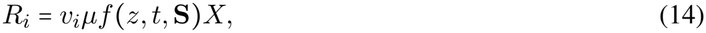

*µ* is the biomass specific maximum reaction rate, and *v*_*i*_ is the signed stoichiometric coefficient of species *i* in the corresponding biochemical reaction (Table 2) The stoichiometric coefficient has units

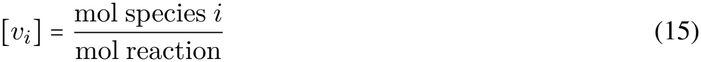

where one mol reaction means that the reaction occurs once according to the stoichiometric equation (Table 2). We define mol reaction as mol_rxn_ for the duration of the paper. *f*(*z,t*,**S**) is the dimensionless Monod kinetic scaling factor. Biomass is expressed on a carbon mole basis Cmol m ^−3^ which we denote as Cmol biomass m ^−3^ (Table 3). Accordingly, to establish consistency across species we convert each maximum reaction rate to units

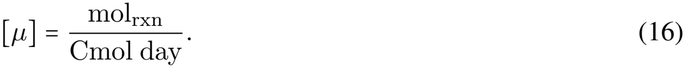

Therefore, the species specific maximum reaction rate is defined by

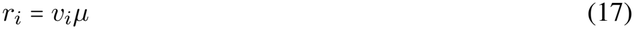

with units mol species Cmol ^−1^day^−1^. We write mol reaction as mol for the duration of the paper. This enforces the fixed stoichiometric yield yield and ensures conservation of carbon among chemical species. Detailed in Supplement S5 for rate specific derivation.

To convert grams biomass units to carbon mols, we assume 100% of CDW reported in grams represents active biomass. CDW has an empirical formulation of 24.6 g biomass Cmol ^−1^. In practice, CDW comprises both active and inert biomass. The reported reaction rate in [44],[51] is normalized to total CDW rather than active biomass alone, dividing by this larger denominator yields a specific rate that is lower than the true per cell activity. This underestimate establishes the reaction rate as a conservative baseline for a minimum feasibility scenario.

A unit conversion is also required, since [29] report their findings per mg total protein rather than CDW. Total protein is generally estimated at 50% of CDW for microbes [52]. Using this estimate, we multiply the specific reaction rate by this fraction to obtain a baseline reaction rate for reverse methanogenesis, expressed on a CDW basis. This yields consistent reaction rate and biomass units across species (Table 2).

The reaction rates considered are centered on acetate production and consumption to evaluate the central bottleneck of the system. By choosing the theoretical catabolic yield (stoichiometry) and equating 2,3-BDO production to acetate consumption, we overestimate the conversion efficiency relative to observed yields and production rates. This provides an upper bound estimate on the maximum production per unit substrate. Combined with conservative acetate production and consumption rates, these assumptions allow for the direct assessment of DL-MEC feasibility. By targeting the coupled reactions between the two species we establish the minimal reaction rates where product synthesis is achievable.

#### 3.2.4 Numerical Methods (for Stationary Biofilm)

The DL-MEC model consists of 2 sets of equations: semilinear parabolic partial differential equations with variable coefficients in the form of a reaction diffusion system with static biofilm layers and ordinary differential equations governing the continuously stirred tank. Due to diffusion and the reaction mechanisms, the biofilm equations are stiff. The number of biofilm equations represents the number of reactants and products. Using the methods of lines with a centered finite difference method in space for the interior nodes with a matlab package *ODE15s* for an ode integrator in time, discretized points are placed directly on the interface between biofilms (or CBL-biofilm). We assume matching conditions at these interfaces to be [27]

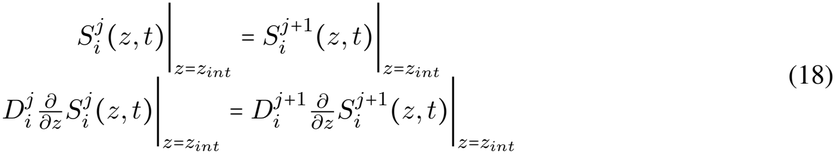

for the *ith* species and the *jth* layer where *z*_*int*_ is the interface node. Due to the interface nodes, we employ a nonuniform mesh with forced continuity of flux and continuity of concentration. Mesh convergence was verified against an analytical steady-state solution for a simplified reaction-diffusion system. Results are presented in Supplement S4 (Fig. 5,6, Table 2). The sensitivity analysis uses the Morris method [14], Sobol sequences from [55] and in house data analysis codes with parameters and verification in the Supplement S3 and S4 (Fig. 7,8, Table 2).

**Figure 5.**
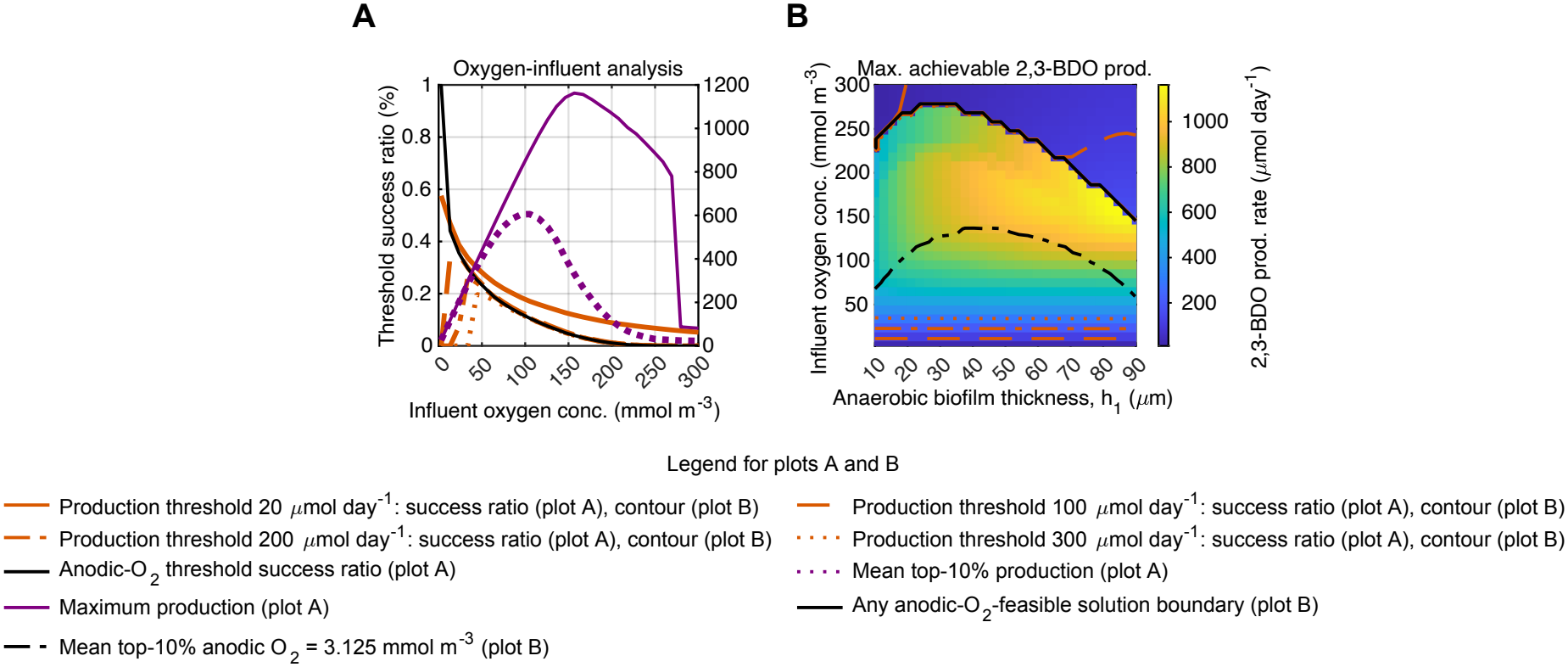
Increased oxygen concentrations trigger higher anodic inundation risk yet can maximize 2,3-BDO production potential, while lower oxygen concentrations decrease inundation risk but restrict production potential (h = **100** *µ*m). A.The threshold success rate and 2,3-BDO production rate at corresponding oxygen influent values for a Sobol sequence of the reaction rates and biofilm densities. **B**. The maximum production potential and corresponding thresholds on the biofilm ratio, oxygen influent concentration grid. Prod = Production, Max = Maximum, Conc = Concentration.

**Figure 6.**
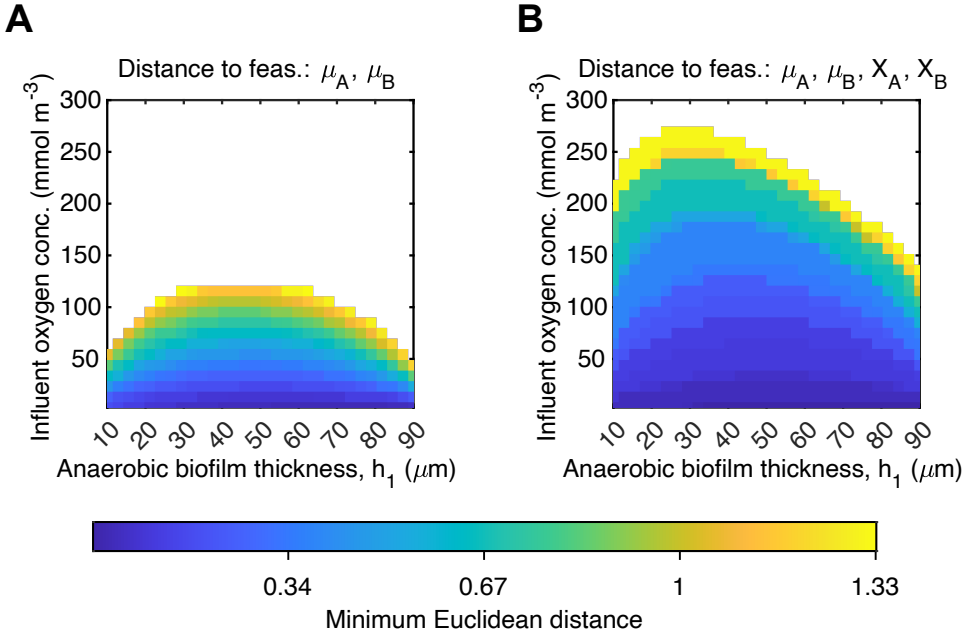
The required increase in maximum reaction rates to surpass anodic oxygen and 2,3-BDO production rate (20 *µ*mol d ^−1^) thresholds is minimized by optimizing both reaction rates and biofilm densities. A.The normalized Euclidean distance from the baseline parameters to the closest point in the productivity region while varying only the two reaction rates. **B**. The normalized Euclidean distance calculated using only the reaction rate components of the closest point in the 4D productivity region while varying both reaction rates and biofilm densities. White background implies no point in the space is feasible. Feas=Feasibility

**Figure 7.**
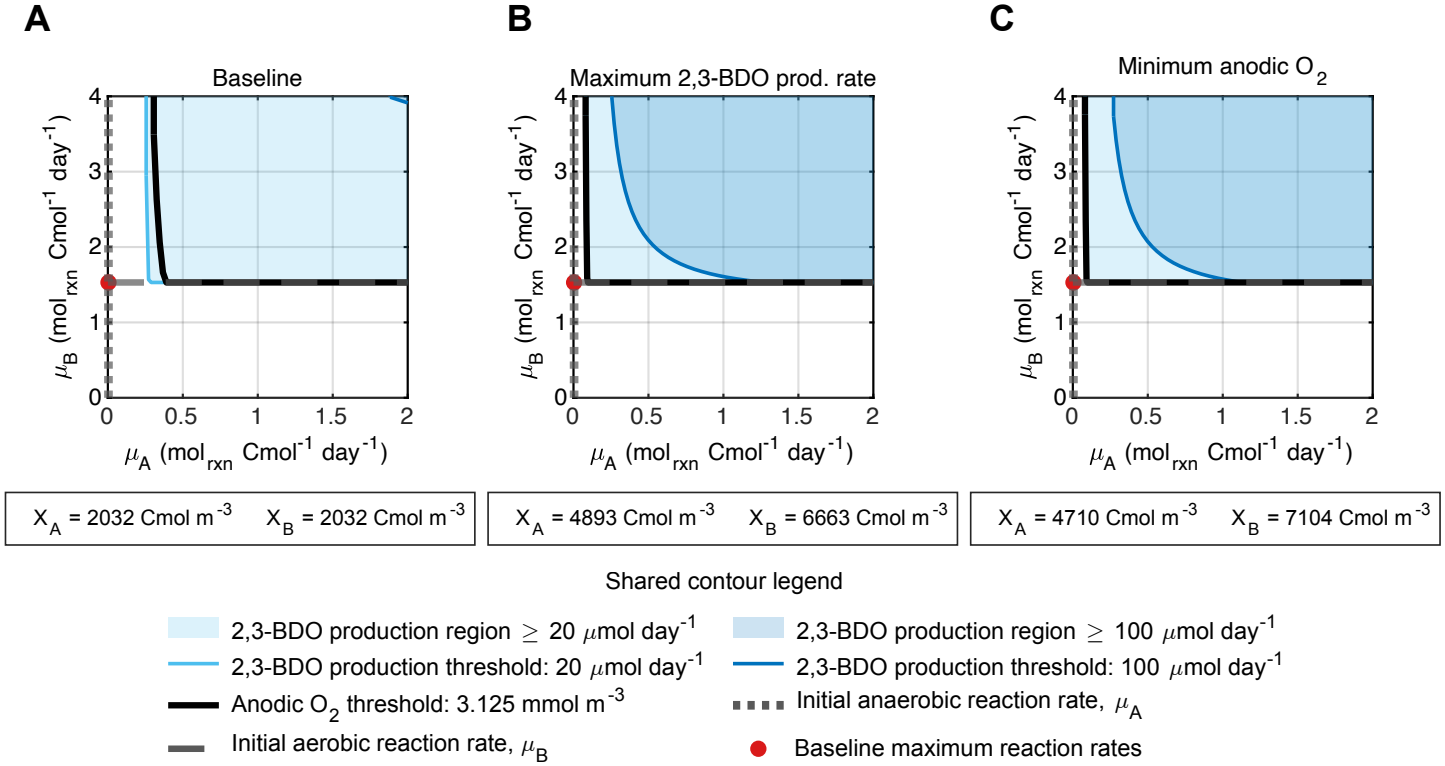
Reaching the productivity regime is enabled by increases in the AOM maximum reaction rate for both near-optimized and baseline biofilm densities. A.A contour trace of maximum reaction rates with baseline densities. **B**. A contour trace of maximum reaction rates corresponding to the mean biofilm density of the top 10% highest production rates. **C**. A contour trace of maximum reaction rates corresponding to the mean biofilm density of the 10% of points that minimize the anodic oxygen concentration. All traces are executed at 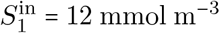, *h*_1_ = 55*µ*m, *h*_2_ = 45*µ*m.

**Figure 8.**
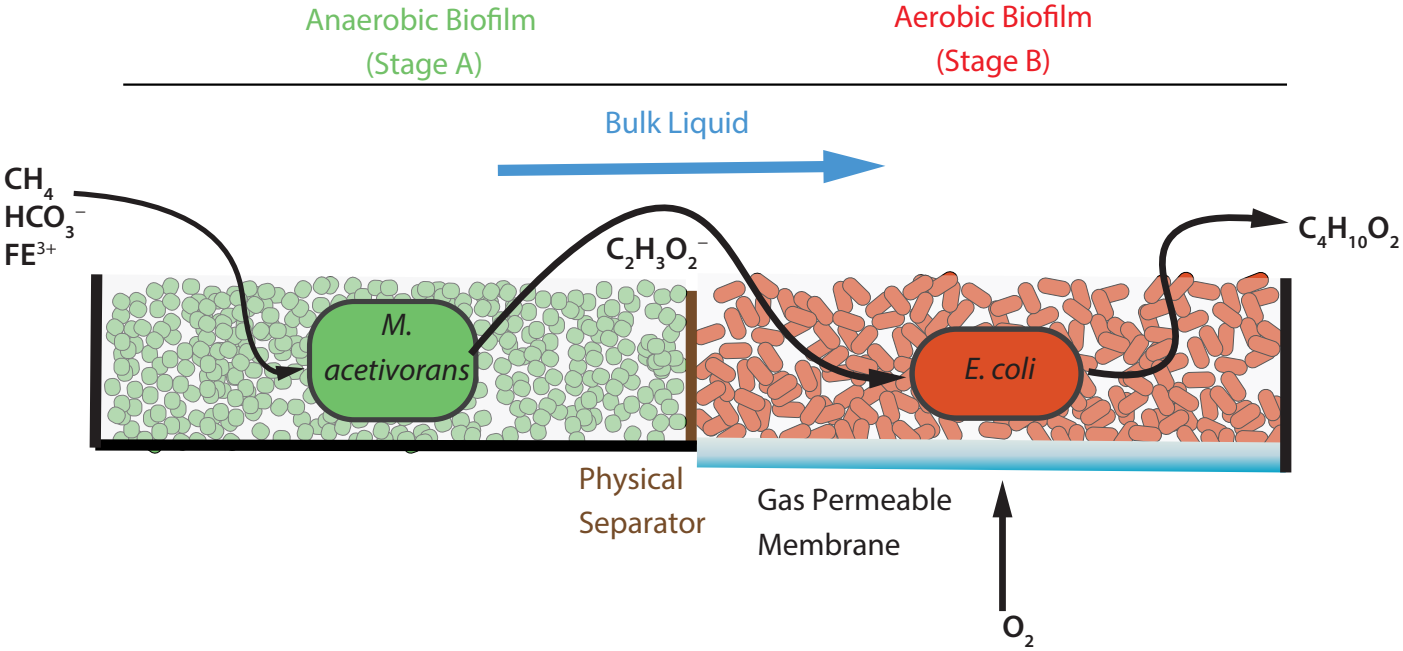
A biologically practical reactor design for AOM and product synthesis enabled by oxidative and biofilm longitudinal partitioning. The longitudinal partitioned flow reactor (LPFR) mitigates oxygen inhibition at the expense of increasing mass transfer resistance.

### 3.3 Electrochemical and pH Considerations

Our primary focus of the DL-MEC is value-added production with secondary goals of hydrogen production and AOM. In order to do so, we must maximize the current DL-MEC to drive the DL-MEC as close to the electric potential non-limiting regime as possible. Therefore, in designing our reactor we must consider the overpotential accrued by our system with specific focus on ohmic overpotential. We discuss the electrochemical components including the applied voltage, current and resistance expected of the DL-MEC. The applied voltage *E*_applied_ required for our system [48]

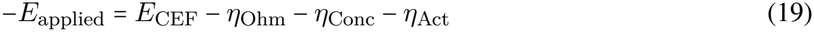

where *E*_CEF_ is the counter electromotive force *E*_CEF_ = − 0.295 *V, η* _Ohm_ = *η*_Ohm,Cath_ + *η*_Ohm,An_, is the ohmic overpotential, *η*_Conc_ = *η*_Conc,Cath_ + *η*_Conc,An_ is the concentration overpotential and *η*_Act_ *η*_Act,Cath_ *η*_Act,An_ is the activation overpotential. We focus on ohmic overpotential which is most controllable: the electrokinetic limitations and current generation governed by methane oxidation. While activation overpotential is present when building the reactor, we do not consider it here and concentration overpotential is handled directly by the model. Ohmic overpotential is defined by

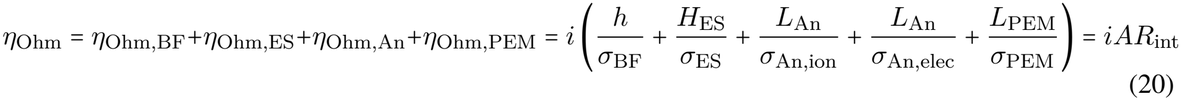

where *i* is the current density, *A* is the biofilm area, *η*_*i*_ represent the ohmic overpotential of the biofilm, catholyte, anode, and proton exchange membrane respectively, *L*_*i*_, is the electrode separation (ES), anode and PEM thickness and distance *σ*_*i*_ is the conductivity of the biofilm, the catholyte solution between electrodes (ES), the ionic and electrical conductivity of the porous anode and the conductivity of the PEM respectively. All specific values and derivations can be found in the Supplement S1 (Table 1). The sum of the resistance terms are group together to define *R*_int_ and the current density is

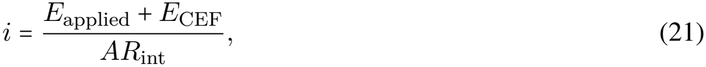

Without wasting additional energy, we aim to drive up the current by minimizing *R*_int_, starting with the conductivities of the electron transfer mechanism.

The central EET mechanisms for EET in *M. acetivorans* are nanowires and diffusible redox mediators [5]. However, *M. acetivorans* does not possess nanowires. With the current biotechnology, diffusible redox mediators could be employed, while incurring significant ohmic overpotential due in particular to biofilm thickness. We estimate a conductivity of *σ*_*BF*_ ≈ 7. 5 × 10 ^−4^ S m ^−1^ (estimated in Supplement S1), which is significantly lower than the nanowire case (*σ*_*BF*_ ≈ 1.25 S m ^−1^ [66]). Hence, for a redox mediated system minimizing biofilm thickness and maximizing mediator efficiency is more critical than optimizing other reactor design features such as electrode spacing or buffer conductivity. Using a co-culture with *Geobacter sulfurreducens* natively expressing nanowires, it has been shown that with *Paracoccus denitrificans* (as an electron shuttle) and *M. acetivorans* oxidizing methane, a current was established for electron transfer [39]. With the addition of humic acids as electron mediators the electric current greatly increased current and oxidation of methane [68]. Thus biofilm resistance plays a much smaller role in ohmic overpotential for a system utilizing nanowires, however, the extent to which conductivity is restricted requires further experimental and computational analysis.

The current density (*i*) utilizing DL-MEC reaction-diffusion equations in a kinetic substrate limited regime (where 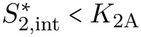), non-electric potential limited regime, with a first order approximation of the Monod kinetics is

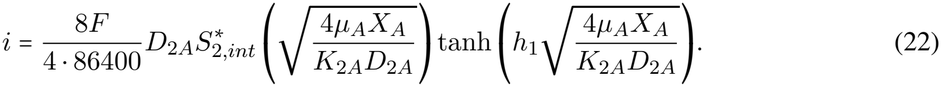

For a system with such assumptions we can estimate a lower bound of the biofilm ohmic losses for both systems 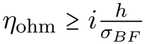. A current density of *i* = 0.1120 A m^−2^ from (22) for *h*_1_ = 50 *µ*m and *h*_2_ = 50 *µ*m with 7. 5 × 10 ^−4^ S m ^−1^ and 1.00 mol m ^−3^, the ohmic overpotential (20) using the redox mediated conductivity *η*_*Ohm*_ ≈0.0149 V, less than 10% of the CEF for the cell while using a co culture is *η*_*Ohm*_ ≈ 1.4 × 10 ^−5^ V significantly less than 1% of the CEF for the cell. At this conductivity using nanowires, ionic conductivity of the anode becomes current limiting. Even under mediated EET, biofilm ohmic losses remain under 10% of the CEF, though the nanowire case reduces this by three orders of magnitude. This is likely an overestimation of the conductivity, yet this value exemplifies the need to utilize a thin biofilm, particularly for mediated systems. While use of nanowires support the non-limiting potential assumption, in mediated pathways biofilm overpotential remains limiting. Ultimately, local reaction rates remain bounded by unavoidable activation, ohmic overpotential and concentration overpotential, some of which is pH related.

For each 4 molecules of methane oxidized, net 9 protons are produced and must diffuse from the biofilm through the porous anode, PEM and electrode gap (catholyte) to the cathode. We choose a highly conductive saline catholyte solution [66]. If any remaining protons from oxidation accumulate, the increased acidity will harm the methanogens decreasing AOM and current. Since the primary channel for ionic transport and conductivity is in the electrode gap which does not consist of the biofilm, the biofilm matrix can be optimized for biological viability by utilizing phosphate buffer solution (PBS) without incurring larger cell resistance losses due to lower conductivity solution of PBS. For instance, with an assumed 2,3-BDO production of 20 *µ* mol d ^−1^, acetate consumption rate of 100 *µ* mol d ^−1^ which is equal to acetate production rate. The proton production rate in the biofilm is 320 *µ* mol d ^−1^ while proton consumption rate at the cathode is approximately 266 *µ* mol d ^−1^. Therefore, for a biofilm thickness of 100 m, a current of *i* = 0.0297A m ^−2^, continuous PBS feed concentration of 10 mM will have the replenish rate of 5 28 mmol d ^−1^. This feed rate is greater than the required 54 mol d^− 1^ thus, significant acidification is unlikely to be limiting even as the AOM reaction rate increases. Even at the current found using *i* 0.1120 A m ^−2^ (22), with net surplus of proton four times the amount above, the PBS still has a replenish rate significantly outpacing proton production [42].

## 4 Numerical Simulations and Results

We analyze the DL-MEC design at laboratory scale to determine its potential practical feasibility using the developed model. The geometry considered consists of square electrodes of area 0.01 m^2^ with a tank of volume *V*_*T*_ = 0.0011 m^3^ as seen in Table 3.

### 4.1 Baseline Results

The total amount of a given substance in the tank at a given time *t* in moles is

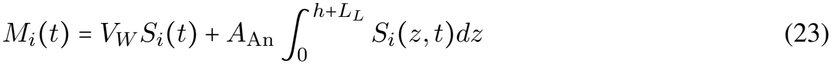

where *V_W_* = *V* − *A*_An_ (*h* + *I_L_*). An example of this, for a batch reactor, can be seen in Figure 2. In the baseline simulations we assume *h*_1_ = *h*_2_ = 490 *µ*m, and we use *A*_anode_ with initial conditions under influent concentrations in Table 3. Complete conversion of methane would result in three moles of 2, 3-BDO for every 20 moles of methane, however due to oxygen inhibition when methane decreases this will not occur.

The batch simulation (Fig. 2) exemplifies an important system characteristic: low acetate concentrations result in oxygen penetration deep into the anaerobic biofilm. This occurs at the initial stage of cultivation and during the production stage. The DL-MEC requires a delay of oxygen influent allowing for acetate accumulation in the aerobic layer. During this initial anaerobic phase, *E. coli* (a facultative anaerobe) survives in a dormant phase (or on some secondary sugar source). *E. coli* rapidly shifts to aerobic respiration utilizing the accumulated acetate upon oxygen delivery. As methane concentrations drop, a cascading effect results in oxygen penetration deep into the anaerobic biofilm.

Two design choices, (i) continuous or fed-batch cultivation, (ii) delayed oxygen delivery, are motivated by the necessity to sustain steady high acetate concentrations throughout cultivation. Further, the biofilm acts as a steady-state catalyst for the operational period. When oxygen reaches a certain concentration in the anaerobic biofilm, significant inhibition of AOM occurs. This exemplifies a tipping point in the DL-MEC between the ‘production/oxygen minimized regime’ and ‘oxygen inhibited regime’ which is the central guiding bottleneck for the sensitivity analysis. Embedded in the tipping point is a trade off, where due to counter diffusion, together the substrates (acetate and oxygen) cannot simultaneously maximize the reaction rate.

The oxygen inhibited regime can also arise in continuous cultivation if the oxygen influent is too high or the methane influent too low, both of which reduce the acetate available to sustain aerobic shielding. This is best exemplified by changing the HRT (Fig. 3). Both regimes are best described by the supply of acetate concentration versus demand of oxygen reduction and highlight a simple yet important dichotomy of biofilm thickness. The *E. coli* biofilm must be thick enough to reduce oxygen but not so thick to limit 2,3-BDO production and increase internal resistance for electric conductivity. Our two main objectives are opposed, high enough oxygen deep into the aerobic biofilm to maximize synthesis of 2,3-BDO but not so high that it enters the anaerobic biofilm.

### Influent Concentrations

To establish consistent comparisons across sensitivity analysis and numerical experiments, we fix several parameters, including HRT, and some influent concentrations for tractability. Such choices of fixed parameters are constrained by theoretical estimates on the DL-MEC. While the HRT is typically application driven, determined by factors such as biomass washout, gas stripping and attachment/detachment, we do not dynamically consider any of those in our model. Instead HRT acts as a proxy for effective concentration influent in order to reasonably determine the influent and effluent concentrations. We select *T*_HRT_ = 1 hr and match it to the oxygen influent time delay of 1 hr. This ensures that oxygen supplied to the DL-MEC occurs after a single residence time has elapsed, maintaining anaerobic conditions in the anaerobic layer prior to aerobic respiration to allow acetate to accumulate. The concentration of acetate, as the coupling chemical between microbes, is the main bottleneck for system feasibility. In practice, the reactor would run continuously over long periods, so the specific numerical values chosen for HRT and oxygen delay are less critical than the underlying design principle they represent: allowing sufficient anaerobic residence time for acetate accumulation before oxygen exposure. This principle, rather than the precise 1 hr value, helps operators to establish a startup protocol for initial biofilm growth.

To minimize oxygen inhibition and sustain oxygen consumption in the aerobic layer, we choose a methane influent to maximize acetate production. The maximum dissolved methane concentration is 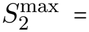 1250 mmol m ^−3^ at ambient pressure and temperature. Assuming a conservative liquid-phase mass transfer coefficient of 100 h ^−1^ [66], the maximum molar methane gas to liquid volumetric flux is roughly 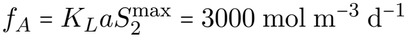. This serves as the upper bound for our methane supply. The influent methane concentration is set to 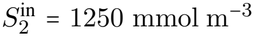. In principle, the methane influent concentration could be controlled dynamically to minimize acetate effluent. In practice, this is exceedingly difficult and not centrally relevant as feasibility hinges on dynamic supply and demand in the aerobic biofilm. Regardless of the HRT, this influent condition provides the maximum possible driving force for the system without concerns that we violate physical laws. To directly compare to the methane volumetric supply in the non electric potential limiting regime, the maximum consumption rate is 0.008 mol m ^−3^ d ^−1^. Even for an increase of 100 times, this value is below the maximum gas to liquid mass transfer limits, demonstrating that the methane gas-to-liquid transfer will remain non-limiting even when the system operates under electric potential limitations.

The influent bicarbonate concentration is set to 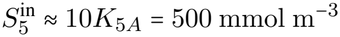. Bicarbonate, produced in the aerobic layer and consumed in the anaerobic layer, is highly soluble [53]. Since the aerobic reaction rate has a baseline significantly higher than the anaerobic rate, ample bicarbonate is expected to be available in most operating scenarios. For the last of the nonzero influent concentrations, we define the range of oxygen values considered in the paper. The maximum oxygen concentration is 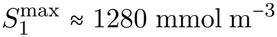. Based on preliminary sensitivity analysis, we choose a maximum oxygen influent of 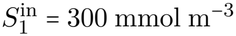 which is significantly below its maximum saturation limit 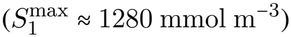. Anything above this value resulted in an infeasible system for the considered parameter bounds. A minimum oxygen influent value of 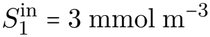 (1% of the maximum value) approximately equal to the half maximal oxygen concentration considered representing the lowest inhibition of AOM physically relevant. Varying oxygen concentrations are subsequently analyzed for their impacts on the DL-MEC. The influent concentrations for all chemical species are summarized in Table 3.

### Optimal Layer Thickness in Continuous Cultivation

To evaluate the plausibility of such a layered design we analyze the production rate surface of 2,3-BDO over varied layer thicknesses. We assume that 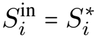 for all chemical species *i*. The removal rate of a chemical species, *S*_*i*_, is defined

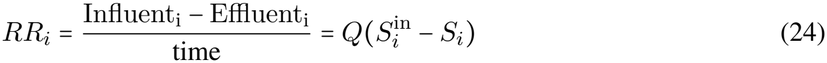

measured in *µ*mol d^−1^ with initial conditions found in Table 3. The production rate of 2,3-BDO is *PR*_4_ = ∣*RR*_4_∣.

Maximum production occurs for the largest *h*_1_ implying that the central bottleneck of production is acetate production. Maximum production (per unit thickness) is achieved for the smallest aerobic biofilm that satisfies supply and demand as close to the biofilm-biofilm interface without inundating the anaerobic biofilm with oxygen. As the anaerobic biofilm thickness increases along the tipping boundary, maximum production increases, indicating that 2,3-BDO synthesis is not saturated within the aerobic layer even when acetate supply is sufficient to meet oxygen demand. Subsequent analysis has shown when acetate supply sufficiently meets oxygen demand and *µ*_*A*_ ≈ *µ*_*B*_, maximum production occurs along the edge of the tipping point (for *h*_2_ ≤ *h*_1_) (Fig. 1 in Supplement S2) as the aerobic reaction rate is not yet saturated. If the aerobic biofilm thickness is larger than the anaerobic biofilm, we may achieve the production region at the expense of a desired production rate (Fig. 4**A**). As oxygen influent and total thickness decrease, so does production (Fig. 4**D**). At low enough values there is no discernible tipping point. However, biofilms of this thickness *h* ≥ 150 are not biotechnologically and electrochemically achievable. Hence, rather than simply decreasing oxygen influent, we must tune different characteristics of the reactor.

### 4.2 Sensitivity Analysis for Synthetic Biology

Biofilms thicker than 100–150 *µ*m are biotechnologically infeasible for a MEC due to large ohmic losses, lessened current and shear due to agitated mixing [42]. The current biofilm values reached are too large. To address this we perform a multi-stage sensitivity analysis to answer (i) how varying parameters impact our productivity and oxygen inundation, (ii) what parameter changes are required to achieve reactor feasibility for a reasonable biofilm thickness, (iii) what parameters have the most leverage to achieve engineering targets. The analysis is presented as follows: (i) analytical justification for the necessity to decrease biofilm thickness, (ii) evaluating individual impact of parameters on the final metrics, (iii) global scale production and oxygen analysis over a range of parameters, (iv) quantification of “effort” to engineer, (v) a trace of one specified oxygen and biofilm thickness set to fully digest the impact of parameter changes. This analysis will point towards the conclusion that, optimized biofilm densities with some reaction rate increases (particularly for the anaerobes) result in attainable feasibility.

For the aerobic biofilm, assuming suitably high acetate (*S*_3_ = 5*K* _3*B*_) , a first order approximate minimum depth of penetration of the oxygen profile (*δ*_*B*_) from the edge of the biofilm *h* assuming *S*_1,*BF* −*CBL*_ >> *K*_1*B*_) can be estimated 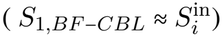

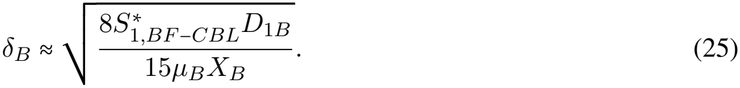

The minimum aerobic biofilm thickness for *S*_1_ = 12 mmol m ^−3^ is 19 and 300 mmol m ^−3^ (the upper bound of our considered range) is *h*_2_ ≈ 93 *µ*m. These two estimates show (i) for *S*_1_ = 12 mmol m ^−3^, as used in the motivating examples, a viable biofilm thickness can be reached assuming sufficient acetate, (ii) it requires over 100-fold increase in oxygen influent to render the aerobic biofilm thickness infeasible. However, acetate is unlikely to reach such values in the aerobic biofilm and its production rate is thus the central limiting bottleneck. The first order estimate identifies relevant limiting behavior but fails to capture the true scope of parameter interactions.

The parameters explicitly considered for our sensitivity analysis are: oxygen influent concentration *S*_1_, which plays a significant role in both the sharpness of the tipping region and total productivity (Fig. 4); maximal reaction rates *µ*_*A*_, *µ*_*B*_; biofilm densities *X*_*A*_, *X*_*B*_, which enhance the catalytic potential at the expense of substrate transport; and biofilm thickness ratio between the microbial layers, with the constraint *h* = *h*_1_ + *h*_2_ = 100 *µ*m. In the DL-MEC, if oxygen was simultaneously varied with the other parameters, the effects would dominate the more subtle changes of the microbial reaction rates and densities. For investigation of the system sensitivity, the biological adjustable parameters are separated from the operational parameters. To maintain parity with foundational bioelectrochemical models [30, 46] and classic biofilm bioreactor modeling [9], a baseline diffusivity for each species is of *D*_*BF*_ 0.8*D*_*iW*_. As the biofilm density varies, it impacts effective diffusivity of chemical species in the biofilm, which the estimate (25) misses. Therefore, local diffusion is dynamically coupled to the biofilm’s void fraction using the Mackie and Meares equation [37]

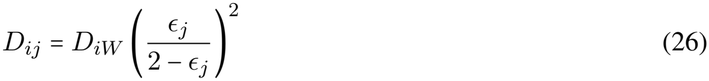

where *i* is the chemical species and *j* is the biofilm layer. At the baseline active biomass density (2032 Cmol m ^−3^) , the corresponding porosity (*ϵ* ≈ 0.95) approximates the 0.8*D*_*iW*_ diffusivity constraint. As simulated biomass density varies during sensitivity analysis, local porosity adjusts proportionally, smoothly varying the effective diffusion coefficient.

At each fixed point over a grid of influent oxygen concentrations and biofilm ratios, many samples are taken of the four parameters *µ*_*A*_, *µ*_*B*_, *X*_*A*_, *X*_*B*_ bounded by established parameter limits (Table 4).

**Table 4:** Parameter values for global sensitivity analysis for *h* 100 *µ*m. All parameters share common baselines and ranges (columns 2–3); analysis-specific deviations are noted in columns 4–7.

| Parameter | Baseline | Range | Analysis-Specific Values |  |  |
|---|---|---|---|---|---|
|  |  |  | Threshold Success Ratio (Fig. 5) | Distance (Fig. 6) | Contour Trace (Fig. 7) |
| *S*<sub>1</sub><sup>in</sup> | 12 mmol m<sup>−3</sup> | 3–300 mmol m<sup>−3</sup> | varied | varied | 12 mmol m<sup>−3</sup> |
| *rat* = *h*<sub>1</sub>/*h* | *h* = 100 *μ*m | 10%–90% | varied | varied | 55% (fixed) |
| *μ*<sub>*A*</sub> | 0.004 mol<sub>rxn</sub> Cmol<sup>−1</sup> d<sup>−1</sup> | 0.004–5 mol<sub>rxn</sub> Cmol<sup>−1</sup> d<sup>−1</sup> | varied<sup>*</sup> | varied<sup>*</sup> | varied<sup>‡</sup> |
| *μ*<sub>*B*</sub> | 1.53 mol<sub>rxn</sub> Cmol<sup>−1</sup> d<sup>−1</sup> | 1.53–5 mol<sub>rxn</sub> Cmol<sup>−1</sup> d<sup>−1</sup> | varied | varied | varied<sup>‡</sup> |
| *X*<sub>*A*</sub> | 2032 Cmol m<sup>−3</sup> | 500–8000 Cmol m<sup>−3</sup> | varied | **A**. fixed<sup>†</sup> / **B**. varied | **A**. fixed<sup>†</sup> / **B**. 5115 Cmol m<sup>−3</sup> |
| *X*<sub>*B*</sub> | 2032 Cmol m<sup>−3</sup> | 500–8000 Cmol m<sup>−3</sup> | varied | **A**. fixed<sup>†</sup> / **B**. varied | **A**. fixed<sup>†</sup> / **B**. 6318 Cmol m<sup>−3</sup> |
| Production Rate Threshold |  |  | 20, 100, 200, 300 *μ*mol d<sup>−1</sup> | 20 *μ*mol day<sup>−1</sup> | 20, 100 *μ*mol d<sup>−1</sup> |
| Anodic Oxygen Threshold |  |  | 3.125 mmol m<sup>−3</sup> | 3.125 mmol m<sup>−3</sup> | 3.125 mmol m<sup>−3</sup> |
| Hydraulic Retention Time |  |  | 1 hour | 1 hour | 1 hour |

“Varied” indicates the parameter is sampled over its full range. <sup>†</sup>Fixed at baseline value (2032 Cmol m<sup>−3</sup>). <sup>\*</sup> Varied on a log scale for sampling. <sup>‡</sup> Results shown on a limited range (*μ*<sub>*A*</sub> from 0.004 – 2 mol<sub>rxn</sub> Cmol<sup>−1</sup> d<sup>−1</sup> and *μ*<sub>*B*</sub> from 1.53 – 4 mol<sub>rxn</sub> Cmol<sup>−1</sup> d<sup>−1</sup>).

At each grid point, the objective functions *PR*_4_ and 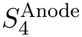 are considered using Morris global sensitivity analysis and the monotonicity index [14, 41]. We refer to Supplement S3 for full analysis but summarize the results here. The 2,3-BDO production rate (Fig. 3a in Supplement S3) exhibits strictly increasing monotonic behavior with respect to *µ*_*A*_, *µ*_*B*_, *X*_*A*_. The maximum anaerobic reaction rate has the largest mean effect, while the aerobic reaction rate has the smallest. This implies *µ*_*A*_ is the key driver to surmount the tipping point. However, the reaction rates, as the biofilm density parameters, have high standard deviation implying volatility; *µ*_*A*_ is highly dependent on the values of the other parameters as a slight parameter shift can trigger a jump over the tipping point while others result in no jump (Fig. 2 in Supplement S3). *X*_*B*_, on the other hand, exhibits non-monotonic effects as it can decrease the production rate at high oxygen values by limiting oxygen flux.While the densities are not the primary drivers of peak production, (Fig. 2,3 in Supplement S3) optimizing them is critical for moving the ‘cliff edge’ to a biochemically more achievable state as their values fundamentally define the accessibility and size of the production plateau. The Morris analysis results including the monotonicity index show the two objective functions largely behave the same when parameters are perturbed.

Now we turn to a quantitative global evaluation aimed at answering the questions (i) at what oxygen influent are we most likely to achieve feasibility, (ii) at what oxygen influent are we to achieve maximum production. Using a deterministic Sobol sequence [50] we evaluate oxygen’s viable interval using the threshold success ratio (Fig. 5):

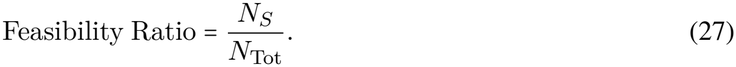

where *N*_Tot_ is the total number of Sobol points in the sequence and *N*_*S*_ is the number of Sobol points in the sequence surpassing a prescribed production rate threshold or anodic oxygen concentration threshold. Four thresholds are chosen for the 2,3-BDO production rate (20, 100, 200, 300 *µ*mol d^− 1^). The first, 20 *µ*mol d ^−1^ represents the maximum 2,3-BDO production rate reached at the infeasible biofilm thickness (Fig. 4). We then frame this threshold to answer the question how much engineering is required to achieve that same value at a feasible biofilm thickness. The production rates span from 20 mol d^− 1^ to a 15 fold increase production rate with thresholds at each 5-fold interval. The anodic oxygen threshold is set to 3.125 *µ* mmol m ^−3^, corresponding to the oxygen half-maximal coefficient for *M. acetivorans* (Table 3) representing when significant metabolic inhibition begins.

At low oxygen influent concentration, a high percentage of parameter combinations satisfy the threshold success ratio of anodic oxygen while failing to reach the lowest production threshold (Fig. 5**A**). At high oxygen influent the anodic oxygen defines feasibility region while low production rate fail to properly bound the region (Fig. 5**B**). This motivates the use of both thresholds together collectively defining an upper and lower bound on physically achievable parameters combinations.

As production rates increases, they converge to the anodic oxygen threshold, outlining the tipping region and indicating the sharpness of the tipping point. Therefore, particularly at higher oxygen influent, nearly all parameters that avoid oxygen inhibition achieve at least one of the defined production ratios.

Between the high oxygen and low oxygen influent, an intermediate or ‘transition’ region is defined. The lower bound of this region is defined by the mean 10% lowest anodic oxygen concentrations satisfying the anodic oxygen threshold. While the upper bound requires at least one parameter combination in the Sobol sequence satisfying the anodic oxygen threshold. In this region, a parameter combination achieving this production is considered an outlier among the Sobol sequence.

There is a clearly a defined oxygen influent region with engineering plausibility between 3 − 200 mmol m ^−3^ (Fig. 5**A**). However, as oxygen decreases, only smaller productivity thresholds can be achieved drifting farther and farther away from maximum productivity. The optimal region for minimum production viability (i.e. the two smallest production thresholds 20, 100 mol d ^−1^) occurs between 10 − 25 mmol m ^−3^. While the maximum productivity region for the top 10% of productivity rates occurs between 100 − 150 mmol m ^−3^. As a result, there is a clear tradeoff between engineering robustness and and maximum production potential. The maximum production peak occurs about *h*_1_ ≈ 75 *µ* m compared to *h* ≈ 25 *µ* m (Fig. 5**B**). This analysis gives us a general topological scan of the region for *h* = 100 *µ* m. The ideal operational region occurs between 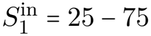 mmol m^−3^ where at least 20% of all ratios are achieved. However, to achieve at least baseline and 5x increase feasibility we focus on 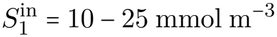.

The next analysis allows us to quantify the ‘effort’ required from baseline to improve the reaction rates with and without engineering microbial densities. We compare the normalized euclidean distance of engineering all four parameters to focusing on just the reaction rates. The smaller the normalized euclidean distance the closer the achievable threshold. To normalize the parameter values we use min-max normalization. Let *S_P R_* be the set of points in the productivity region and *S_OX_* be the set of points in the oxygen minimized region. Let 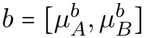 be the baseline data defined in Table 3. We consider two Sobol sequences: first *x*_1_ = [*µ*_*A*_, *µ*_*B*_, *X*_*A*_, *X*_*B*_] any point in the Sobol sequence that surpasses 20 mol d^−1^ and surpasses the anodic oxygen threshold and second *x*_2_ = [*µ*_*A*_, *µ*_*B*_] where *X*_*A*_, *X*_*B*_ are set to experimentally observed values. To consistently compare the euclidean distance from baseline of *x*_1_ and *x*_2_ for any point in the oxygen, biofilm ratio grid we use the weighted euclidean distance, defined by the matrix *W* , to the closest point of the viable operating regime is

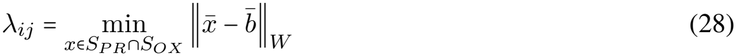

where 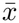 and 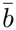 represent the normalized values. We project the four dimensional Sobol sequence into two dimensional space by strictly considering the resultant distance between reaction rates. Since we cannot define the weights as they are based on “effort to engineer” we assume *W* = *I*, the identity matrix.

As expected, the smallest euclidean distance to clear the minimum productivity threshold occurs at the lowest oxygen values (Fig. 6). Interestingly, the smallest euclidean distances occur from *h*_1_ ≈50 60 *µ* m as opposed to *h*_1_ ≈75 *µ* m which is better for maximum production. Since acetate is the limiting bottleneck, *h*_1_ ≈ 55 *µ* m allows for sufficiently satisfying oxygen demand without requiring massively large reaction rates, while *h*_1_ ≈ 75 *µ*m allows for potential acetate production as rates get to the highest levels of the parameter ranges. This mirrors what happened previously in the grid search (Fig. 4**A**). However, the right biofilm ratio is crucially important for higher oxygen influent but much more consistent at lower oxygen influent levels. Two crucially important features standout by comparing with and without optimized densities. Optimizing densities (Fig. 6**B**) both expands the region of feasibility for oxygen influent values considered and decreases the distance to the region.

We evaluate the specific required increases in microbial reaction rates with and without optimized densities using contour tracing. The considered optimized densities are (i) the top 10% of Sobol points the maximize the production rates (ii) the top 10% of Sobol points that minimize the anodic oxygen value. At low oxygen levels the use of both production rate and anodic oxygen thresholds is established by the feasibility analysis (Fig. 5**A**). The contour tracing is performed using a predictor-corrector algorithm consisting of the secant method (predictor) and bisection method (corrector). As an example, we use 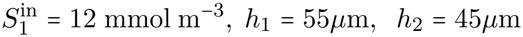 (Table 4). This method can be applied to all oxygen influent and biofilm ratio values where a change in these values alters the required increase to achieve feasibility.

To achieve the minimal production rate threshold and the anodic oxygen threshold, an increase of the aerobic biofilm reaction rate is not required (Fig. 7**A**). However, to achieve a 5-fold higher production rate, aerobic reaction rate increase and anaerobic reaction rate increase are required, even for optimized biofilm densities. At baseline, an increase of approximately 100-fold is required (from 0.004 to 0.39 mol_rxn_ Cmol ^−1^ d ^−1^) for the anaerobic reaction to achieve feasibility (Fig. 7**A**). While the production plateau cannot be mounted by optimizing the biofilm densities, a significant decrease in the engineering required to achieve that plateau is observed, corroborating previous analysis (Fig. 7**B,C**). To optimize for productivity, an increase of under 25-fold (from 0.004 to ≈ 0.095 mol_rxn_ Cmol^− 1^ d ^−1^) is required, while to optimize for minimization of anodic oxygen an increase of about 22-fold is required (from 0.004 to ≈ 0.09 mol_rxn_ Cmol ^−1^ d^− 1^). A distinguished difference between the optimized biofilm densities is not observed, further emphasizing the point that high production thresholds and anodic oxygen thresholds aim to achieve the same task. While we do not recommend to engineer only a single parameter, prioritizing *µ*_*A*_, the rate of AOM, is superior for feasibility at this point. While both rates necessitate optimization to achieve higher productivity, holistically optimizing the biofilm densities allows the DL-MEC to surmount the production tipping point with reasonable increases in the anaerobic reaction rate.

### 4.3 The Biotechnological Stepping Stone

The DL-MEC utilizes vertical stratification to govern distinct oxidative environments, producing a value added-product while avoiding mass transfer limitations between AOM and aerobic respiration. To effectively do so, reverse methanogenesis requires use of a non depletable anode. This is limited by efficient charge transfer to the anode. As established, this reactor, while outlined to be achievable with required engineering, is not currently practical for experimentally observed (Table 3). We introduce a second reactor that can serve as a gap between future feasibility and current infeasibility. This second reactor is used to contextualize DL-MEC performance against a currently feasible alternative; a detailed optimization is beyond the present scope.

We define the longitudinal partitioned flow reactor (LPFR) (Fig. 8), which (i) utilizes plug flow like reactor geometry through parallel plates utilizing advective flow with a much smaller tank volume, (ii) separates oxidative environments into longitudinal stages (anaerobic then aerobic) via physical mechanisms, (iii) uses an alternative method for oxygen delivery without a delay, (iv) uses ferric iron Fe^3+^ as the TEA[29],[56]

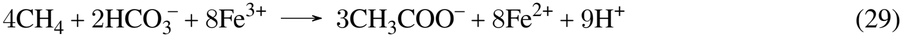

Unlike the DL-MEC where the anode-coupled reaction is endergonic, the iron-coupled AOM reaction is thermodynamically favorable 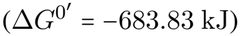, eliminating the need for an applied voltage.

We contrast several important features: (i) while the LPFR has a smaller volume, it suffers increased concentration boundary layer resistance due to laminar flow; DL-MEC limits traversing the boundary layer more than once, while LPFR requires three times, (ii) DL-MEC simultaneously recovers hydrogen gas, (iii) the LPFR requires an exceptionally thin biofilm for methane oxidation to be sustained at similar reaction rates due to diffusion limitations of insoluble iron, (iv) increased acidity of the biomass may occur in the aerobic biofilm as the protons are produced in the anaerobic biofilm and brought into the aerobic biofilm without the cathodic reaction, (v) oxygen gas is administered using permeable membrane that acts a substratum for the aerobic biofilm [38], (vi) influent iron, in particular as a particulate, may be caught in the bulk advective flow which in practice may warrant the use of an MEC/MFC for phase 1.

In the rest of the section, we introduce the model for the LPFR and compare results between the DL-MEC (Fig. 1**B**) and LPFR (Fig. 8).

### Longitudinal Partitioned Flow Reactor (LPFR) Model

We take a simplified approach to model the LPFR with plane Poiseuille flow with a biofilm at the wall. The concentration *C*_*i*_ represents the three dimensional concentration of species i:

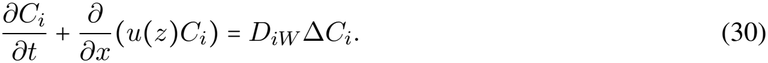

with the *x* boundary defined by Danckwerts condition

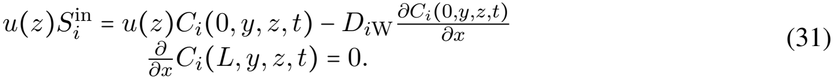

where *H* and *W* are the total channel height and width of the LPFR respectively. The *y, z* are boundary defined by

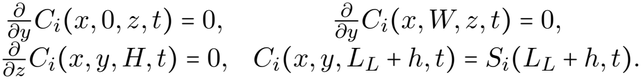

We approximate the three dimensional flow through the tank by a compartment model with averaged bulk flow by a Tank-in-Series (TIS) model. The TIS model is coupled to a fixed thickness CBL along the reactor using our previously established DL-MEC model, with one microbial species per longitudinal phase. This approach allows us to directly utilize our previous model to approximate the 1D reduced generalized axial dispersion model (ADE) with dynamic dispersion coefficient *D*_ax,*i*_ (*t*_res_) [20] with a weak reactive wall approximation to representing coupling between the bulk flow and the spatially resolved CBL and biofilm domains. While *D*_ax_(*t*_res_) governs both pre-Taylor dispersion and Taylor dispersion regimes, for ease of numerical tractability, we assume that *D*_ax,*i*_(*t*_res_) → *D*_ax,*i*_ (*t*_∞_) = *D*_ax,*i*_. While this will overestimate mixing and thus underestimate production in high flow rate regimes, it serves as a reasonable approximation to the ADE using the TIS where we limit high flow rates for LPFR. To approximate the ADE using TIS, we define the number of total tanks by

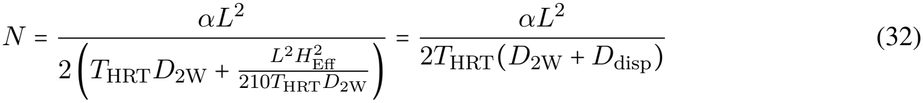

where is the length of the axial length of the tank such that 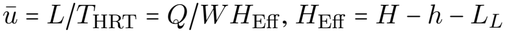 is the effective bulk flow depth, *D*_ax,2_ *D*_2W_ *D*_disp_ and *α* is the effective velocity correction term defined next. To keep the same number of tanks for each substance, all relationships are derived using our substrate methane and the number of tanks is rounded to the nearest number to ensure equal number of tanks per phase. The axial dispersion coefficient and the effective substrate velocity used in the TIS approximation is

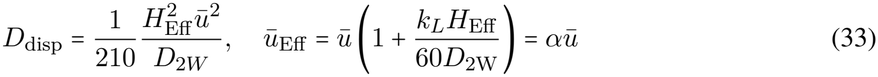

where *k*_*L*_ = *D*_2W_/*L*_*L*_ is the mass transfer coefficient.

We prescribe a CBL thickness of 1000 *µ*m. This value is not predicted by the model, but represents an optimistic engineering assumption associated with baffles or spacers that enhance transverse mixing [7]. For undisturbed laminar flow between parallel plates, the effective CBL would generally be thicker, further reducing mass transfer as detailed in Supplement S6. The selected value therefore provides one example of a best-case estimate of LPFR performance for comparison with the DL-MEC, which similarly assumes an optimistic impeller-maintained CBL thickness of 50 *µ*m.

We compare the reactors on the basis of three metrics with the following assumptions: equal biofilm volume (*h*_1_ = 55 *µ*m, *h*_2_ = 45 *µ*m), thickness per phase, and influent flow concentration 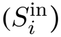 while varying over HRT or flow rate. However, the oxygen delivery systems are fundamentally different (sparge vs membrane aeration), to facilitate a true comparison, we saturate the oxygen concentration at the membrane (1280 mmol m ^−3^) in the LPFR to take full advantage of its delivery system. As before, the focus is on 2,3-BDO production as opposed to methane removal rates. We use three targeted metrics for reactor comparison with specific aims:

(i) total productivity given the same total influent (equal flow rates)

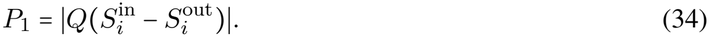

(ii) volumetric production (for equalized HRT), (iii) areal production (for equalized HRT)

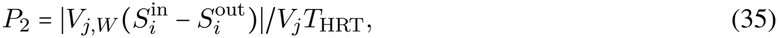

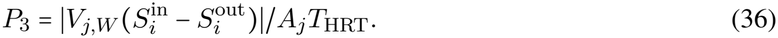

where *Q*_*j*_ *V*_*j,W*_ *T*_HRT_ where *j*∈{DL MEC, LPFR}. Using the value *µ*_*A*_ 0.4 mol_rxn_ Cmol ^−1^ d ^−1^ approximated from contour trace (Fig. 7**A**) we compare the reactors.

We upper bound our maximum flow rate used for the LPFR to align with values as detailed in Supplement S6.

Most of the comparison lies within the estimated Taylor dispersion regime where the highest flow values modestly exceed the criterion in Supplements S6 and are included as heuristic estimates. Over the range of equalized flow rates, the values represent the valid flow regimes for the LPFR rather than optimized flow regimes for the DL-MEC (Fig. 9). For the equalized HRT comparison, this is not the case. Based on the HRT values chosen, this results in vastly different flow rates for each reactor. The maximum flow rate for the DL-MEC in the equalized HRT case 10x larger than the maximum flow rate for the equalized flow rate results, while in both cases the flow rates for the LPFR remain valid (Fig. 9**B,C**).

**Figure 9.**
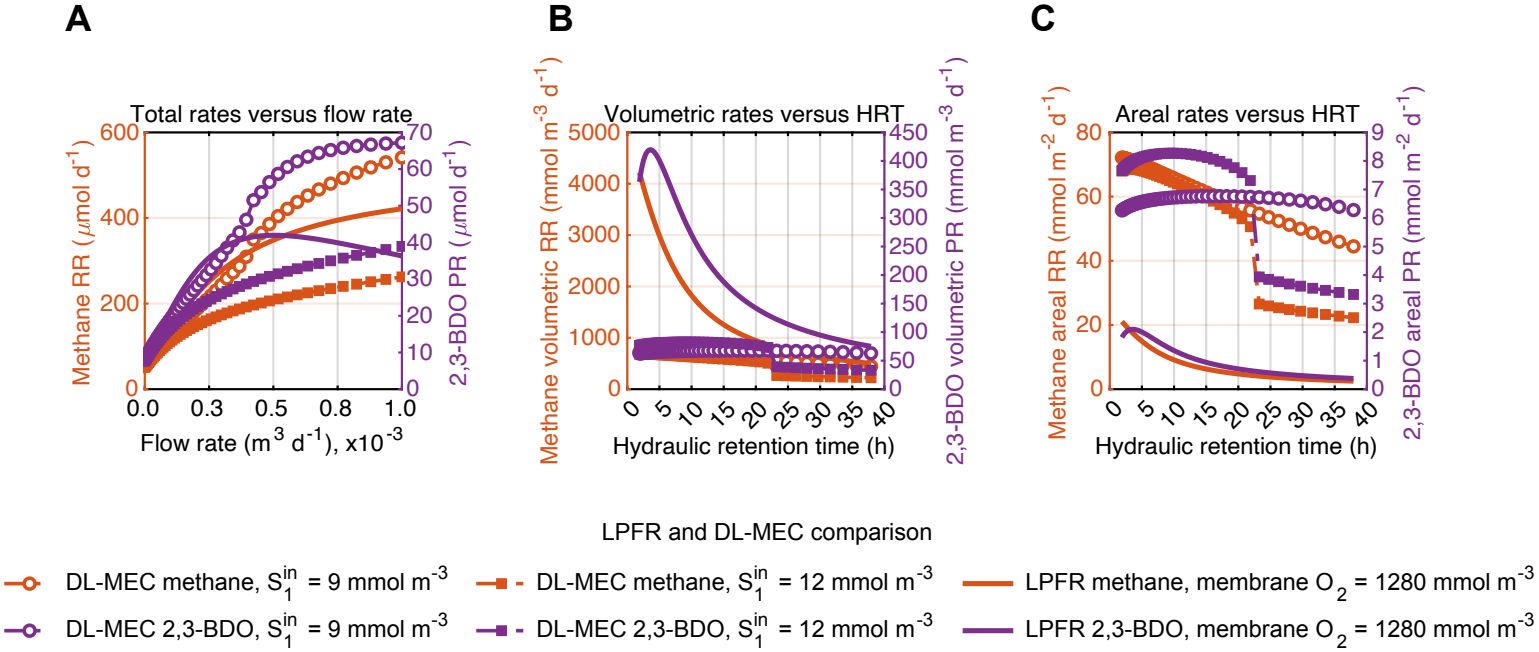
The DL-MEC achieves higher peak production despite LPFRs volumetric superiority. A.A comparison of reactors for methane removal rate and 2,3-BDO production rate for varied flow rates (34). **B**. A volumetric production comparison for varied HRT where *V*_DL −MEC_ is 10 fold larger than *V*_LPFR_ (35). **C**. A production per unit biofilm area comparison for varied HRT using where the DL-MEC biofilm surface area is half the LPFR (36). For **A**-**C**, *µ*_*A*_ = 0.4 mol_rxn_ Cmol ^−1^ d ^−1^ for both reactors. RR = Removal Rate, PR = Production Rate

The key comparative findings are as follows. For high flow rates, specifically due to mixing and minimized mass transfer limitations, the DL-MEC has a significant advantage in total production and areal production (Fig. 9**A**). At low flow rates, the DL-MEC is in the oxygen inhibited regime. As a result the LPFR out performs in total production rate.

The DL-MEC anolyte volume is 10 times bigger than the LPFR since the DL-MEC requires a larger tank for mixing, while the LPFR only needs a small channel for flow. As a result, the LPFR is superior in volumetric productivity (Fig. 9**B**). Optimal 2,3-BDO production in the LPFR occurs at an HRT of just under *T*_HRT_ ≈ 5 hours (Fig. 9**B**). When increasing HRT above 5, there is not enough acetate being produced to sustain a large production of 2,3-BDO. However, even as production declines it is still sustains higher volumetric production than the DL-MEC. This implies that as HRT decreases (and thus effective influent concentration decreases), acetate production rates are not saturated and decline accordingly.

In contrast to volumetric productivity, since LPFR utilizes longitudinal separation LPFR requires a larger and longer substratum with an oxygen membrane (Fig. 9**C**). Under low HRT, the DL-MEC outperforms the optimistically modeled LPFR by a factor of almost 5. A summarized comparison between the reactors can be found in Table 5.

**Table 5:**
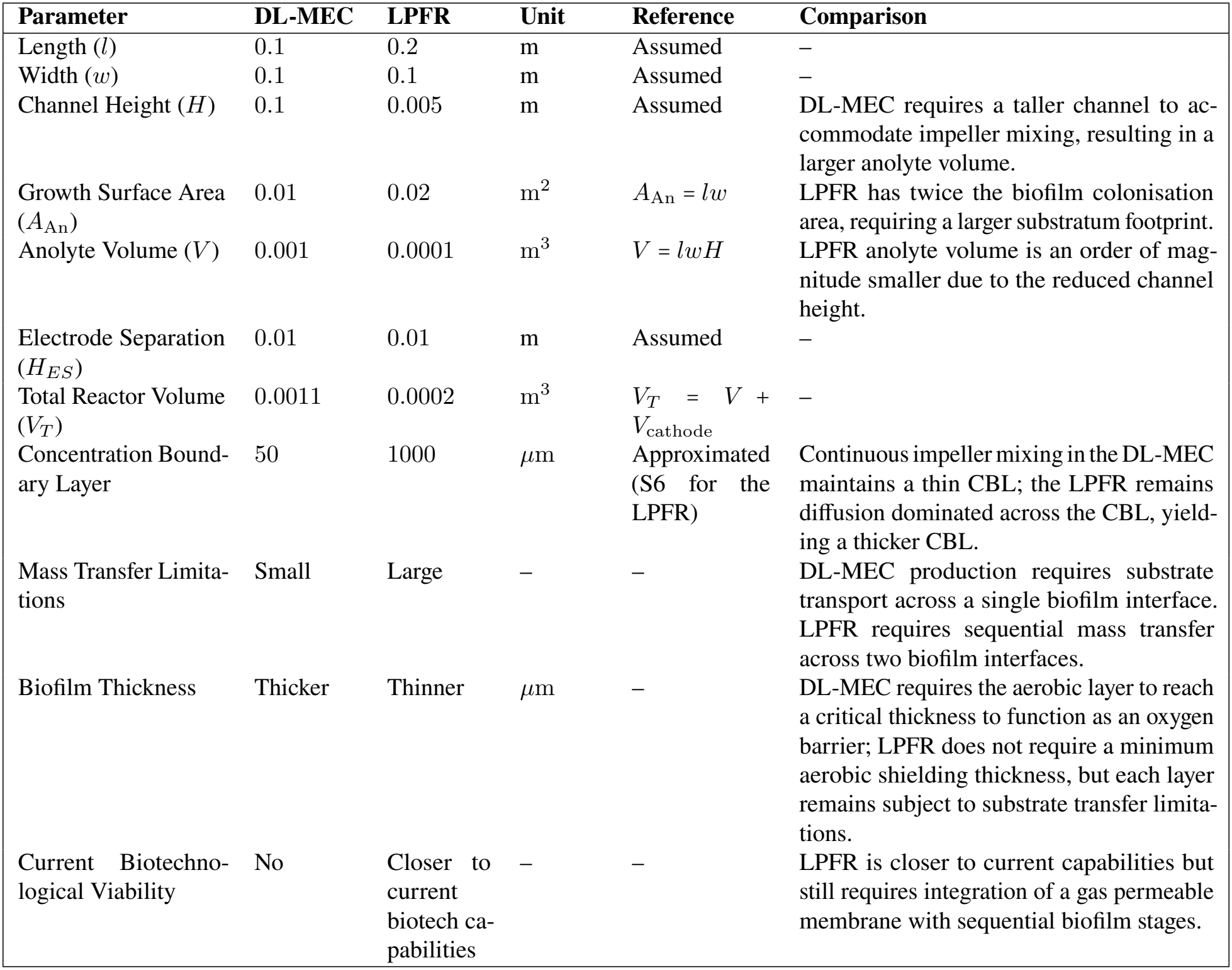
Reactor dimensions and comparison between the Double Layer Microbial Electrolysis Cell (DL-MEC) and Longitudinal Partitioned Flow Reactor (LPFR).

| **Parameter** | **DL-MEC** | **LPFR** | **Unit** | **Reference** | **Comparison** |
|---|---|---|---|---|---|
| Length (*l*) | 0.1 | 0.2 | m | Assumed | – |
| Width (*w*) | 0.1 | 0.1 | m | Assumed | – |
| Channel Height (*H*) | 0.1 | 0.005 | m | Assumed | DL-MEC requires a taller channel to ac-commodate impeller mixing, resulting in a larger anolyte volume. |
| Growth Surface Area (*A*<sub>An</sub>) | 0.01 | 0.02 | m<sup>2</sup> | *A*<sub>An</sub> = *lw* | LPFR has twice the biofilm colonisation area, requiring a larger substratum footprint. |
| Anolyte Volume (*V*) | 0.001 | 0.0001 | m<sup>3</sup> | *V* = *lwH* | LPFR anolyte volume is an order of mag-nitude smaller due to the reduced channel height. |
| Electrode Separation (*H*<sub>*ES*</sub>) | 0.01 | 0.01 | m | Assumed | – |
| Total Reactor Volume (*V*<sub>*T*</sub>) | 0.0011 | 0.0002 | m<sup>3</sup> | *V*<sub>*T*</sub> = *V* + *V*<sub>cathode</sub> | – |
| Concentration Bound-ary Layer | 50 | 1000 | µm | Approximated (S6 for the LPFR) | Continuous impeller mixing in the DL-MEC maintains a thin CBL; the LPFR remains diffusion dominated across the CBL, yield-ing a thicker CBL. |
| Mass Transfer Limita-tions | Small | Large | – | – | DL-MEC production requires substrate transport across a single biofilm interface. LPFR requires sequential mass transfer across two biofilm interfaces. |
| Biofilm Thickness | Thicker | Thinner | µm | – | DL-MEC requires the aerobic layer to reach a critical thickness to function as an oxygen barrier; LPFR does not require a minimum aerobic shielding thickness, but each layer remains subject to substrate transfer limita-tions. |
| Current Biotechno-logical Viability | No | Closer to current biotech ca-pabilities | – | – | LPFR is closer to current capabilities but still requires integration of a gas permeable membrane with sequential biofilm stages. |

## 5 Discussion

We introduced a pseudo three chamber bioreactor design, denoted DL-MEC, for the simultaneous electrode assisted methane oxidation, hydrogen production and 2,3-butanediol synthesis (Fig. 1). Here the *E. coli* have two functions, i.e., converting acetate to 2,3-BDO and to shield the *M. acetivorans* from oxygen inundation. *M. acetivorans* has the functions to convert methane into acetate which then feeds both the 2,3-BDO production and the oxygen reduction in the *E. coli* layer. We implemented a 1D in-silico model where this two-layer biofilm is in steady state conversion with constant biofilm thickness, and where the externally applied electric potential results in idealized non-limiting electrokinetics to isolate biological processes (Section 3.3). Our computational analysis under steady-state cultivation shows that such a two-stage anaerobic-aerobic biofilm redox process can work in principle, it reveals the required engineering targets to reach feasibility, and it provides deeper insights into such oxygen shielding. Three aspects require deeper consideration as discussed in the following paragraphs: operational limits of the current design, the potential for future microbial engineering, and the comparison to alternative reactor designs.

Reactor productivity depends on the delicate balance of the counter-diffusive supplies of acetate and oxygen within the aerobic biofilm (Fig. 10**A**). The counter diffusion is a feature of the multi-layer design reducing interlayer mass transfer and shortening transport distance between species specific reactions (Fig. 10**A**). Local 2,3-BDO production is greatest in that part of the biofilm where sufficiently high acetate and oxygen concentrations overlap. To obtain a working reactor at current literature values (Table 3), either a severely low oxygen influent or a currently impracticable high biofilm thickness is needed. However, a low oxygen influent comes with diminished 2,3-BDO production (Figs. 4 and 10b). For slight increases in oxygen influent, acetate production cannot meet oxygen demand to achieve at least our lowest considered production threshold: 20 *µ*mol d ^−1^. As alternative, the acetate supply to the aerobic layer could be increased by using a secondary acetate source or by biofilm engineering to increase AOM-driven acetate production (Fig. 10**B**). Given the feasibility limits of current parameter values including a biofilm thickness of *h* ≤ 150 *µ*m, the most effective path forward seems to be to co-engineer parameters - such as reaction rates, biofilm densities and enzyme turnover rates - to achieve the acetate supply needed for optimizing reactor performance.

**Figure 10.**
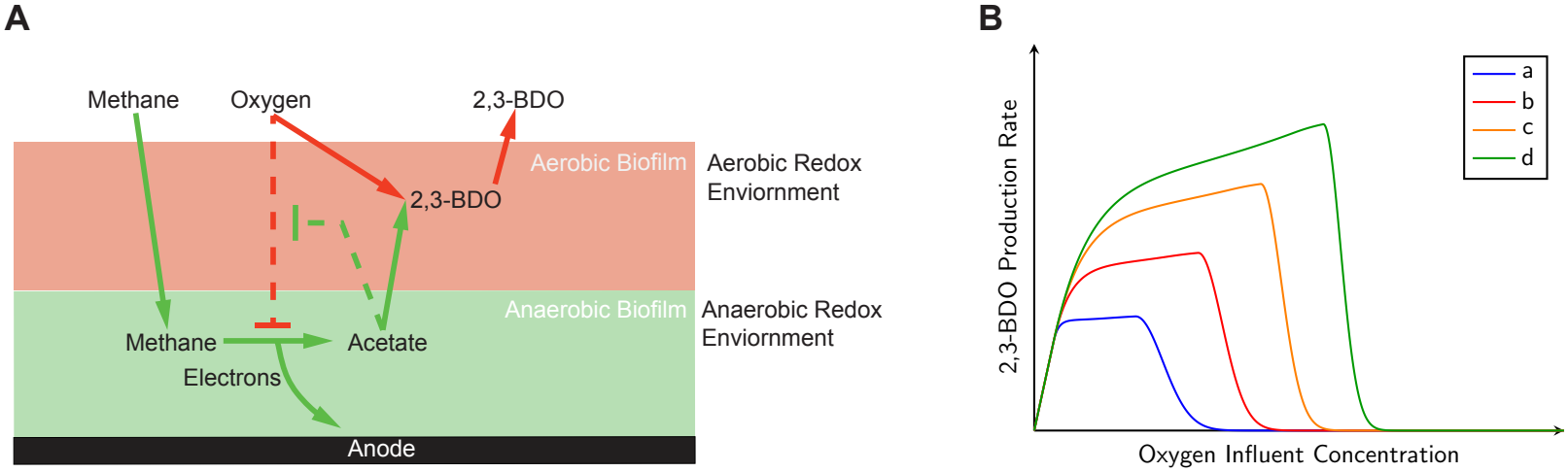
Microbial engineering influences the feedback loop between 2,3-BDO production and oxygen inhibition by impacting biofilm reaction rates. A.Multi-step reaction feedback loop between the stratified anaerobic and aerobic biofilms. **B**. Improved 2,3-BDO production as a function of oxygen influent for progressively increased reaction rates via increase of biofilm thickness, reaction rates, biofilm densities and/or enzyme turnover rates; the colored lines from a to d illustrate the increase in production and the widening of the range of usable oxygen influent when improving some combination of these parameters.

At a physically feasible biofilm thickness of both layers combined, acetate production per unit anaerobic biofilm volume can be increased by improving the maximum AOM reaction rate, increasing acetate yield, or increasing active biomass density (Fig. 10**B**). For example, at an oxygen influent 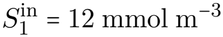 and biofilm thickness *h* 100 *µ*m, increasing the AOM reaction rate alone requires an approximately 100-fold improvement (Fig. 7**A**). By jointly engineering microbial density, the required target is lessened to about 25-fold (Fig. 7**B**,**C**). Recent, potentially complementary microbial engineering relative to established reverse methanogenesis controls [56] suggest that this target is attainable. First, a 5.2-fold increase in cultivation averaged acetate production rate was achieved by engineering a reverse methanogenic MCR variant [29]. This variant rate is used to establish our baseline estimates. Second, during co-production with ethanol, a 12-fold increase in cultivation averaged acetate production rate was achieved using antisense-RNA suppression of the native methanogenic MCR [28]. This experimental evidence supports the conclusion that remaining challenges identified by the model can plausibly be resolved with additional microbial engineering.

As the AOM-driven acetate production increases, aerobic conversion will become 2,3-BDO production limiting. Although the AOM activity is the primary constraint on feasibility, the aerobic conversion capacity is essential to obtaining even higher 2,3-BDO production (Fig. 10**B**). A three fold increase in acetate consumption was achieved [51] from the baseline established in the literature before [44]. As with the AOM rate, increased biofilm densities can additionally serve to increase aerobic production. However, since the aerobic biofilm serves as a diffusional gap between the bulk liquid and the anaerobic biofilm, excessive densification may restrict acetate and oxygen transport. Reactor operation must therefore balance increased aerobic activity against diffusion limitations and biofilm stability.

Our comparison of the DL-MEC performances to the more conventional multi-stage flow through reactor, denoted LPFR (Fig. 8), reveals advantages and challenges of either design. The LPFR utilizes horizontal stratification, mechanical shielding, oxygen saturated membrane aeration and insoluble ferric iron as the TEA. This reactor represents a more stable and robust operational baseline, as its physical separation of biofilm layers inherently protects against oxygen inhibition in the anaerobic biofilm without relying entirely on biological consumption rates. Compared to the DL-MEC with equal biomass volume, the LPFR excelled in 2,3-BDO production rate per tank volume as by design it has a smaller footprint (1/10th smaller), yet it falls short in peak production, total production at higher flow rates and areal production as it is highly mass transfer restricted (Fig. 9). Hence the LPFR provides an accessible, easier-to-handle baseline technology against which performance of real-world DL-MEC with optimized microbial kinetic parameters should be compared.

We suggest the following future experimental and theoretical work for both reactor designs: Experimental investigation is needed to (i) determine the growth and decay rates of the considered microbes, (ii) impact of pH on the growth and production rates, (iii) quantify the interplay between oxygen concentration, its ability to divert redox equivalents from the anode and degrade the anaerobic biofilm, (iv) investigate the efficiency of charge transfer mechanisms of *M. acetivorans*, and (v) analyze the microbial interactions at the interface of the aerobic-anaerobic biofilms and determine the extent to which the biofilm ratios can be controlled. Increases in microbial reaction rates can be achieved by upregulating charge transfer via multi-heme complexes, elevating the expression of crucial enzymes, increasing catalytic turnover rates, improving affinity at low methane concentrations and strengthening cellular adhesion for denser or thicker biofilms [10, 21, 67]. The next phase of modeling should center around microbial growth and electrochemical kinetics. The most direct approach is to utilize the classic Wanner-Gujer growth model [64, 65] coupled to improved biological and electrochemical reaction kinetics [1] governed by intracellular biological modeling [26, 43].

In conclusion, our analysis shows that the DL-MEC has the potential to overcome the mass transfer limitations of a more conventional longitudinal flow-through reactor by reducing the chemical transport between biofilms. The stratified biofilm layers introduce a metabolically coupled feedback loop where the bottom anaerobic layer maintains acetate production, while the top aerobic layer serves as a living membrane that simultaneously shields the oxygen from the anaerobic layer and executes the final synthesis to the value-added product using acetate (Fig. 10**A**). Other microbes with suitable consumption rates could replace *E. coli*, thereby expanding the product range from the original substrate of methane. The DL-MEC discussed here represents one use case of harnessing the metabolic feedback between interacting layered consortia for biochemical conversion. Thus, this approach serves as a design framework for microbial upgrading utilizing MECs and MxCs (Figs. 1,10) [31, 70].

## References

[1] Maliheh Abdollahi, Sara Al Sbei, Miriam A. Rosenbaum, and Falk Harnisch. The oxygen dilemma: The challenge of the anode reaction for microbial electrosynthesis from co2. Frontiers in Microbiology, Volume 13-2022, 2022. ISSN 1664-302X. doi: 10.3389/fmicb.2022.947550. URL https://www.frontiersin.org/journals/microbiology/articles/10.3389/fmicb.2022.947550.

[2] Jayaseelan Arun, PanneerSelvam SundarRajan, Kirubanandam Grace Pavithra, Packiyadoss Priyadharsini, Sivaprasad Shyam, Rangarajan Goutham, Quynh Hoang Le, and Arivalagan Pugazhendhi. New insights into microbial electrolysis cells (mec) and microbial fuel cells (mfc) for simultaneous wastewater treatment and green fuel (hydrogen) generation. Fuel, 355:129530, 2024. ISSN 0016-2361. doi: 10.1016/j.fuel.2023.129530. URL https://www.sciencedirect.com/science/article/pii/S0016236123021440.

[3] Janis E. Baeten, Christophe Walgraeve, Rafael Cesar Granja, Mark C.M. van Loosdrecht, and Eveline I.P. Volcke. Unaerated feeding alters the fate of dissolved methane during aerobic wastewater treatment. Water Research, 204:117619, 2021. ISSN 0043-1354. doi: 10.1016/j.watres.2021.117619. URL https://www.sciencedirect.com/science/article/pii/S0043135421008149.

[4] Soumyadeep Bhaduri and Manaswini Behera. From single-chamber to multi-anodic microbial fuel cells: A review. Journal of Environmental Management, 355:120465, 2024. ISSN 0301-4797. doi: 10.1016/j.jenvman.2024.120465. URL https://www.sciencedirect.com/science/article/pii/S0301479724004511.

[5] Dylan G Boucher, Emily Carroll, Zachary A Nguyen, Rohit G Jadhav, Olja Simoska, Kevin Beaver, and Shelley D Minteer. Bioelectrocatalytic synthesis: concepts and applications. Angewandte Chemie International Edition, 62(46):e202307780, 2023.

[6] Chen Cai, Ying Shi, Jianhua Guo, Gene W. Tyson, Shihu Hu, and Zhiguo Yuan. Acetate production from anaerobic oxidation of methane via intracellular storage compounds. Environmental Science & Technology, 53(13):7371–7379, 2019. doi: 10.1021/acs.est.9b00077. URL https://doi.org/10.1021/acs.est.9b00077.

[7] Simone Camarri, Alessandro Mariotti, Chiara Galletti, Elisabetta Brunazzi, Roberto Mauri, and Maria Vittoria Salvetti. An overview of flow features and mixing in micro t and arrow mixers. Industrial & Engineering Chemistry Research, 59(9):3669–3686, 2020. doi: 10.1021/acs.iecr.9b04922. URL https://doi.org/10.1021/acs.iecr.9b04922.

[8] Irene Cano, Isabel de la Torre, Jaime Muñoz-Triviño, Miguel G. Acedos, Jorge Barriuso, and José L. Garćia. Impact of oxygen availability on escherichia coli metabolism to produce 2,3-butanediol from acetate. Chemical Engineering Journal, 524:169016, 2025. ISSN 1385-8947. doi: 10.1016/j.cej.2025.169016. URL https://www.sciencedirect.com/science/article/pii/S1385894725098584.

[9] Guanghao Chen, Mark C.M. van Loosdrecht, George A. Ekama, and Damir Brdjanovic. Biological Wastewater Treatment: Principles, Modelling and Design. IWA Publishing, 07 2023. ISBN 9781789060362. doi: 10.2166/9781789060362. URL https://doi.org/10.2166/9781789060362.

[10] Stefana A. Costan, Paul M. Ryan, Honesty Kim, Charles W. Wolgemuth, and Ingmar H. Riedel-Kruse. Biophysical characterization of synthetic adhesins for predicting and tuning engineered living material properties. Matter, 7(6):2125–2143, 2024. ISSN 2590-2385. doi: 10.1016/j.matt.2024.03.019. URL https://www.sciencedirect.com/science/article/pii/S259023852400153X.

[11] P. Cristiani, M.L. Carvalho, E. Guerrini, M. Daghio, C. Santoro, and B. Li. Cathodic and anodic biofilms in single chamber microbial fuel cells. Bioelectrochemistry, 92:6–13, 2013. ISSN 1567-5394. doi: 10.1016/j.bioelechem.2013.01.005. URL https://www.sciencedirect.com/science/article/pii/S1567539413000108.

[12] Matthijs RJ Daelman, Ellen M van Voorthuizen, Udo GJM van Dongen, Eveline IP Volcke, and Mark CM van Loosdrecht. Methane emission during municipal wastewater treatment. Water research, 46(11): 3657–3670, 2012.

[13] Paolo Dessí, Laura Rovira-Alsina, Carlos Sánchez, G. Kumaravel Dinesh, Wenming Tong, Pritha Chatterjee, Michele Tedesco, Pau Farrás, Hubertus M.V. Hamelers, and Sebastiá Puig. Microbial electrosynthesis: Towards sustainable biorefineries for production of green chemicals from co2 emissions. Biotechnology Advances, 46:107675, 2021. ISSN 0734-9750. doi: 10.1016/j.biotechadv.2020.107675. URL https://www.sciencedirect.com/science/article/pii/S0734975020301774.

[14] Vaibhav Kumar Dixit and Christopher Rackauckas. Globalsensitivity. jl: Performant and parallel global sensitivity analysis with julia. Journal of Open Source Software, 7(76):4561, 2022.

[15] Qiang Fei, Michael T. Guarnieri, Ling Tao, Lieve M.L. Laurens, Nancy Dowe, and Philip T. Pienkos. Bioconversion of natural gas to liquid fuel: Opportunities and challenges. Biotechnology Advances, 32(3):596–614, 2014. ISSN 0734-9750. doi: 10.1016/j.biotechadv.2014.03.011. URL https://www.sciencedirect.com/science/article/pii/S0734975014000470.

[16] Andreas H. Foerster, Sebastian Beblawy, Frederik Golitsch, and Johannes Gescher. Electrode-assisted acetoin production in a metabolically engineered escherichia coli strain. BIOTECHNOLOGY FOR BIOFUELS, 10, MAR 14 2017. ISSN 1754-6834. doi: 10.1186/s13068-017-0745-9.

[17] Siddharth Gadkari, Sai Gu, and Jhuma Sadhukhan. Two-dimensional mathematical model of an air-cathode microbial fuel cell with graphite fiber brush anode. Journal of Power Sources, 441: 227145, 2019. ISSN 0378-7753. doi: 10.1016/j.jpowsour.2019.227145. URL https://www.sciencedirect.com/science/article/pii/S0378775319311383.

[18] Siddharth Gadkari, Mobolaji Shemfe, J. Annie Modestra, S. Venkata Mohan, and Jhuma Sadhukhan. Understanding the interdependence of operating parameters in microbial electrosynthesis: a numerical investigation. Phys. Chem. Chem. Phys., 21:10761–10772, 2019. doi: 10.1039/C9CP01288E. URL http://dx.doi.org/10.1039/C9CP01288E.

[19] Maryam Ghasemi, Sheng Chang, and Sivabal Sivaloganathan. Modeling and simulation study of simultaneous nitrification–denitrification in membrane aerated bioreactor. Journal of Membrane Science, 668:121210, 2023. ISSN 0376-7388. doi: 10.1016/j.memsci.2022.121210. URL https://www.sciencedirect.com/science/article/pii/S0376738822009553.

[20] W. N. Gill and R. Sankarasubramanian. Exact analysis of unsteady convective diffusion. Proceedings of the Royal Society of London. Series A, Mathematical and Physical Sciences, 316(1526):341–350, 2026/06/17/1970. URL http://www.jstor.org/stable/77597.

[21] David S. Glass and Ingmar H. Riedel-Kruse. A synthetic bacterial cell-cell adhesion toolbox for programming multicellular morphologies and patterns. Cell, 174(3):649–658.e16, 2026/07/06 2018. doi: 10.1016/j.cell.2018.06.041. URL https://doi.org/10.1016/j.cell.2018.06.041.

[22] Guiping Gong, Bo Wu, Linpei Liu, Jianting Li, Qili Zhu, Mingxiong He, and Guoquan Hu. Metabolic engineering using acetate as a promising building block for the production of bio-based chemicals. Eng Microbiol, 2(4):100036, Dec 2022. ISSN 2667-3703 (Electronic); 2097-4280 (Print); 2667-3703 (Linking). doi: 10.1016/j.engmic.2022.100036.

[23] Xiaodi Hao, Joseph J Heijnen, and Mark CM Van Loosdrecht. Sensitivity analysis of a biofilm model describing a one-stage completely autotrophic nitrogen removal (canon) process. Biotechnology and bioengineering, 77(3):266–277, 2002.

[24] Falk Harnisch and Dirk Holtmann, editors. Bioelectrosynthesis. Advances in biochemical engineering/biotechnology. Springer International Publishing, Cham, 2019.

[25] Chad A Haynes and Ramon Gonzalez. Rethinking biological activation of methane and conversion to liquid fuels. Nat. Chem. Biol., 10(5):331–339, May 2014.

[26] Michael A Henson and Timothy J Hanly. Dynamic flux balance analysis for synthetic microbial communities. IET Syst Biol, 8(5):214–229, Oct 2014. ISSN 1751-8849 (Print); 1751-8857 (Electronic); 1751-8849 (Linking). doi: 10.1049/iet-syb.2013.0021.

[27] R.I. Hickson, S.I. Barry, G.N. Mercer, and H.S. Sidhu. Finite difference schemes for multilayer diffusion. Mathematical and Computer Modelling, 54(1):210–220, 2011. ISSN 0895-7177. doi: 10.1016/j.mcm.2011.02.003. URL https://www.sciencedirect.com/science/article/pii/S0895717711000938.

[28] Hyeon-Ji Hwang, Ruchira Mitra, Rodolfo Garćia-Contreras, Ilke Gurgan, Vanesa Angarita-Zapata, Mrugesh Parasa, Viviana Sanchez-Torres, Ingmar H. Riedel-Kruse, and Thomas K. Wood. Enhancing reverse methanogenesis using antisense rna-mediated silencing. Chemical Engineering Journal, page 182219, 2026. ISSN 1385-8947. doi: 10.1016/j.cej.2026.182219. URL https://www.sciencedirect.com/science/article/pii/S1385894726096828.

[29] Hyeon-Ji Hwang, Mrugesh Parasa, Ruchira Mitra, Rodolfo Garćia-Contreras, Vanesa Angarita-Zapata, Ingmar H. Riedel-Kruse, and Thomas K. Wood. Engineering methyl-coenzyme m reductase for enhanced methane and carbon dioxide capture through biofilm growth. bioRxiv, 2026. doi: 10.64898/2026.06.26.734734. URL https://www.biorxiv.org/content/early/2026/06/27/2026.06.26.734734.

[30] Andrew Kato Marcus, César I Torres, and Bruce E Rittmann. Conduction-based modeling of the biofilm anode of a microbial fuel cell. Biotechnology and bioengineering, 98(6):1171–1182, 2007.

[31] Benjamin Korth and Falk Harnisch. Microbial electrochemical technologies can support sustainable energy, waste treatment, and resource recovery. Communications Sustainability, 1(1):69, 2026.

[32] Regina Kutscha and Stefan Pflügl. Microbial upgrading of acetate into value-added products—examining microbial diversity, bioenergetic constraints and metabolic engineering approaches. International Journal of Molecular Sciences, 21(22), 2020. ISSN 1422-0067. doi: 10.3390/ijms21228777. URL https://www.mdpi.com/1422-0067/21/22/8777.

[33] Jieling Li, Songwei Lin, Ting Wang, Haiping Luo, Guangli Liu, and Renduo Zhang. Spatial stratification of cathodic biofilm on nitrogen removal in the microbial fuel cell under different carbon/nitrogen ratios. Journal of Water Process Engineering, 53:103912, 2023. ISSN 2214-7144. doi: 10.1016/j.jwpe.2023.103912. URL https://www.sciencedirect.com/science/article/pii/S2214714423004312.

[34] Shen-zhou Li, Yun-feng Yan, Zi-fan Kang, Xian-zheng Yuan, Shu-guang Wang, and Zhen Yan. Synergistic integration of engineered methanogenic archaea and biochar for methane upcycling to biodegradable plastics: A circular economy approach. Environmental Science & Technology, 59(43): 23362–23373, 10 2025. ISSN 0013-936X. doi: 10.1021/acs.est.5c06115. URL https://doi.org/10.1021/acs.est.5c06115.

[35] Tao Liu, Bin Ma, Xueming Chen, Bing-Jie Ni, Yongzhen Peng, and Jianhua Guo. Evaluation of mainstream nitrogen removal by simultaneous partial nitrification, anammox and denitrification (snad) process in a granule-based reactor. Chemical Engineering Journal, 327:973–981, 2017. doi: 10.1016/j.cej.2017.06.173. URL https://www.sciencedirect.com/science/article/pii/S1385894717311245.

[36] Zilong Liu, Haoran Cui, Yuxuan Jiang, Zhiwen Wang, and Tao Chen. Mutagenesis and analysis for enhancing (2r,3r)-2,3-butanediol tolerance and production in corynebacterium glutamicum. Molecules, 31(15), 2026. ISSN 1420-3049. doi: 10.3390/molecules31152714. URL https://www.mdpi.com/1420-3049/31/15/2714.

[37] J. S. Mackie and P. Meares. The diffusion of electrolytes in a cation-exchange resin membrane. i. theoretical. Proceedings of the Royal Society of London. Series A, Mathematical and Physical Sciences, 232(1191):498–509, 2026/05/06/1955. URL http://www.jstor.org/stable/99826.

[38] Kelly J. Martin and Robert Nerenberg. The membrane biofilm reactor (mbfr) for water and wastewater treatment: Principles, applications, and recent developments. Bioresource Technology, 122:83–94, 2012. ISSN 0960-8524. doi: 10.1016/j.biortech.2012.02.110. URL https://www.sciencedirect.com/science/article/pii/S0960852412003616. Membrane Bioreactors (MBRs): State-of-Art and Future.

[39] Michael J McAnulty, Venkata G. Poosarla, Kyoung-Yeol Kim, Ricardo Jasso-Chávez, Bruce E Logan, and Thomas K Wood. Electricity from methane by reversing methanogenesis. Nature Communications, 8(1):15419, 2017.

[40] Ruchira Mitra, Yunjeong Choi, Hyeon-Ji Hwang, Rodolfo Garćia-Contreras, Vanesa AngaritaZapata, Shotaro Toya, Ilke Gurgan, Ingmar H. Riedel-Kruse, and Thomas K. Wood. Reversing methanogenesis to produce ethanol. Communications Earth & Environment, 7(1):599, 2026. doi: 10.1038/s43247-026-03601-0. URL https://doi.org/10.1038/s43247-026-03601-0.

[41] Max D Morris. Factorial sampling plans for preliminary computational experiments. Technometrics, 33 (2):161–174, 1991.

[42] Andreas Netsch, Inka Latussek, Harald Horn, and Michael Wagner. Detecting excess biofilm thick-ness in microbial electrolysis cells by real-time in-situ biofilm monitoring. Biotechnology and Bioengineering, 122(8):2049–2062, 2025. doi: 10.1002/bit.29017. URL https://analyticalsciencejournals.onlinelibrary.wiley.com/doi/abs/10.1002/bit.29017.

[43] Elad Noor, Avi Flamholz, Arren Bar-Even, Dan Davidi, Ron Milo, and Wolfram Liebermeister. The protein cost of metabolic fluxes: Prediction from enzymatic rate laws and cost minimization. PLOS Computational Biology, 12(11):1–29, 11 2016. doi: 10.1371/journal.pcbi.1005167. URL https://doi.org/10.1371/journal.pcbi.1005167.

[44] Katharina Novak, Regina Kutscha, and Stefan Pflügl. Microbial upgrading of acetate into 2, 3-butanediol and acetoin by e. coli w. Biotechnology for biofuels, 13:1–14, 2020.

[45] David Parkhurst and C. A. J. Appelo. Description of input and examples for phreeqc version 3: A computer program for speciation, batch-reaction, one-dimensional transport, and inverse geochemical calculations. Report, Reston, VA, 2013. URL https://pubs.usgs.gov/publication/tm6A43.

[46] Cristian Picioreanu, Ian M Head, Krishna P Katuri, Mark CM van Loosdrecht, and Keith Scott. A computational model for biofilm-based microbial fuel cells. Water research, 41(13):2921–2940, 2007.

[47] Cristian Picioreanu, Mark CM van Loosdrecht, Thomas P Curtis, and Keith Scott. Model based evaluation of the effect of ph and electrode geometry on microbial fuel cell performance. Bioelectrochemistry, 78 (1):8–24, 2010.

[48] R. P. Pinto, B. Srinivasan, A. Escapa, and B. Tartakovsky. Multi-population model of a microbial electrolysis cell. Environmental Science & Technology, 45(11):5039–5046, 06 2011. doi: 10.1021/es104268g. URL https://doi.org/10.1021/es104268g.

[49] R.P. Pinto, B. Tartakovsky, and B. Srinivasan. Optimizing energy productivity of microbial electrochemical cells. Journal of Process Control, 22(6):1079–1086, 2012. ISSN 0959-1524. doi:10.1016/j.jprocont.2012.04.005. URL https://www.sciencedirect.com/science/article/pii/S095915241200090X.

[50] Marissa Renardy, Louis R. Joslyn, Jess A. Millar, and Denise E. Kirschner. To sobol or not to sobol? the effects of sampling schemes in systems biology applications. Mathematical Biosciences, 337:108593, 2021. ISSN 0025-5564. doi: 10.1016/j.mbs.2021.108593. URL https://www.sciencedirect.com/science/article/pii/S0025556421000419.

[51] Luca Ricci, Xuecong Cen, Yuexuan Zu, Giacomo Antonicelli, Zhen Chen, Debora Fino, Fabrizio C Pirri, Gregory Stephanopoulos, Benjamin M Woolston, and Angela Re. Metabolic engineering of e. coli for enhanced diols production from acetate. ACS Synthetic Biology, 14(4):1204–1219, 2025.

[52] B.E. Rittmann and P.L. McCarty. Environmental Biotechnology: Principles and Applications, Second Edition. McGraw Hill LLC, 2020. ISBN 9781260441611. URL https://books.google.com/books?id=VXLwDwAAQBAJ.

[53] John R. Rumble, editor. CRC Handbook of Chemistry and Physics. CRC Press, Boca Raton, FL, 104th edition, 2023.

[54] Dipayan Samanta and Rajesh K Sani. Methane oxidation via chemical and biological methods: Challenges and solutions. Methane, 2(3):279–303, 2023.

[55] SciML and contributors. QuasiMonteCarlo.jl: Quasi-Monte Carlo samples made easy in Julia. https://github.com/SciML/QuasiMonteCarlo.jl, 2025.

[56] Valerie WC Soo, Michael J McAnulty, Arti Tripathi, Fayin Zhu, Limin Zhang, Emmanuel Hatzakis, Philip B Smith, Saumya Agrawal, Hadi Nazem-Bokaee, Saratram Gopalakrishnan, et al. Reversing methanogenesis to capture methane for liquid biofuel precursors. Microbial cell factories, 15:1–14, 2016.

[57] Markus Stöckl, Thomas Lange, Paniz Izadi, Sera Bolat, Niklas Teetz, Falk Harnisch, and Dirk Holtmann. Application of gas diffusion electrodes in bioeconomy: An update. Biotechnology and Bioengineering, 120(6):1465–1477, 2023. doi: 10.1002/bit.28383. URL https://analyticalsciencejournals.onlinelibrary.wiley.com/doi/abs/10.1002/bit.28383.

[58] Kyle A. Stone, Matthew V. Hilliard, Q. Peter He, and Jin Wang. A mini review on bioreactor configurations and gas transfer enhancements for biochemical methane conversion. Biochemical Engineering Journal, 128:83–92, 2017. ISSN 1369-703X. doi: 10.1016/j.bej.2017.09.003. URL https://www.sciencedirect.com/science/article/pii/S1369703X17302231.

[59] Boya Sun, Peijing Kuang, Yubo Cui, Yuesuo Yang, and Changhong Zheng. Simultaneous nitrification and denitrification microbial fuel cells (snd-mfc) for nitrogen removal and bioelectricity recovery: a review of performances, mechanisms, microorganisms, and applications. Journal of Environmental Management, 404:129278, 2026. ISSN 0301-4797. doi: 10.1016/j.jenvman.2026.129278. URL https://www.sciencedirect.com/science/article/pii/S0301479726007383.

[60] Marcel J. M. te Riele, Erwin D. Snijder, and Wim P. M. van Swaaij. Diffusion coefficients at infinite dilution in water and in n-methylpyrrolidone. Journal of Chemical & Engineering Data, 40(1):34–36, 01 1995. doi: 10.1021/je00017a009. URL https://doi.org/10.1021/je00017a009.

[61] Xavier Alexis Walter, Iwona Gajda, Samuel Forbes, Jonathan Winfield, John Greenman, and Ioannis Ieropoulos. Scaling-up of a novel, simplified mfc stack based on a self-stratifying urine column. Biotechnology for Biofuels, 9(1):93, 2016. doi: 10.1186/s13068-016-0504-3. URL https://doi.org/10.1186/s13068-016-0504-3.

[62] Xavier Alexis Walter, Carlo Santoro, John Greenman, and Ioannis Ieropoulos. Self-stratifying microbial fuel cell: The importance of the cathode electrode immersion height. Int J Hydrogen Energy, 44(9):4524–4532, Feb 2019. ISSN 0360-3199 (Print); 0360-3199 (Linking). doi: 10.1016/j.ijhydene.2018.07.033.

[63] Xavier Alexis Walter, Jiseon You, Jonathan Winfield, Ugnius Bajarunas, John Greenman, and Ioannis A. Ieropoulos. From the lab to the field: Self-stratifying microbial fuel cells stacks directly powering lights. Applied Energy, 277:115514, 2020. ISSN 0306-2619. doi: 10.1016/j.apenergy.2020.115514. URL https://www.sciencedirect.com/science/article/pii/S0306261920310266.

[64] Oskar Wanner and Willi Gujer. A multispecies biofilm model. Biotechnology and bioengineering, 28 (3):314–328, 1986.

[65] Oskar Wanner and Peter Reichert. Mathematical modeling of mixed-culture biofilms. Biotechnology and bioengineering, 49(2):172–184, 1996.

[66] Thomas K. Wood, Ilke Gurgan, Ethan T. Howley, and Ingmar H. Riedel-Kruse. Converting methane into electricity and higher-value chemicals at scale via anaerobic microbial fuel cells. Renewable and Sustainable Energy Reviews, 188:113749, 2023. ISSN 1364-0321. doi: 10.1016/j.rser.2023.113749. URL https://www.sciencedirect.com/science/article/pii/S1364032123006068.

[67] Philip H. Woods, Daan R. Speth, Rafael Laso-Pérez, Daniel R. Utter, S. Emil Ruff, and Victoria J. Orphan. Identification of key steps in the evolution of anaerobic methanotrophy in ¡i¿candidatus¡/i¿ methanovorans (anme-3) archaea. Science Advances, 11(25):eadq5232, 2025. doi: 10.1126/sciadv.adq5232. URL https://www.science.org/doi/abs/10.1126/sciadv.adq5232.

[68] Ryota Yamasaki, Toshinari Maeda, and Thomas K. Wood. Electron carriers increase electricity production in methane microbial fuel cells that reverse methanogenesis. Biotechnology for Biofuels, 11(1):211, 2018. doi: 10.1186/s13068-018-1208-7. URL https://doi.org/10.1186/s13068-018-1208-7.

[69] Jiawei Yang, Shaoan Cheng, Peng Li, Haobin Huang, and Kefa Cen. Sensitivity to oxygen in microbial electrochemical systems biofilms. iScience, 13:163–172, Mar 2019. ISSN 2589-0042 (Electronic); 2589-0042 (Linking). doi: 10.1016/j.isci.2019.01.022.

[70] Xueqin Zhang, Zhiguo Yuan, and Shihu Hu. Current trends in electromicrobiology of methane oxidation. Trends in Microbiology, 33(7):782–795, 2026/08/08 2025. doi: 10.1016/j.tim.2025.02.012. URL https://doi.org/10.1016/j.tim.2025.02.012.

