## Supplements for "Modeling Aerobic-Anaerobic Microbial Electrolysis Cells for Multi-step Methane Conversion"

November 26, 2026

#### 1 Detailed Explanation of Design Choices and Approximated Conductivities/Resistivity

| **Component** ($i$) | **Conductivity** $\sigma_i$ ($\mathrm{S\,m^{-1}}$) | **Length** $L_i$ **(m)** |
|---|---|---|
| Biofilm – Redox Mediated (BF) | $7.5 \times 10^{-4}$ | $h_1 = 50 \times 10^{-6}$, $h_2 = 50 \times 10^{-6}$ |
| Biofilm – Nanowires (BF) | $1.25$ | $h_1 = 50 \times 10^{-6}$, $h_2 = 50 \times 10^{-6}$ |
| Electrode Separation (ES) | $5$ | $0.01$ |
| Anode, ionic (An,ion) | $0.68$ | $0.005$ |
| Anode, electrical (An,elec) | $1.25 \times 10^{4}$ | $0.005$ |
| Proton Exchange Membrane (PEM) | $10$ | $100 \times 10^{-6}$ |

Table 1: **Conductivities and lengths of reactor components.** All references and explanations are contained in the discussion.

Ohmic overpotential is defined by

$$\eta_{\mathrm{Ohm}} = \eta_{\mathrm{Ohm,BF}} + \eta_{\mathrm{Ohm,ES}} + \eta_{\mathrm{Ohm,An}} + \eta_{\mathrm{Ohm,PEM}} \tag{1}$$

where $i$ is the current density, $\eta_i$ represent the ohmic overpotential of the biofilm, catholyte, anode, and proton exchange membrane respectively. It can be further decomposed into

$$= i\left(\frac{h}{\sigma_{\mathrm{BF}}} + \frac{H_{\mathrm{ES}}}{\sigma_{\mathrm{ES}}} + \frac{L_{\mathrm{An}}}{\sigma_{\mathrm{An,ion}}} + \frac{L_{\mathrm{An}}}{\sigma_{\mathrm{An,elec}}} + \frac{L_{\mathrm{PEM}}}{\sigma_{\mathrm{PEM}}}\right) \tag{2}$$

where $L_i$, is the electrode separation (ES), anode and PEM thickness and distance $\sigma_i$ is the conductivity of the biofilm, the catholyte solution between electrodes (ES), the ionic and electrical conductivity of the porous anode and the conductivity of the PEM respectively. The conductivity of the biofilm directly relates to how microbes perform EET. For MxCs there are two main charge transfer mechanisms for *M. acetivorans*: nanowires and diffusible redox mediators (Boucher et al., 2023). For direct electron transfer via nanowires or pili we estimate the conductivity of nanowires is $\sigma_{\mathrm{n}} = 250\ \mathrm{S/m}$, if we assume that nanowires make $1\%$ of the anaerobic biofilm volume then the conductance $\sigma_{\mathrm{BF}} = \left(0.01\sigma_{\mathrm{n}}\frac{h_1}{h}\right)$ (Wood et al., 2023). However, the anodic facing, anaerobic microorganisms of consideration *M. acetivorans* do not possess nanowires. With the current biotechnology, diffusible redox mediators could be employed however, they would incur significant current limitations.

The conductivity of the biofilm arises from the diffusive flux of the charge carriers. Using Faraday's Law of Electrolysis and the steady state redox cycling arising via the Nernst-Planck equation for mediator flux $J_i$ ($J_{ox}$+$J_{red}$ = 0), electron transport via diffusion can be used to solve for electrical conductivity. Equating $i = F \sum_i n_i J_i$ via Ohm's law for a constant material,the conductivity is found using the Nernst-Einstein relation

$$\sigma_{\mathrm{BF}} = \frac{F^2}{R_g T}(n_{ox}^2 D_{ox} S_{ox} + n_{red}^2 D_{red} S_{red}). \tag{3}$$

where $n_i$ is the charge of the $ith$ species, $R_g$ is the universal gas constant and $T$ is the temperature and $S$ are the respective concentrations of the redox mediators. Using thionine as a representative mediator ($D_{\mathrm{ox}} = D_{\mathrm{red}} = D \approx 1.7 \times 10^{-9} \mathrm{m}^2/\mathrm{s}$, $n_{\mathrm{ox}} = n_{\mathrm{red}} = 1$ $S_{ox} = S_{red} = 0.5\,\mathrm{mol/m}^3$), we estimate a conductivity of $\sigma_{BF} \approx 7.5 \times 10^{-4} S/\mathrm{m}$, which is significantly lower than the nanowire case ($\sigma_{BF} \approx 1.25\,S/\mathrm{m}$ with $h_1/h = 0.5$). The mediator concentrations were estimated based on (Picioreanu et al., 2010) initial conditions, we use an arithmetic mean of these values as a representative concentration.

For each 4 mol of methane oxidized, 11 protons are produced in the oxidation half reaction with two corresponding protons consumed in the reudction half reaction. These protons must diffuse from the biofilm through the porous anode, PEM and electrode gap (catholyte) to the cathode. Proton transport is fastest via a combination of proton hopping and travel via protonated electrolytes. As a result, we choose a highly conductive saline catholyte solution with conductivity $\sigma_{\mathrm{ES,Sal}}$ = 5 S/m (Wood et al., 2023). It is expected there to be ample available protons for hydrogen gas reduction found in the catholyte so cathodic concentration overpotential is neglected. If protons experience significant resistance diffusing through the anode, in addition to the significant ohmic resistances incurred, the accumulation of the remaining protons in the anodic chamber lowers the pH harming the methanogens and decreasing current. This is anodic concentration overpotential. To mitigate the potential of this event occurring, we utilize a phosphate buffer solution (PBS) to stabilize the pH within the biofilm matrix. Since the primary channel for ionic transport and conductivity is in the electrode gap, the biofilm matrix can be optimized for biological viability. As a result, utilizing PBS without incurring larger cell resistance loss's due to lower conductivity solution of PBS ($\sigma_{\mathrm{ES,PBS}} = 1.45$ S/m(Wood et al., 2023)) is advantageous for our setup. PBS has a smaller conductivity than the saline solution, however, since the biofilm is not between the electrodes the solution of the anolyte is not a significant factor in cell resistance.

For our macroscopic approximations for the conductivities of the bioanode, the standard Bruggman relation is used ($\sigma_{\mathrm{eff}} = \sigma\epsilon^{1.5}$) as is widely used across reactor scale MxC modeling (Picioreanu et al., 2007). The porous bioanode is divided into liquid portion $\epsilon_{\mathrm{An}}$ (i.e. the anolyte) and solid portion $1 - \epsilon_{\mathrm{An}}$ (physical anode metal). Protons move through the anode via the anolyte, thus the calculated effective ionic conductivity $\sigma_{\mathrm{An,ion}} = \sigma_{\mathrm{PBS}}\epsilon_{\mathrm{An}}^{1.5}$. Electrons move through to the cathode via the conductive anode the calculated effective electrical conductivity is $\sigma_{\mathrm{An,elec}} = \sigma_{\mathrm{material}}(1 - \epsilon_{\mathrm{An}})^{1.5}$. Assuming carbon materials, $\sigma_{\mathrm{material}} = \sigma_{\mathrm{carbon}} \sim 10^4 - 10^5$ S/m (Baudler et al., 2015). Choosing the porosity value $\epsilon_{\mathrm{An}} = 0.6$ (Picioreanu et al., 2010) and utilizing the PBS conductivity for ionic conductivity, the effective conductivity is approximately $\sigma_{\mathrm{An,ion}} = 0.68$ S/m. For the final conductivity value, we consider a Nafion PEM where the hydrated regions of the PEM are highly conductive ($\sigma_{\mathrm{PEM}} = 10$ S/m(Springer et al., 1991),(Zawodzinski et al., 1993)). Since the conductivity is considerably large and the thickness of the PEM is small, it will not provide significant resistance to ionic transport.

### 2 Shift in Biofilm Ratio for Optimal Production rates

For $\mu_A \approx \mu_B \approx 0.5$ Cmol$^{-1}$ day$^{-1}$, one can observe that high productivity occurs along the tipping point where acetate production is favored. As the anaerobic biofilm gets thicker, methane is more utilized and production of 2,3-BDO is increased towards $\mu_B$ (Fig 1).

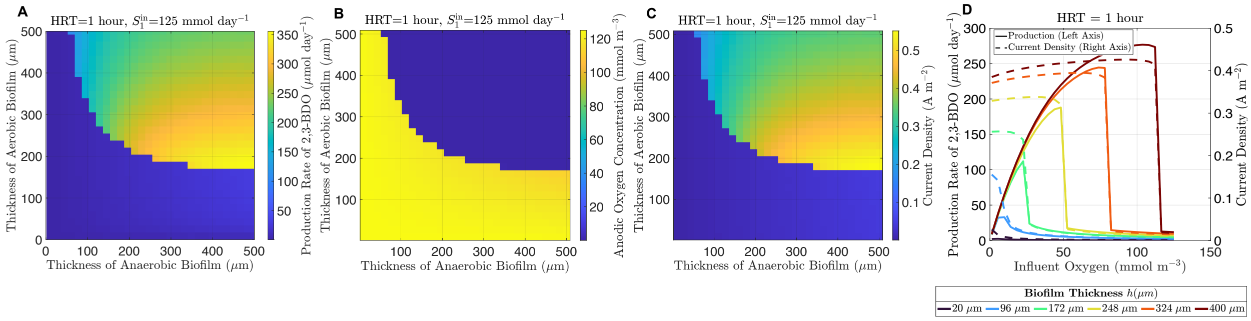


Figure 1: **The production rate of 2,3-BDO is maximized along the tipping point boundary where the highest acetate supply meets fixed oxygen demand.** For **A**-**C** $T_{\text{HRT}}$ = 1 hour for $S_1$ = 12 mmol day$^{-1}$ **A.** Production rate of 2,3-BDO , **B.** Oxygen concentration at the anode, **C.** Non potential limiting current density **D.** Production rate of 2,3-BDO (left axis) and current density (right axis) for varied oxygen influent concentrations $S_1^{\text{in}} \approx 1 - 100$ mmol m$^{-3}$ and varied biofilm thicknesses where $h_1 = h_2$ and $h \approx 20 - 1000$ $\mu$m.

### 3 Further Sensitivity Analysis

#### 3.1 Morris Sensitivity Analysis

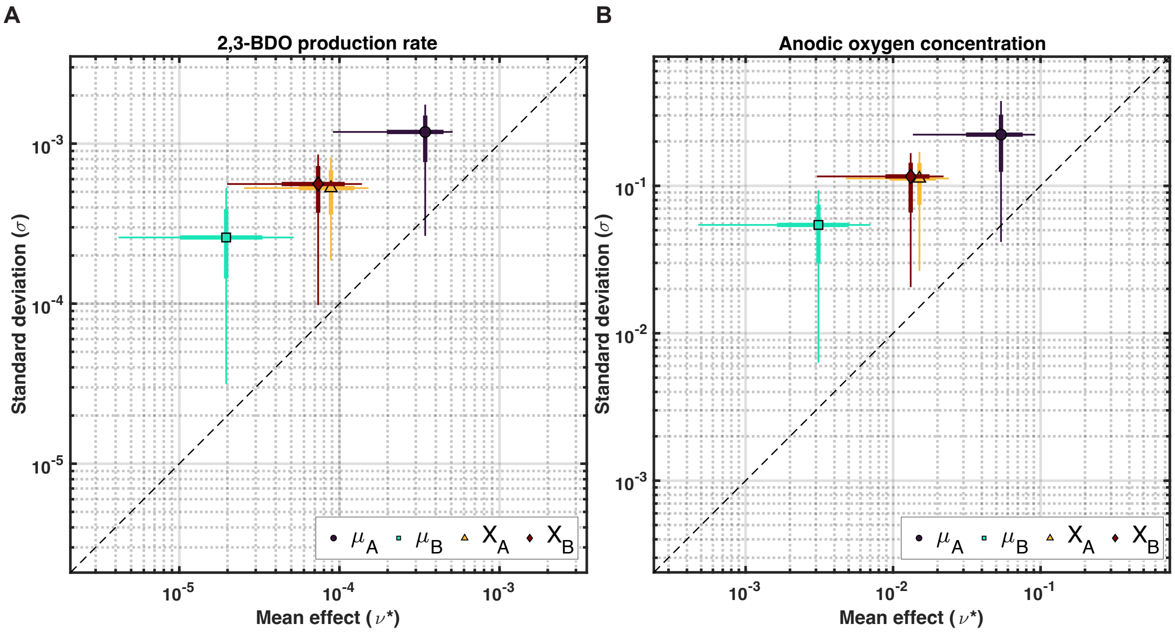


Figure 2: **Morris Sensitivity screening reveals that $\mu_A$ has the highest mean effect i.e. highest sensitivity, for increasing production and minimizing oxygen in the anode across the viable region, while the standard deviation shows that each of the parameters are influenced greatly by the other parameters. A.** the average mean effect, and standard deviation for each parameter in our screening on production of 2,3-BDO, with an interquartile range (thick line) between [25%-75%] and extended range to (thin line) [5%-95%], **B.** the average mean effect, and standard deviation for each parameter in our screening on minimizing oxygen in the anode, with an interquartile range (thick line) between [25%-75%] and extended range to (thin line) [5%-95%].

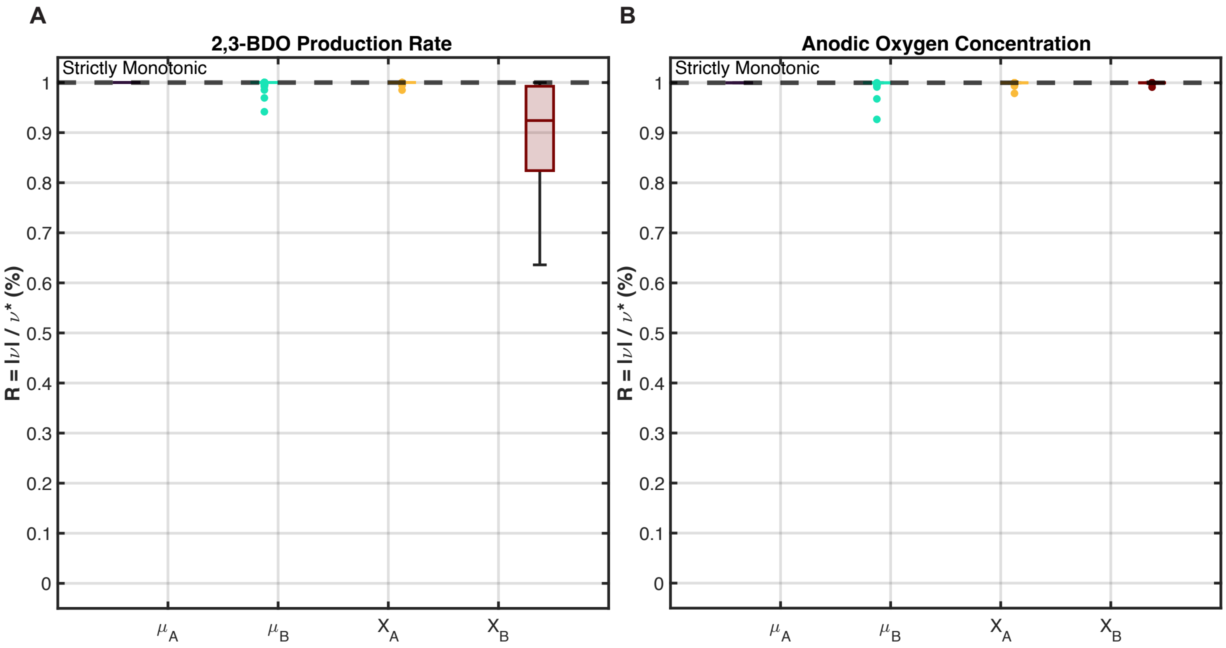


Figure 3: **All parameters are monotonic with respect to each objective function except $X_B$ for increasing productivity R ≈ 1** The monotonicity index (R) representing the monotonicity of the parameter with respect to an objective function where the box represents the interquartile range [25%-75%], and whiskers represents the extended quartile range [5%-95%]. **A.** Monotonicity index with the production rate as the objective function, **B.** monotonicity index with the objective function as the anodic oxygen concentration.

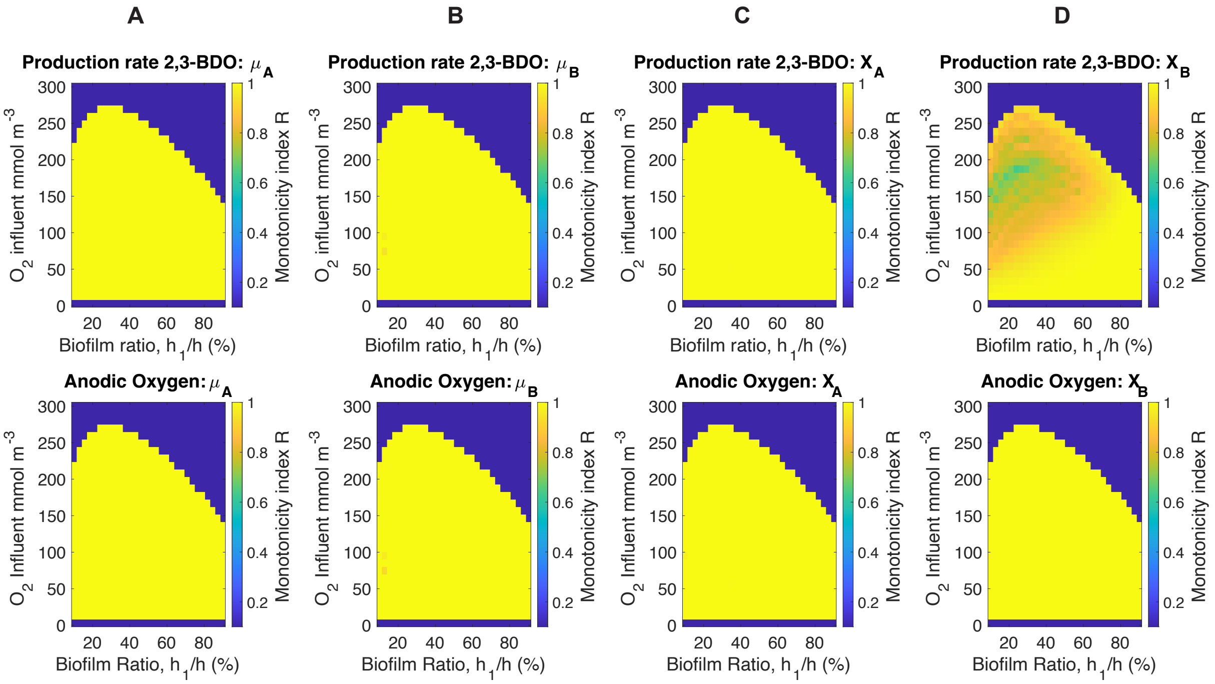


Figure 4: **The monotonicity index ($R$) over the $(h_1, O_2)$, grid for each parameter with for $h = 100\ \mu\text{m}$.** The monotonocity index of the 2,3-BDO production rate (first row) and of the anodic oxygen concentration (second row) of each parameter is shown **A.** $\mu_A$, **B.** $\mu_B$, **C.** $X_A$, **D.** $X_B$.

The qualitative effect of the parameters on the objective functions $PR_4$ and $S_4^{\text{Anode}}$ are considered using the monotonicity index. This metric quantifies the directional consistency of a parameter with respect to the objective function. As a result, learning that the objective function is monotonic or non-monotonic with respect to the given parameter (Dixit and Rackauckas, 2022), (Morris, 1991)

$$R_p^{ij} = \frac{|\nu_{ij}|}{\nu_{ij}^*} \tag{4}$$

where $p \in \{\mu_A,\ \mu_B,\ X_A,\ X_B\}$, $\nu_{ij}$ is the global mean slope of parameter $p$ across p's domain at the ith oxygen influent concentration, jth biofilm ratio concentration and $\nu_{ij}^*$ global mean of the absolute values of the slope across the parameters' domain. To summarize the behavior, as noted in the main section, $\mu_A$, $\mu_B$ and $X_A$ are strictly monotonic with respect to the production rate and Anodic oxygen concentration (Fig. 3). $X_B$ is slightly non monotonic (especially for higher oxygen influent) for increased productivity while is is strictly monotonic for minimizing anodic oxygen concentration. While the densities, cannot push the system into feasibility, thus decreasing its Morris mean, they severely decrease the distance to the tipping, alleviating the amount of biological engineering by focusing on optimal growth and operating conditions (Fig 2). We expand upon the behavior below.

The Morris plots calculate the variability and mean effect of each parameter by taking global derivatives with respect to individual paths along random trajectories defined by the algorithm. While certain parameters achieve a higher mean effect, all parameters exhibit high volatility. This implies that the ability of any single parameter to shift the system across the tipping region depends strongly on the values of the remaining parameters. For both anodic oxygen and production rate it is evident that the anaerobic reaction rate is most effective, while the densities will never mount the tipping points themselves, they severely decrease the proximity to the tipping points.

At elevated oxygen concentrations, elementary effects of both signs establish that productivity is nonmonotonic with respect to $X_B$. Because logarithmic parameter sampling provides relatively limited resolution in the upper range of $X_B$, the estimated magnitude of the monotonicity index does not fully stablize in this region. Therefore, while our stablization results confirm that $R < 1$, its precise value should be interpreted cautiously. This is evidenced in the Morris stabilization plot for nonmonotonic grid points for $X_B$.

Lastly, it is important to note that although the failures here are sufficiently small, the Morris method (Dixit and Rackauckas, 2022) requires values for all trajectories. Thus to suffice, for each point that failed in a Morris trajectory we found its 3 closest neighbor over all sampled points and calculated its inverse weighted average to supply a value. This happened most frequently as $h_1/h \to 10\%$. This, in addition to log sampling which sufficiently samples the lower regions of our bounds but undersample the higher regions shows why some $R < 1$ for $\mu_B$.

### 4 Convergence, Stabilization and Numerical Schemes

For our numerical schemes we use a number of different methods that required specified sampling points. Those specified points can be found in the following table

| | Morris | | Sobol | |
|---|---|---|---|---|
| | Number of Trajectories | Sampled Regions | Sequence Length | Scrambler |
| | 2500* | 10 | 1024 | None |
| Grid | $h \times S_1 = 30 \times 30$ | | | |
| Number of Parameters Per Grid Point | 2 or 4 (simulation dependent) | | | |
| | Finite Difference Method | | | |
| Mesh Size | 100 (200 for grid search) | | | |
| Sample Failures | No grid point exceeds $50/25000 = 0.2\%$ failures | | 0 | |

Table 2: **Numerical methods and sensitivity analysis parameters**.*The Julia GSA package (Dixit and Rackauckas, 2022) uses trajectories with two times (number of params +1) resulting in 25000 samples per grid point with the highest number occurring for small anaerobic biofilms. This number is used to sample the entire space, when sampling small pockets that are fully monotonic we use a smaller number as evidenced in Morris stabilization plots.

In order for our results to be valid, we must show convergence of our numerical scheme. To do so we utilize three different tests. Note, this is performed with $\mu_A = 0.4$ and $\mu_B = 1.53$.

1. (Interface accuracy) To verify proper interface handling and and numerical consistency of diffusion across the grid, we compare our finite difference numerical method to a piecewise linear diffusion equation with 3 phases dependent upon the diffusion coefficients. This replicates what occurs for each given species at the numerical interface and across phases. Since we choose a piecewise linear true solution, our method resolves the solution below tolerance's of the ODE15s numerical method (Fig. 5**A**).
2. (Mesh Convergence) To validate time dependent method of lines numerical scheme, we utilize a two phase solution to show convergence of our numerical scheme at steady state as the number of discretized points increases compared to a single phase that acts as a CBL and a second phase as the biofilm with a first order reaction and one chemical species. While we utilize a second order finite difference scheme on the interior, due to the non uniform grid at the interfaces, globally we achieve first order convergence (Fig. 5**B**).

3. (Conservation of Carbon Moles) To confirm carbon mole conservation as a function of time we consider batch cultivation full model. As we note, we do not conserve water and protons numerically but focus on carbon conservation. We replicate that here. It is important to note, that the small deviations seen, which are very small in nature as can be seen occur at the initialization and the introduction of the oxygen, as noted. Rather than a solver error, this is an error in how we process the data utilizing the trapezoidal method during a steep moving front. As time goes on an the moving front smooths out, the trapezoidal rule recovers the initial carbon amount. Carbon moles are calculated from the total mol at each time step

$$M_i(t) = V_W S_i(t) + A_{\mathrm{An}} \int_0^{h+L_L} S_i(z,t) dz$$

to total Carbon mol (the conserved quantity)

$$C(t) = \sum_i^M n_i M_i(t) \tag{5}$$

where $n_i$ is the number of carbon moles per mole of substance (e.g. 1 mole of methane contains 1 carbon mole) $M_i(0) = V_W S_i(0)$. Therefore, carbon moles is always conserved (Fig. 6).

4. (Sobol Stabilization) This confirms that the ratio metric converges to a fixed value as we increase the length of the Sobol sequence (Fig 7). This is first found for one representative point up to a sequence of $N$ = 2048 with no direct sub sampling. Then across the 30x30 grid we found how much the percentage points change as we increase the Sobol sequence using nested sequences that are all valid Sobol sequences themselves.

5. (Morris Stabilization)To demonstrate reliability of Morris analysis we use three figures. We choose a grid level size of 10, to evaluate the global jumps taken to surmount the tipping point.

   (a) First, we confirm that the estimated monotonicity index stabilizes as the number of Morris trajectories increases at a representative point in the region where most of the subsequent analysis is performed (Fig. 8). Monotonic cases generally stabilize with fewer trajectories. We use the grid point ($S_1$ = 50 $\mathrm{mmol\,m^{-3}}$, $h_1$ = 60 $\mu$m, and $h_2$ = 40 $\mu$m) and generate ten independent Morris designs at each trajectory count. The mean monotonicity index and its confidence interval are calculated across these independent replicates. The confidence intervals generally narrow as the number of trajectories increases, supporting stabilization of the estimated monotonicity index at this representative point.

   (b) We next evaluate sampling stability over a coarse $5 \times 5$ grid spanning the same oxygen and biofilm ratio as the full $30 \times 30$ grid (Fig. 9). At each grid point, we use five independent Morris designs and calculate the absolute difference between successive estimates of the monotonicity index,

$$\Delta R_j = \left| R(r_j) - R(r_{j-1}) \right|, \tag{6}$$

   as the number of trajectories increases from 500 to 3000. The plotted lines show the mean $\Delta R$ averaged over the replicate across the relevant grid points, while the shaded regions show the minimum-to-maximum range across those grid points. Thus, the upper edge of each shaded region represents the least-stable grid point for that parameter.

   In panels **A,B**, we consider the interior grid points with $h_1 > 10$ $\mu$m. For these points, the successive differences are either small from the beginning or decrease as the number of trajectories increases. The most noticeable remaining difference occurs for $X_B$, which is also the parameter that exhibits nonmonotonic effects.

   In panels **C,D**, we consider only the boundary points with $h_1 = 10$ $\mu$m. Greater variability is observed for $\mu_B$ and $X_B$ than in the interior grid, whereas the differences for $\mu_A$ and $X_A$ are negligible. We attribute the greater uncertainty at this boundary to a combination of (i) lower sampling resolution in the upper portion of the log-spaced parameter domain, (ii) variability among the independently generated replicate Morris designs, and (iii) increased model and numerical sensitivity at the smallest anaerobic layer thickness, where failed model evaluations replaced with inverse-distance weighted values.

(c) Finally, at each point in the representative $5 \times 5$ grid at which the final mean monotonicity index for $X_B$ across all 5 independent replicates satisfies $R < 0.95$ (Fig. 10). Each panel shows the mean value of $R$ and its 95% confidence interval as the number of trajectories increases. At these grid points, the confidence interval narrows and the estimated mean remains below the monotonicity threshold as the trajectory count increases. These results show that the qualitative nonmonotonic effectd of $X_B$ are stable at the specified oxygen and biofilm ratio conditions, yet with uncertainty remaining in the precise numerical value of $R$.

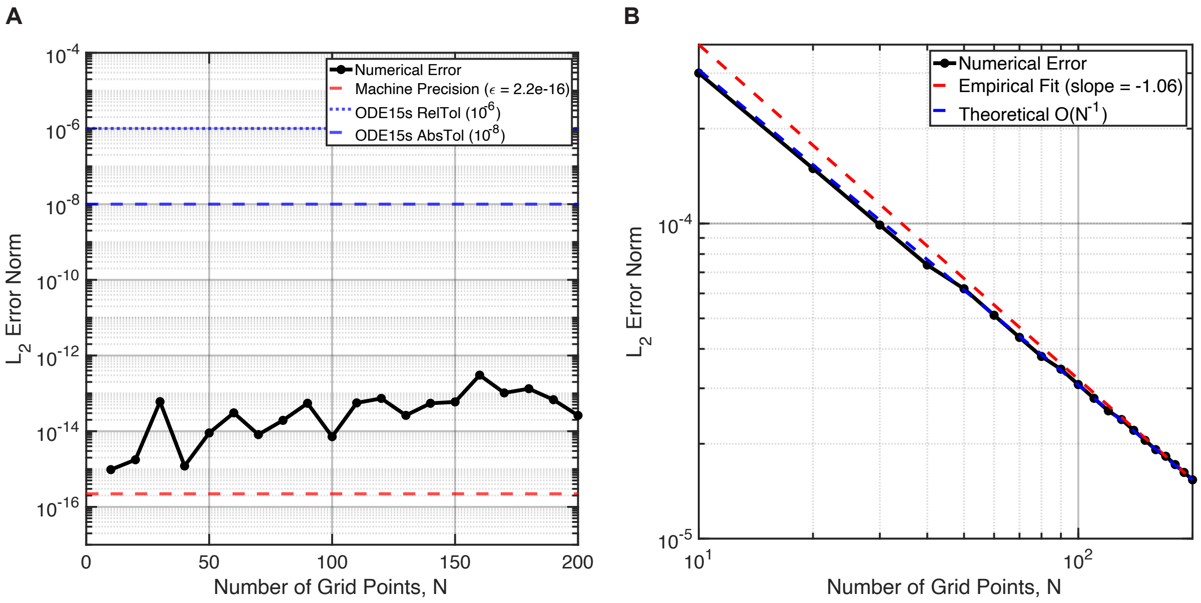


Figure 5: **The numerical method used to simulate the Double Reactor has global first order convergence while interface points matched exactly. A.** L2 error to resolve a steady state piecewise linear three phase solution to the diffusion equation exactly, **B.** L2 error to resolve a steady state two phase solution to a first order reaction diffusion equation.

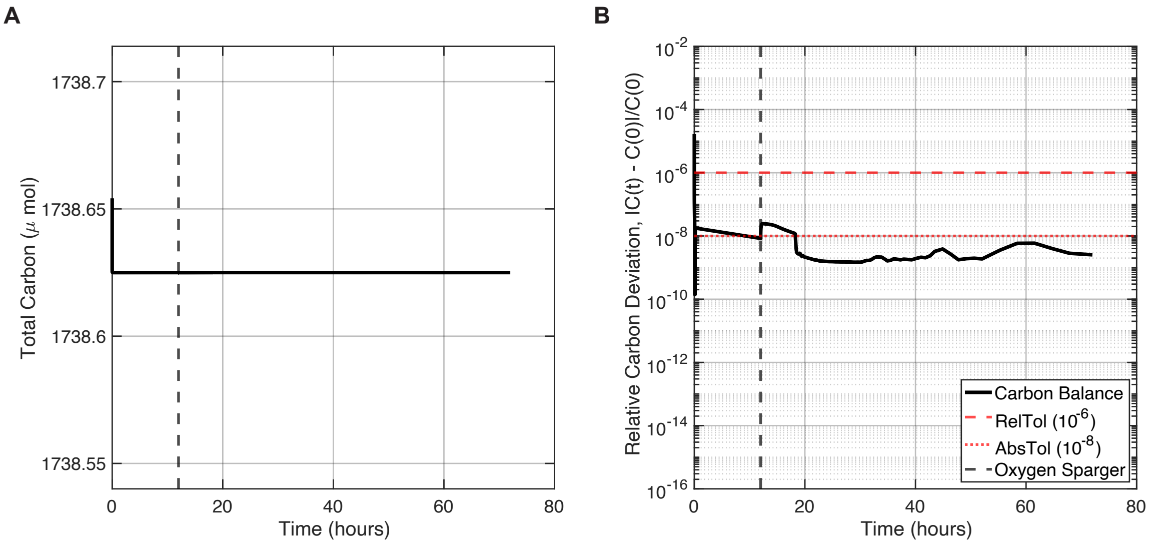


Figure 6: **Conservation of carbon mass is achieved across time**. **A.** Conservation of mass is achieved for total carbon with two deviations due to a fast moving front and data collection using the trapezoidal rule, **B.** The deviation of the solution over time compared to the initial solution disregarding the trapezoidal rule error.

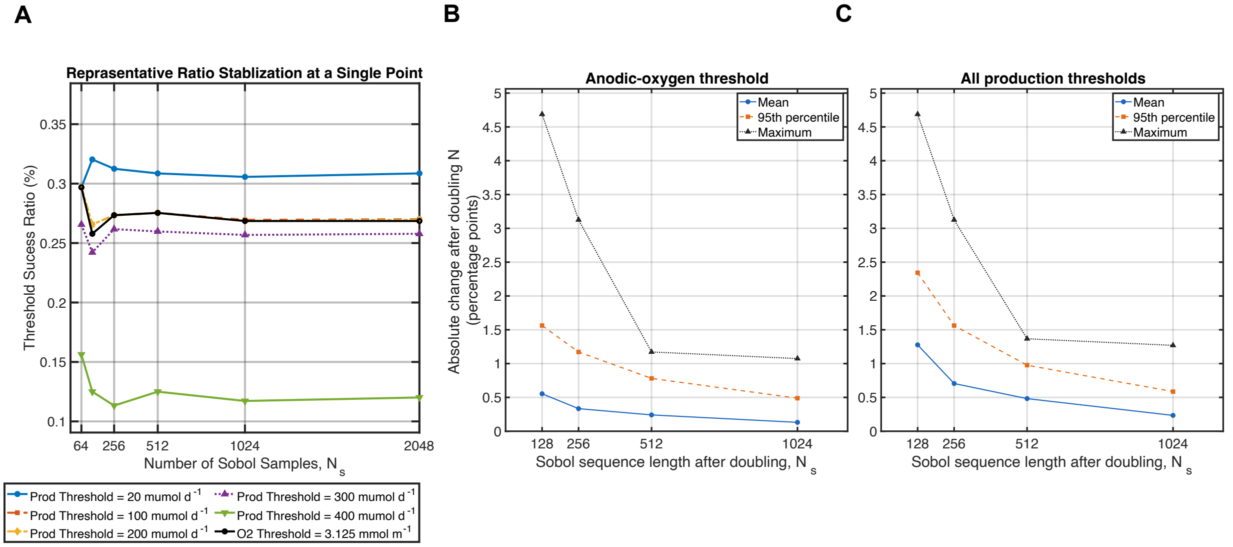


Figure 7: **Stabilization of the threshold successive ratio across the whole sampled grid using Sobol sequences.** **A**. Stabilization of the formula Ratio = $\frac{N_S}{N_{\text{Tot}}}$ as $N$ grows $h_1$ = 40 $\mu$mol, $h_2$ = 60 $\mu$mol, $S_1$ = 50 mmol m$^{-3}$, **B**. On the 30x30 oxygen influent biofilm grid, a successive difference in the threshold success ratio between Sobol prefix subsamples for the anodic oxygen threshold decreases, **C** On the 30x30 oxygen influent biofilm grid, across all production thresholds, successive difference in the threshold sucess ratio between Sobol prefix subsamples decreases.

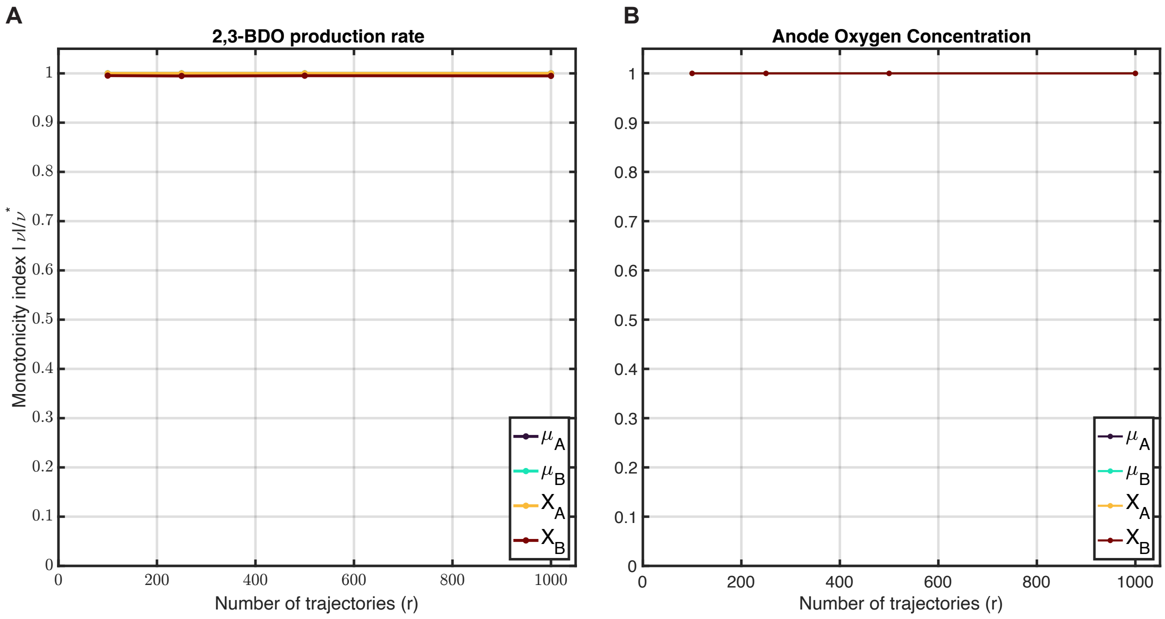


Figure 8: **Stablization of Monotonicity Index via the Morris method.** Convergence of the monotonicity index as the number of trajectories grows $h_1$ = 60 $\mu$mol, $h_2$ = 40 $\mu$mol, $S_1$ = 50 mmol m$^{-3}$.

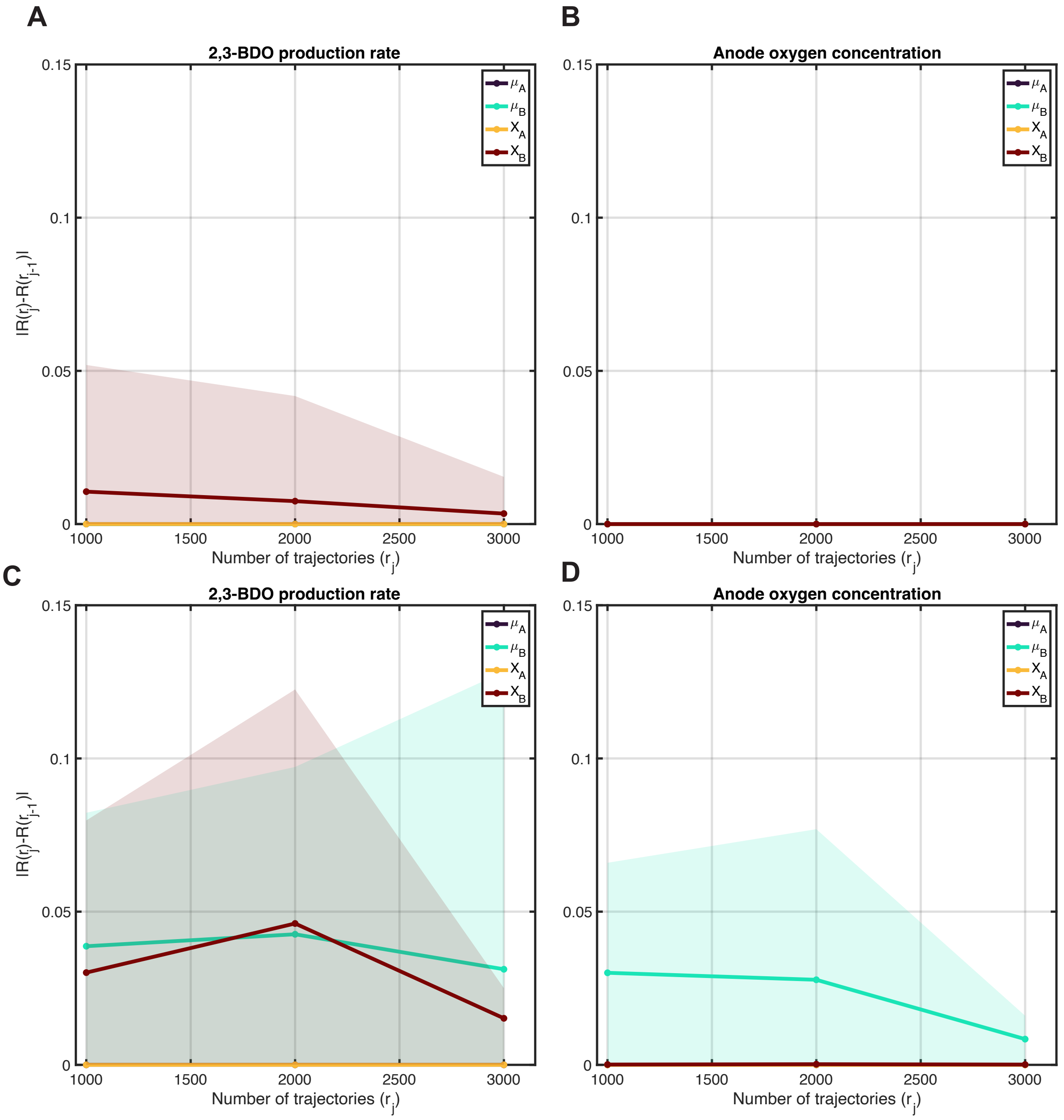


Figure 9: **Successive differences of the monotonicity index as the number of trajectories increase in a reprasentative 5x5 grid. A.,B.** Successive differences of the monotonicity index for all points with $h_1 > 10\ \mu m$ for the production rate and anodic oxygen concentration respectively, **C.,D.** Successive differences of the monotonicity index for only points with $h_1 = 10\ \mu m$ for the production rate and anodic oxygen concentration respectively.

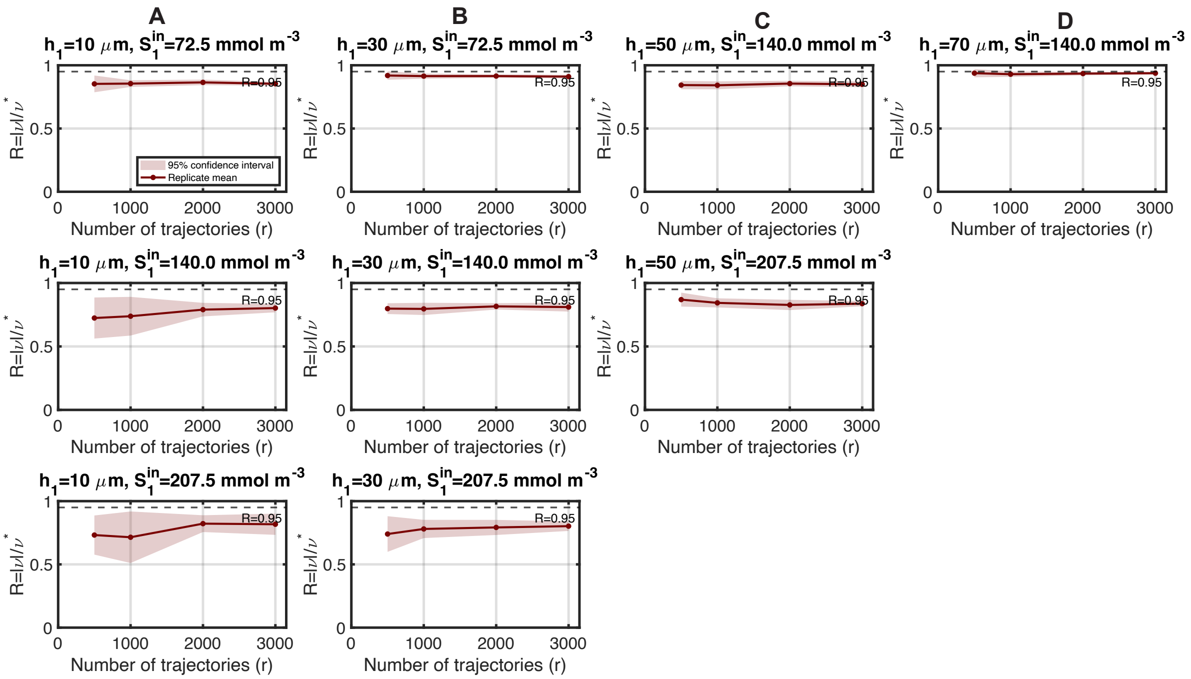


Figure 10: **Stabilization of monotonicity index at non-monotonic points (**$R < 0.95$**.** As the number of replicates increase, the confidence interval tightens around the mean. Points at **A.** $h_1 = 10\ \mu m$, **A.** $h_1 = 10\ \mu m$, **B.** $h_1 = 30\ \mu m$, **C.** $h_1 = 50\ \mu m$, **D.** $h_1 = 70\ \mu m$

### 5 Derivation of Reaction Rates

We derive the maximum reaction rates from the reported data established in (Novak et al., 2020), (Ricci et al., 2025),(Hwang et al., 2026). First we start with the aerobic reaction rate. We assume cell dry weight (CDW) is 100 % active biomass.

In Table 6 of (Novak et al., 2020), two specific reaction rates are listed from acetate only pulses, the first below representing diol production rate (acetoin and 2,3-butanediol) and the second representing acetate consumption rate:

$$q_{diols} = 0.016 \text{ g diols } (\text{g CDW})^{-1} \text{ h}^{-1}, \quad q_{ace} = 0.24 \text{ g acetate } (\text{g CDW})^{-1} \text{ h}^{-1}.$$

For our model we choose $q_{ace}$. While this significantly overestimates the diol production rate, it accurately captures acetate utilization which is central to analyzing reactor practicality allowing us to accurately estimate the minimum reaction rate necessary to surmount the tipping point. We need to convert to appropriate units. Adjusting to days and using molar mass we obtain:

$$\begin{aligned} q_{ace} &= 5.76 \text{ g ace } (\text{g CDW})^{-1} \text{ d}^{-1} \\ &= 5.76 \left( \frac{1/60 \text{ mol acetate}}{1/24.6 \text{ Cmol biomass}} d^{-1} \right) \\ &= 2.39 \left( \frac{\text{mol acetate}}{\text{Cmol biomass}} d^{-1} \right) \end{aligned}$$

Then using the stoichiometry, we can find the appropriate formulation for one mole of reaction per Cmol per day

$$= 2.39\left(\frac{\text{mol acetate}}{\text{Cmol biomass}}d^{-1}\right)\frac{1}{a}\frac{\text{mol reaction}}{\text{mol acetate}}$$
$$= 0.47\ \text{mol}_{\text{rxn}}\ (\text{Cmol biomass})^{-1}\text{d}^{-1}$$

So now to get the total reaction rate max, the respective stoichiometric value can be multiplied the maximum reaction rate. For the duration we write $\text{mol}_{\text{rxn}} = \text{mol reaction}$.

Recently, Ricci et al. (2025) improved upon the reaction rate $q_{ace}$ over 3 fold, $0.78\,\text{g acetate}\ (\text{g CDW})^{-1}\ \text{hour}^{-1}$. Using the same method as above we obtain a reaction rate

$$\mu_B = 1.53\ \text{mol}_{\text{rxn}}(\text{Cmol biomass})^{-1}\text{d}^{-1}.$$

Both reaction rates were found using fedbatch protocols, which we assume is the same for all protocols in this model.

Next, using (Hwang et al., 2026), they reported a maximum of $5.4\ \mu\text{mol acetate}\ (\text{mg protein})^{-1}$ over a 6 day culture. As we did for our baseline biofilm density values, converting to biomass we achieve approximately $0.022\ \mu\text{mol acetate}\ (\text{Cmol protein})^{-1}\ \text{day}^{-1}$. Then dividing by 3, as we are normalizing by acetate so we can be consistent with units we achieve

$$0.0074\ \text{mol}_{\text{rxn}}(\text{Cmol protein})^{-1}\ \text{day}^{-1}.$$

Since CDW is typically 50-55$\%$ active protein, and we assumed CDW equals 100$\%$ active biomass then (Rittmann and McCarty, 2020)

$$0.004\ \text{mol}_{\text{rxn}}(\text{Cmol biomass})^{-1}\ \text{day}^{-1}$$

Forward methanogenic pathway: from (Pinto et al., 2012) they report a maximum reaction rate of $14.12\ \text{mg acetate}\ (\text{mg biomass})^{-1}\text{d}^{-1}$ in the supplements. Converting this, we obtain $5.88\ \text{mol acetate}\ (\text{Cmol biomass})^{-1}\text{d}^{-1}$. Then dividing by 3 we achieve a forward rate of approximately $2\ \text{mol}_{\text{rxn}}\ (\text{Cmol biomass})^{-1}\text{d}^{-1}$.

therefore

$$\mu_A = 0.004\ \text{mol}_{\text{rxn}}(\text{Cmol biomass})^{-1}\ \text{day}^{-1}$$

and

$$\mu_{B,\text{Novak}} = 0.48\ \text{mol}_{\text{rxn}}(\text{Cmol biomass})^{-1}\text{d}^{-1}$$
$$\mu_{B,\text{Ricci}} = 1.53\ \text{mol}_{\text{rxn}}(\text{Cmol biomass})^{-1}\text{d}^{-1}$$

and

$$\mu_{A,F} = 2\ \text{mol}_{\text{rxn}}(\text{Cmol biomass})^{-1}\text{d}^{-1}.$$

### 6 Assumptions and limitations of the LPFR reduction

The bulk flow, concentration boundary layer (CBL), and biofilm regions are treated as separate spatial domains. The CBL has prescribed thickness $L_L$, allowing the mass transfer resistance to vary as an engineering design parameter. A smaller prescribed $L_L$ represents enhanced transverse transport associated with favorable reactor design. Independently, the reactive wall contribution is assigned an $O(\epsilon)$ ordering in the axial dispersion reduction. This weak wall assumption is introduced for analytical tractability, since retaining an $O(1)$ interfacial flux would require solving the fully coupled leading order transverse bulk–biofilm problem. The resulting model should therefore be interpreted as an optimistic, reduced-order approximation used for reactor comparison. While these are independent assumptions, there coupling through interfacial flux motivates such decisions. If the CBL was not treated as a separate region, its effective transverse transport reflecting CBL like behavior would be defined by solution to the transverse closure problem. This would not give us an explicit value for the CBL but rather is implicitly defined in the solutions behavior. By decoupling, we are given more engineering design control.

The effective height of the channel height is

$$H_{\text{Eff}} = H - h - L_L, \tag{7}$$

where $H$ is the total channel height and $h$ is the biofilm thickness. The Poiseuille velocity profile and cross-sectional averaging are applied only over $H_{\text{Eff}}$.

If a distinct asymptotically thin boundary layer existed, its effective transport could be derived by matching an inner near wall solution to the outer bulk flow solution via the method of matched asymptotics. However, for the present system, the reaction may effect the bulk behavior in the leading order $O(1)$ solution to reduced flow problem. Such an analysis is outside the scope of the present reactor comparison.

For analytical closure of the cross-sectionally averaged equation, the CBL flux is approximated by

$$J_{i,\text{CBL}} \approx \frac{D_{iW}}{L_L}\left(\bar{C}_i - S_{i,\text{BF}}\right). \tag{8}$$

In dimensionless form, the wall contribution scales as

$$\frac{H_{\text{Eff}}}{L_L}\left(\bar{\theta}_i - \theta_{i,\text{BF}}\right). \tag{9}$$

We assign

$$\frac{H_{\text{Eff}}}{L_L} = O(1), \qquad \bar{\theta}_i - \theta_{i,\text{BF}} = O(\epsilon), \tag{10}$$

such that the flux contribution is $O(\epsilon)$.

The wall sink contribution is assigned the weak sink ordering used in the first-order asymptotic expansion. Consequently, the resulting effective velocity

$$\bar{u}_{\text{Eff}} = \bar{u}\left(1 + \frac{k_L H_{\text{Eff}}}{60 D_{iW}}\right), \tag{11}$$

should be interpreted as a first-order reduced-order approximation, rather than an arbitrary strength reactive wall solution. However, we use the original substrate to govern all diffusion coefficients for the wall transfer, thus resulting in

$$\bar{u}_{\text{Eff}} = \bar{u}\left(1 + \frac{k_L H_{\text{Eff}}}{60 D_{2W}}\right), \tag{12}$$

The assumed value of $L_L$ is an effective engineering parameter and is not predicted by the model. In practice, it may vary with axial position, flow rate, reactor internals, species properties, and biofilm activity (Horn and Lackner, 2014; Leal, 2007). Accordingly, the LPFR results represent an optimistic performance estimate rather than a fully optimized reactor design.

Now we estimate the engineering benchmark CBL to contextualize our optimized reactor enhancements by use of the Sherwood number (Shah and London, 1978)

$$Sh = \frac{k_L D_{H_f}}{D_{iW}}, \tag{13}$$

where $D_{H_f}$ is the hydraulic diameter and where $k_m$ is the bulk mass transfer defined by equivalent film approximation

$$k_{L,i} = \frac{D_{iW}}{L_L}. \tag{14}$$

which together can provide a benchmark for the model CBL (Horn and Lackner, 2014; Leal, 2007)

$$L_L = \frac{D_{H_f}}{Sh}. \tag{15}$$

The boundary condition coupling the biofilm and the bulk is neither a prescribed Dirichlet concentration nor a prescribed Neumann flux but rather represents the coupled system. Therefore we use these limits as engineering benchmarks.

These scenarios are driven by different biofilm kinetics resulting in idealized boundary conditions at the bulk-biofilm interface: (i) under substrate saturated conditions in the biofilm, Monod kinetics approach zeroth order reaction rates in the biofilm effectively. The biofilm uptake rate is concentration independent approximating uniform concentration flux boundary condition at the bulk-biofilm interface. Where the other wall, i.e. the parallel plate to the biofilm has a no flux boundary condition, the fully developed parallel plate has a Sherwood number of (Shah and London, 1978)

$$Sh_{\text{flux}} = 5.385. \tag{16}$$

(ii)Under substrate limited conditions, the Monod rate approaches a first order reaction rate. If the effective biofilm uptake rate is also much faster than bulk concentration influx, the bulk-biofilm concentration approaches a fixed depleted value. In the limiting case, the interface behaves as a perfect absorber and is approximated by a homogeneous Dirichlet boundary condition The corresponding Sherwood number where the other wall has a no flux BC is (Shah and London, 1978)

$$Sh_{\text{Dir}} = 4.861. \tag{17}$$

With a channel height of $H_{\text{f}} = 4.95$ mm, a hydraulic diameter $D_{H_{\text{f}}} = 4A/P = 2\frac{WH_{\text{f}}}{W+H_{\text{f}}} \approx 2H_f = 9.9$ mm since $H_f$ is roughly 5% of W. The CBL is

$$\delta_{\text{flux}} = \frac{9.9}{5.385} \approx 1.84, \quad \delta_{\text{Dir}} = \frac{9.9}{4.861} \approx 2.04 \text{ mm}. \tag{18}$$

This provides an approximate fully developed flow CBL under idealized biofilm kinetics as an engineering benchmark. Thus, by applying an "enhancement factor" via reactor design with baffles or mixers, we can decrease the CBL. In the case shown in the paper, the boundary layer enhancement is approximately equal to 2 from our engineering benchmark $L_L = 1$ mm $= 1000$ $\mu$m.

#### 6.1 Flow Rates

Taylor Dispersion occurs when the transverse diffusion time is smaller than axial advection time, thus the flow rates must obey

$$Q \leq \frac{1}{M}\frac{D_{iW}LW}{H_{\text{Eff}}}. \tag{19}$$

where $M \geq 1$ is a safety factor quantifying how far into the valid Taylor regime a given flow rate falls.For our system $Q \approx 6.6 \times 10^{-4}$ m$^3$ day$^{-1}$ for $M = 1$.We choose the upper bound $Q_{\text{max}} = 10 \times 10^{-4}$ m$^3$ day$^{-1}$. Although it is slightly larger than the allowed region, it allows for the comparison in the DL-MEC bias region. The minimum however, $Q_{\text{min}} \approx 0.5 \times 10^{-4}$ m$^3$ day$^{-1}$ is firmly in the Taylor regime where $M > 10$.

For the HRT calculations, they correspond to local flow rates of the reactors rather allowing us to widely vary the input flow rates. For example, for a $T_{\mathrm{HRT}} \approx 1.9\ \mathrm{hours}$ for the DL-MEC corresponds to a flow rate of $Q_{\mathrm{DL-MEC,max}} \approx 126{\times}10^{-4}\ \mathrm{m}^3\ \mathrm{day}^{-1}$ while that same HRT corresponds to $Q_{\mathrm{LPFR,max}} = 10{\times}10^{-4}\ \mathrm{m}^3\ \mathrm{day}^{-1}$ This allows us to analyze see how productive these reactors are per residence time. The discrepancy is due to the respective volumes.

| Symbol | Description | Units |
|---|---|---|
| $S_1$ | Concentration of Oxygen | mol m$^{-3}$ |
| $S_2$ | Concentration of Methane | mol m$^{-3}$ |
| $S_3$ | Concentration of Acetate | mol m$^{-3}$ |
| $S_4$ | Concentration of 2,3-Butanediol | mol m$^{-3}$ |
| $S_5$ | Concentration of Bicarbonate | mol m$^{-3}$ |
| $M_{\mathrm{H}_2}(t)$ | Cumulative Hydrogen Produced | mol |
| $S_i^{\mathrm{in}}$ | Influent concentration of species $i$ | mol m$^{-3}$ |
| $S_i^{\max}$ | Maximum saturation concentration of species $i$ | mol m$^{-3}$ |
| $X_A$ | Biomass density of anaerobic biofilm (Biofilm A) | Cmol m$^{-3}$ |
| $X_B$ | Biomass density of aerobic biofilm (Biofilm B) | Cmol m$^{-3}$ |
| $h_1$ | Thickness of anaerobic biofilm (Biofilm A) | m |
| $h_2$ | Thickness of aerobic biofilm (Biofilm B) | m |
| $h$ | Total biofilm thickness ($h = h_1 + h_2$) | m |
| $L_L$ | Concentration boundary layer thickness | m |
| $\mu_A$ | Maximum reaction rate of anaerobic biofilm | mol$_{\mathrm{rxn}}$ Cmol$^{-1}$ day$^{-1}$ |
| $\mu_B$ | Maximum reaction rate of aerobic biofilm | mol$_{\mathrm{rxn}}$ Cmol$^{-1}$ day$^{-1}$ |
| $K_{1A}$ | Oxygen inhibition half-saturation constant (Biofilm A) | mol m$^{-3}$ |
| $K_{1B}$ | Oxygen half-saturation constant (Biofilm B) | mol m$^{-3}$ |
| $K_{2A}$ | Methane half-saturation constant (Biofilm A) | mol m$^{-3}$ |
| $K_{3B}$ | Acetate half-saturation constant (Biofilm B) | mol m$^{-3}$ |
| $K_{5A}$ | Bicarbonate half-saturation constant (Biofilm A) | mol m$^{-3}$ |
| $D_{iW}$ | Diffusion coefficient of species $i$ in water | m$^2$ day$^{-1}$ |
| $D_{ij}$ | Effective diffusion coefficient of species $i$ in biofilm layer $j$ | m$^2$ day$^{-1}$ |
| $\epsilon_j$ | Porosity of biofilm layer $j$ | – |
| $Q$ | Volumetric flow rate | m$^3$ day$^{-1}$ |
| $V_W$ | Anolyte (water) volume | m$^3$ |
| $A$ | Biofilm/electrode surface area | m$^2$ |
| $T_{\mathrm{HRT}}$ | Hydraulic retention time | h |
| $RR_i$ | Removal rate of species $i$ | $\mu$mol day$^{-1}$ |
| $PR_4$ | Production rate of 2,3-BDO | $\mu$mol day$^{-1}$ |
| $K_La$ | Volumetric mass transfer coefficient | h$^{-1}$ |
| $i$ | Current density | A m$^{-2}$ |
| $E_{\mathrm{CEF}}^{0'}$ | Standard counter electromotive force | V |
| $\eta_{\mathrm{ohm}}$ | Ohmic overpotential | V |
| $\sigma_{\mathrm{BF}}$ | Biofilm conductivity | S m$^{-1}$ |
| $H_{\mathrm{ES}}$ | Electrode separation distance | m |
| $N$ | Number of TIS tanks | – |
| $N_{\mathrm{Tot}}$ | Number of Sobol points | – |
| $N_S$ | Number of Sobol points surpassing threshold | – |
| $R_p^{ij}$ | Monotonicity index | – |
| $\lambda_{ij}$ | Distance metric (normalized Euclidean) | – |
| $P_1$ | Total production metric (equal flow rate) | $\mu$mol day$^{-1}$ |
| $P_2$ | Volumetric production metric (equal HRT) | $\mu$mol m$^{-3}$ day$^{-1}$ |
| $P_3$ | Areal production metric (equal HRT) | $\mu$mol m$^{-2}$ day$^{-1}$ |
| $\bar{u}$ | Mean velocity in LPFR | m day$^{-1}$ |
| $D_{\mathrm{ax}}$ | Axial dispersion coefficient | m$^2$ day$^{-1}$ |
| $D_{\mathrm{disp}}$ | Taylor dispersion coefficient | m$^2$ day$^{-1}$ |
| $k_L$ | Film mass transfer coefficient (LPFR) | m day$^{-1}$ |

Table 3: **Model variables and parameters**.

### References


[1] Dylan G Boucher, Emily Carroll, Zachary A Nguyen, Rohit G Jadhav, Olja Simoska, Kevin Beaver, and Shelley D Minteer. Bioelectrocatalytic synthesis: concepts and applications. *Angewandte Chemie International Edition*, 62(46):e202307780, 2023.

[2] Thomas K. Wood, Ilke Gurgan, Ethan T. Howley, and Ingmar H. Riedel-Kruse. Converting methane into electricity and higher-value chemicals at scale via anaerobic microbial fuel cells. *Renewable and Sustainable Energy Reviews*, 188:113749, 2023. ISSN 1364-0321. doi: https://doi.org/10.1016/j.rser.2023.113749. URL https://www.sciencedirect.com/science/article/pii/S1364032123006068.

[3] Cristian Picioreanu, Mark CM van Loosdrecht, Thomas P Curtis, and Keith Scott. Model based evaluation of the effect of ph and electrode geometry on microbial fuel cell performance. *Bioelectrochemistry*, 78 (1):8–24, 2010.

[4] Cristian Picioreanu, Ian M Head, Krishna P Katuri, Mark CM van Loosdrecht, and Keith Scott. A computational model for biofilm-based microbial fuel cells. *Water research*, 41(13):2921–2940, 2007.

[5] André Baudler, Igor Schmidt, Markus Langner, Andreas Greiner, and Uwe Schröder. Does it have to be carbon? metal anodes in microbial fuel cells and related bioelectrochemical systems. *Energy Environ. Sci.*, 8:2048–2055, 2015. doi: 10.1039/C5EE00866B. URL http://dx.doi.org/10.1039/C5EE00866B.

[6] T. E. Springer, T. A. Zawodzinski, and S. Gottesfeld. Polymer electrolyte fuel cell model. *Journal of The Electrochemical Society*, 138(8):2334, aug 1991. doi: 10.1149/1.2085971. URL https://doi.org/10.1149/1.2085971.

[7] Thomas A. Zawodzinski, C. R. Derouin, Susan D. Radzinski, Ruth J. Sherman, Van T. Smith, Thomas E. Springer, and Shimshon Gottesfeld. Water uptake by and transport through nafion® 117 membranes. *Journal of The Electrochemical Society*, 140:1041–1047, 1993. URL https://api.semanticscholar.org/CorpusID:93471671.

[8] Vaibhav Kumar Dixit and Christopher Rackauckas. Globalsensitivity. jl: Performant and parallel global sensitivity analysis with julia. *Journal of Open Source Software*, 7(76):4561, 2022.

[9] Max D Morris. Factorial sampling plans for preliminary computational experiments. *Technometrics*, 33 (2):161–174, 1991.

[10] Katharina Novak, Regina Kutscha, and Stefan Pflügl. Microbial upgrading of acetate into 2, 3-butanediol and acetoin by e. coli w. *Biotechnology for biofuels*, 13:1–14, 2020.

[11] Luca Ricci, Xuecong Cen, Yuexuan Zu, Giacomo Antonicelli, Zhen Chen, Debora Fino, Fabrizio C Pirri, Gregory Stephanopoulos, Benjamin M Woolston, and Angela Re. Metabolic engineering of e. coli for enhanced diols production from acetate. *ACS Synthetic Biology*, 14(4):1204–1219, 2025.

[12] Hyeon-Ji Hwang, Mrugesh Parasa, Ruchira Mitra, Rodolfo García-Contreras, Vanesa Angarita-Zapata, Ingmar H. Riedel-Kruse, and Thomas K. Wood. Engineering methyl-coenzyme m reductase for enhanced methane and carbon dioxide capture through biofilm growth. *bioRxiv*, 2026. doi: 10.64898/2026.06.26.734734. URL https://www.biorxiv.org/content/early/2026/06/27/2026.06.26.734734.

[13] B.E. Rittmann and P.L. McCarty. *Environmental Biotechnology: Principles and Applications, Second Edition*. McGraw Hill LLC, 2020. ISBN 9781260441611. URL https://books.google.com/books?id=VXLwDwAAQBAJ.

[14] R.P. Pinto, B. Tartakovsky, and B. Srinivasan. Optimizing energy productivity of microbial electrochemical cells. *Journal of Process Control*, 22(6):1079–1086, 2012. ISSN 0959-1524. doi: https://doi.org/10.1016/j.jprocont.2012.04.005. URL https://www.sciencedirect.com/science/article/pii/S095915241200090X.

[15] Harald Horn and Susanne Lackner. Modeling of biofilm systems: a review. *Productive biofilms*, pages 53–76, 2014.

[16] L. Gary Leal. *Advanced Transport Phenomena: Fluid Mechanics and Convective Transport Processes*. Cambridge Series in Chemical Engineering. Cambridge University Press, Cambridge, 2007. ISBN 9780521849104. doi: DOI:10.1017/CBO9780511800245. URL https://www.cambridge.org/product/9F2A633638780413DA73C6CB70A1D341.

[17] R.K. Shah and A.L. London. Chapter v - circular duct. In *Laminar Flow Forced Convection in Ducts*, pages 78–152. Academic Press, 1978. ISBN 978-0-12-020051-1. doi: https://doi.org/10.1016/B978-0-12-020051-1.50010-3. URL https://www.sciencedirect.com/science/article/pii/B9780120200511500103.
